# Active mode of excretion across digestive tissues predates the origin of excretory organs

**DOI:** 10.1101/136788

**Authors:** Carmen Andrikou, Daniel Thiel, Juan A. Ruiz-Santiesteban, Andreas Hejnol

**Affiliations:** Sars International Centre for Marine Molecular Biology, University of Bergen, Thormøhlensgate 55, 5006 Bergen, Norway

## Abstract

Most bilaterian animals excrete toxic metabolites through specialized organs, such as nephridia and kidneys, which share morphological and functional correspondences. In contrast, the excretory mechanisms in non-nephrozoans are largely unknown, and therefore the reconstruction of ancestral excretory mechanisms is problematic. Here, we investigated the excretory mode of members of the Xenacoelomorpha, the sister group to Nephrozoa, and Cnidaria, the sister group to Bilateria. By combining gene expression, inhibitor experiments and exposure to varying environmental ammonia conditions we show that both, Xenacoelomorpha and Cnidaria, are able to excrete across digestive-associated tissues. Based on these results we propose that digestive-associated tissues functioned as excretory sites before the evolution of specialized organs in nephrozoans. We conclude that diffusion was likely the ancestral mode of excretion, whilst the emergence of a compact, multiple-layered bilaterian body plan necessitated the evolution of active transport excretory mechanisms that was later recruited into the specialized excretory organs.

## Introduction

Excretory organs are specialized organs that remove toxic metabolic waste products and control water and ion balance in animals based on the principles of ultrafiltration, active transport and passive transport/diffusion^1^. They are only present in Nephrozoa (Deuterostomia + Protostomia)^2^ (Fig. 1a) and based on morphological correspondences can be grouped into two major architectural units: the protonephridia, only found in Protostomia, and the metanephridia, present in both Deuterostomia and Protostomia^3,4^. Both organs are organized into functionally similar compartments: the terminal cells of protonephridia and the podocytes associated to metanephridial systems conduct ultrafiltration, and the tubule and duct cells modify the filtrate through a series of selective reabsorption and secretion, via passive and active transport mechanisms^5^ (Fig. 1b). There exist also other, taxon-specific excretory organs and excretory sites, which perform either ultrafiltration (such as the nephrocytes of insects^6^ and the rhogocytes of gastropods^7^), or absorption and secretion (such as the malpighian tubules of various tardigrades, arachnids and insects^8^, the excretory cells of nematodes^9^, the gills of fish, shore crabs and annelids^10,11^, and the epidermis of planarians^12^). Molecular studies have shown that a suite of orthologous genes is involved in the excretory mechanisms of different nephrozoan species, regardless whether they possess specialized excretory organs or not^9-30^ (see also Supplementary Table 1). The passive ammonia transporters Rhesus/AMTs, the active transporter Na^+^/K^+^[NH_4_^+^] ATPase (NKA), the hyperpolarization-activated cyclic nucleotide-gated K^+^[NH_4_^+^] channels (HCN), the vacuolar H^+^-ATPase (v-ATPase subunits A and B), members of the Carbonic Anhydrase A (CA) and the water/glycerol/ammonia channels (aquaporins) are commonly used for excreting ammonia, the most toxic metabolite (Fig. 1c) (summarized in^5,31^). Also, a group of orthologous slit diaphragm structural components (nephrin/kirre, cd2ap, zo1, stomatin/podocin), whose function is associated with the maintenance of the ultrafiltration apparatus by interacting with the actin cytoskeleton and forming tight junctions^32^, is localized at the ultrafiltration site of the podocytes of the rodent kidney^33^ (Fig. 1b) as well as in the *Drosophila* nephrocytes^34^ and the rhogocytes of gastropods^7^. Finally, a number of ion transporters (solute carrier transporters (SLCs)) are spatially expressed in the corresponding compartments of protonephridia of planarians and metanephridia (e.g. kidneys) of vertebrates^35-37^ (Fig. 1b).

**Figure 1.**
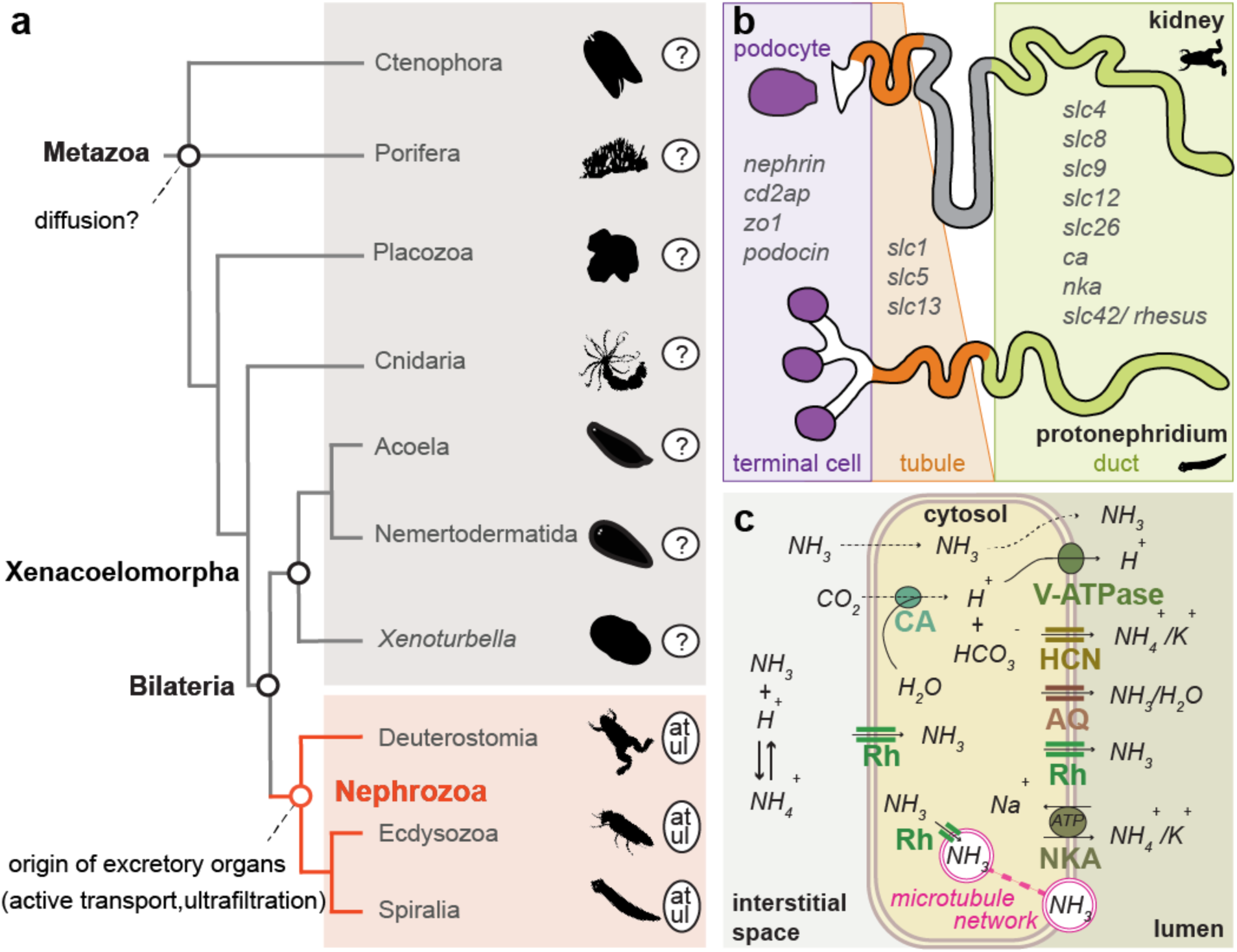
Traditional diffusion hypothesis, ammonia transport mechanism and structural and functional correspondences of protonephridial and metanephridial systems. (**a**) Illustrated phylogenetic relationship between Nephrozoa, Xenacoelomorpha and non-bilaterians^47^. Excretory organs or specialized excretory cells/tissues using active transport (at) and ultrafiltration (ul) are so far only reported in the group of Nephrozoa. (**b**) Cartoon depiction of the structural components of metanephridia (podocyte, duct, tubule) in comparison to protonephridia (terminal cell, duct, tubule) and summary of the expression domains of orthologous selected genes in relation to their components. Animal illustrations are taken from phylopic.org (CC BY 3.0). (**c**) Ammonia cellular transport. Ammonia (NH_3_) is secreted into the lumen fluid via parallel H^+^ and ammonia transport. This involves passive diffusion through the cell membrane (dashed lines), facilitated diffusion via the Rhesus glycoproteins (Rh), active transport via the Na^+^/K^+^[NH4^+^] ATPase (NKA), the hyperpolarization-activated cyclic nucleotidegated K^+^[NH4^+^] channel (HCN) and aquaporin transporters (AQ) as well as the generation of H^+^ gradient by a vacuolar H^+^-ATPase proton pump (v-ATPase) and the carbonic anhydrase (CA), which transforms CO_2_ into H^+^ and HCO_3_^−^. Vesicular ammonia-trapping mechanism is also illustrated.

The excretory sites and mechanisms in non-nephrozoans, however, are largely unknown. It is commonly stated that excretion is presumably occurring via diffusion across the body wall due to the loose (e.g. sponges) or single-epithelial (cnidarians and ctenophores) cellular organization of these animals^1,38,39^ (Fig. 1a) (herein stated as “diffusion hypothesis”). Based on this idea, it was hypothesized that the emergence of the first excretory organs coincided with the evolution of multilayered, solid parenchymes and increased body sizes due to the need of more elaborate excretory mechanisms^40,41^. However, since excretion in non-nephrozoans was never investigated in detail, the ancestral mechanisms of excretion and the evolutionary origin of excretory organs remain unresolved^1,40-45^.

An important animal group for our understanding nephrozoan evolution is their bilaterian sister group^2,46,47^, the Xenacoelomorpha (*Xenoturbella* + (Nemertodermatida + Acoela)). These small, worm-like animals exhibit a bilateral symmetric, multilayered body plan, but except for a special cell type with a putative excretory function (dermonephridia)^48^ that seems unique to the acoel *Paratomella*, xenacoelomorphs lack excretory organs and no defined excretory sites have yet been described. To understand the excretory mechanisms outside Nephrozoa and to gain insights into ancient excretory mechanisms, we therefore investigated the excretory mode in two xenacoelomorph species and compared our findings with the non-bilaterian cnidarian *Nematostella vectensis*.

## Results

### Most genes involved in excretion in Nephrozoa are already present in non-nephrozoan and non-bilaterian animals

To get an overview of the presence of excretion-related genes in xenacoelomorphs and non-bilaterian animals we first searched for orthologous sequences of 20 nephrozoan candidate genes in the available transcriptomes and draft genomes of 13 xenacoelomorph species as well as in representatives of cnidarians, placozoans and sponges (Supplementary Fig. 1a and 2). We found that most of these genes were present in almost all groups with the exceptions of *slc5* (a sodium glucose cotransporter) that was only present in Cnidaria and Bilateria, and the ultrafiltration component *nephrin*/*kirre* that was only present in Bilateria (Supplementary Table 1 and Supplementary Fig. 1a). This analysis also revealed that the last common ancestor of placozoans, cnidarians and bilaterians already had at least two paralogs of *amts* (*amt2/3* and *am1/4*) and one *rhesus*, with independent duplications of one or both of these in various animal lineages (Supplementary Fig. 2). To identify potential excretory sites in xenacoelomorphs we examined the expression of the entire set of these candidate excretion-related genes in the acoel *Isodiametra pulchra* and the nemertodermatid *Meara stichopi*^49^, which differ in their digestive system composition (*I. pulchra* has a syncytial, lumen-less gut, whereas *M. stichopi* has an epithelia-lined, cellular gut^50^) (Supplementary Fig. 1b).

### Genes encoding solute carrier transporters and slit diaphragm-related components are expressed broadly in *Isodiametra pulchra* and *Meara stichopi*

Genes encoding SLCs were broadly expressed within neural (brain and nerve cords), parenchymal/subepidermal, digestive and gonadal-associated cells in both animals (Fig. 2 and 3). In particular, the three *slc1* paralogs isolated from *I. pulchra* were expressed in the brain (Fig. 2A1, A3) and around the male gonopore (Fig. 2A2, A3) and the single copy found in *M. stichopi* was restricted to scattered cells, resembling neurons (Fig. 3A). The two *slc5* paralogs found in *I. pulchra* had diverse expression patterns, with one copy being expressed around the male gonopore (Fig. 2B1) and the other one labeling brain and parenchymal cells (Fig. 2B2), whilst the *slc5* copy of *M. stichopi* was expressed in scattered subepidermal cells and two posterior lateral rows of cells (Fig. 3B). The single *slc13* copy of *I. pulchra* was marking brain and parenchymal cells (Fig. 2C) and in the same fashion all four paralogs of *M. stichopi* were expressed in scattered cells, resembling neurons (Fig. 3C1-C3). The three *slc4* paralogs isolated from *I. pulchra* were labeling individual cells at the anterior tip (Fig. 2D1, D3), brain (Fig. 2D2) and parenchymal cells (Fig. 2D2-D3), whilst in *M. stichopi* the three *slc4* paralogs were broadly expressed including gut-wrapping cells and the mouth (Fig. 3D1), scattered subepidermal cells (Fig. 3D2-D3), gonads (Fig. 3D3) and nerve cords (Fig. 3D2 and Supplementary Fig. 3E). *Slc8* was restricted in a cup-shape region of the anterior tip in *I. pulchra* (Fig. 2E) and in *M. stichopi* was expressed in the gut epithelium (including the mouth) and two posterior lateral rows of cells (Fig. 3E). *Slc9* was marking the mouth in *I. pulchra* (Fig. 2F) and in *M. stichopi* was expressed in two proximal lateral rows of cells, resembling neurite bundles (Fig. 3F). Both *slc12* paralogs found in *I. pulchra* were labeling neural (brain) and parenchymal cells (Fig. 2G1-G2); whilst in *M. stichopi* the expression of the two paralogs investigated was associated with the male and female gonad region (Fig. 3G1-G2). One *slc26* copy did not reveal any expression in *I. pulchra* (Fig. 2H) although in *M. stichopi* the two copies were expressed differently, with one gene being restricted in the female gonads (Fig. 3H1) and the second one to be labeling gut-wrapping cells (Fig. 3H2).

**Figure 2.**
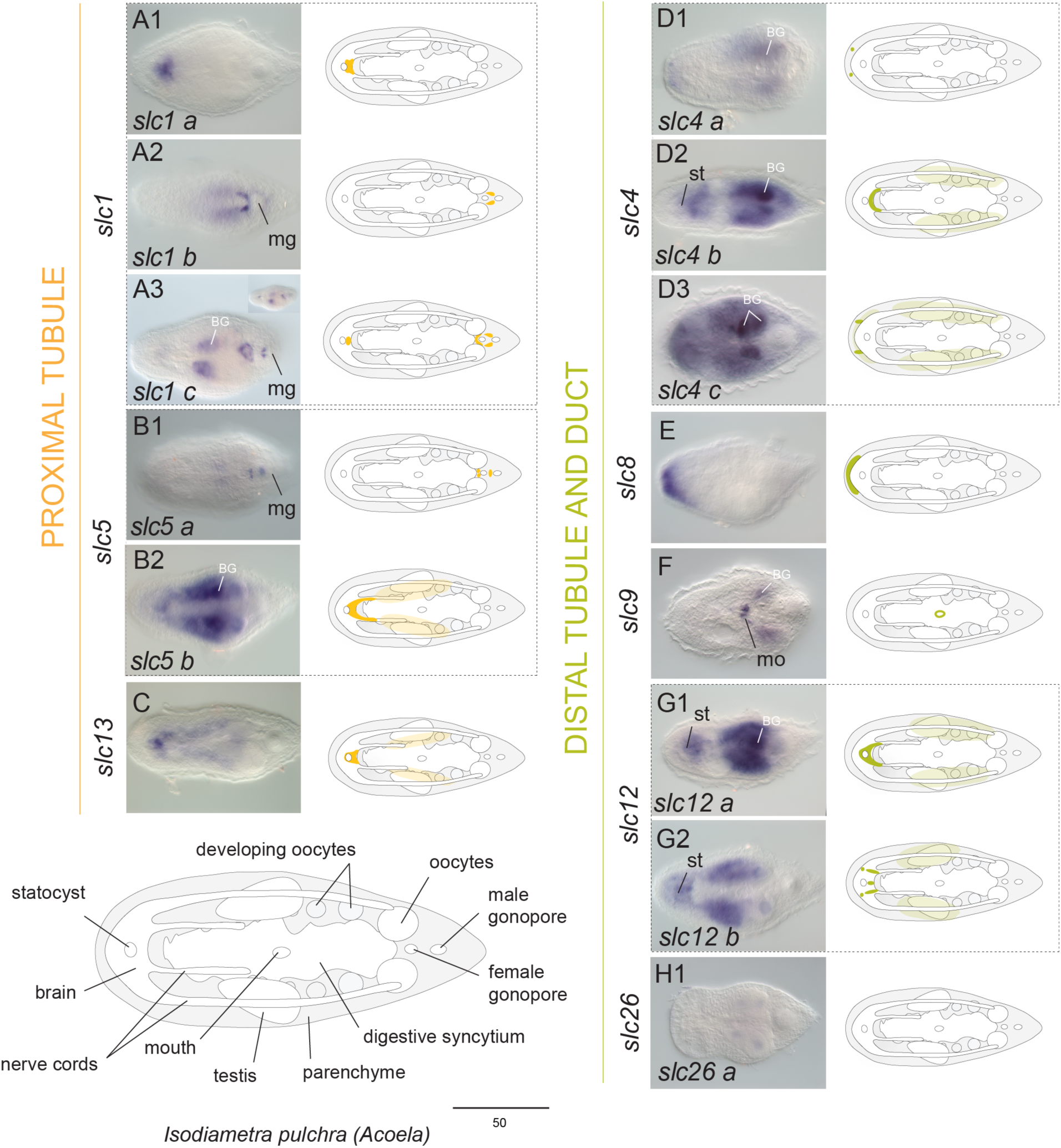
Expression of solute carrier transporters related to excrete modification in *I. pulchra.* Whole mount in situ hybridization (WMISH) of genes encoding *slc1, slc5* and *slc13* (proximal tubule related), *slc4, slc8, slc9, slc12* and *slc26* (distal tubule and duct related). The inset in panel A3 shows a different focal plane. Illustrations with colored gene expression correspond to the animals shown in the previous column. Anterior is to the left. The depicted expression patterns are for guidance and not necessarily represent exact expression domains. Drawings are not to scale. BG indicates background staining. Abbreviations: mo, mouth; mg, male gonopore; st, statocyst.

**Figure 3.**
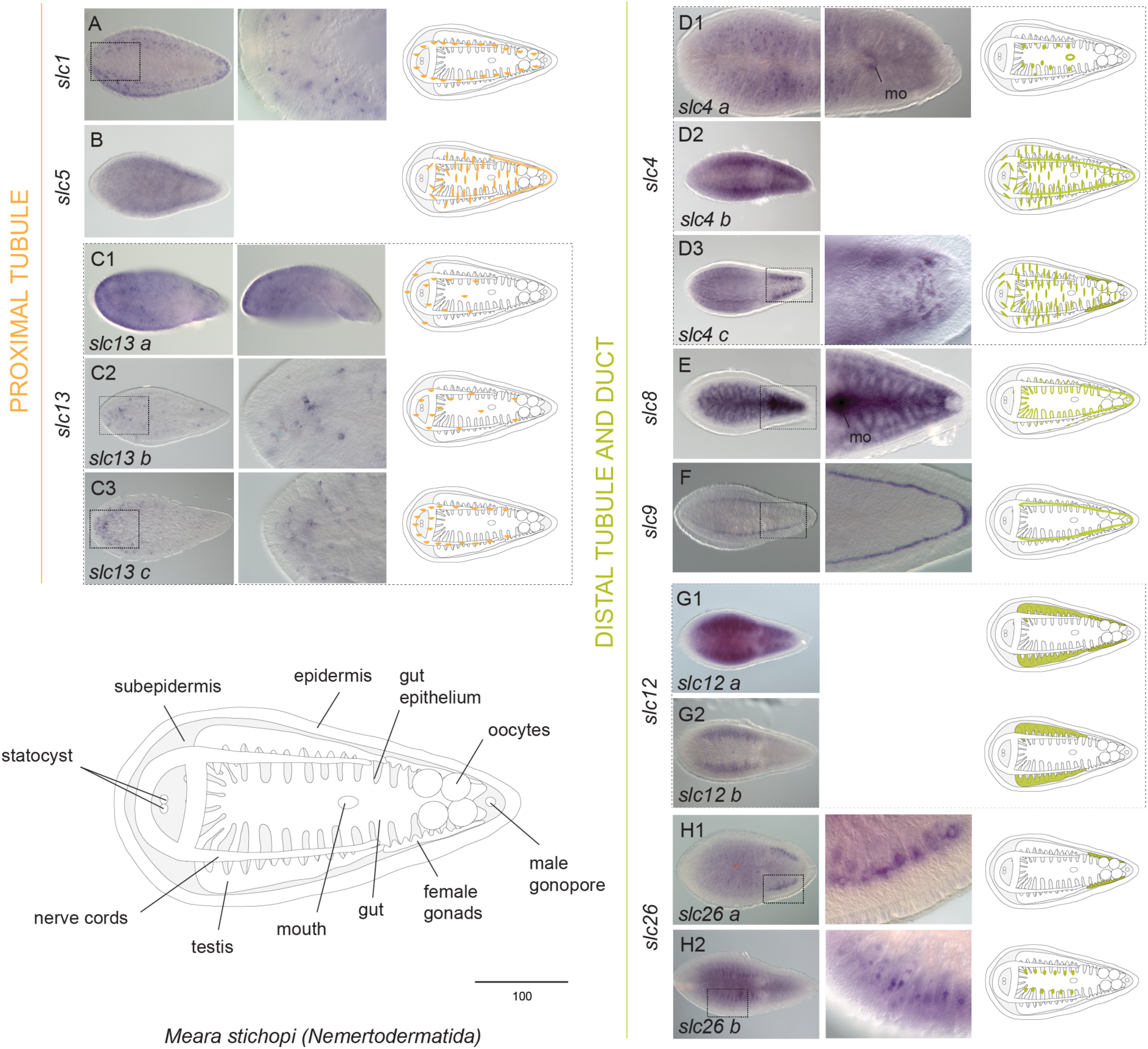
Expression of solute carrier transporters related to excrete modification in *M. stichopi.* Whole mount in situ hybridization (WMISH) of genes encoding *slc1, slc5* and *slc13* (proximal tubule related), *slc4, slc8, slc9, slc12* and *slc26* (distal tubule and duct related). The columns next to the panels show higher magnifications of the indicated domains, except of C1 that shows a different view form the animal (side view). Illustrations with colored gene expression correspond to the animals shown in the previous column. Anterior is to the left. The depicted expression patterns are for guidance and not necessarily represent exact expression domains. Drawings are not to scale. Abbreviations: mo, mouth.

Genes related to ultrafiltration sites had also broad expression patterns in both animals covering neural, parenchymal/subepidermal cells as well as digestive and gonadal-associated cells (Supplementary Fig. 4). All three *nephrin/kirre* paralogs were labeling neural cells both in *I. pulchra* and *M. stichopi* (Supplementary Fig.4A). *Nephrin/kirre 1* was expressed in the brain and around the male gonopore in *I. pulchra* (Supplementary Fig. 4A1) whilst *nephrin/kirre 2* and *3* were expressed in scattered cells, resembling neurons (Supplementary Fig. 4A2-3). In *M. stichopi*, all three paralogs were also expressed in proximal lateral rows of cells, resembling neurite bundles (Supplementary Fig. 4A1’-3’). *Cd2ap* was expressed in few cells posterior to the statocyst and in scattered parenchymal cells, resembling neural cells in *I. pulchra* (Supplementary Fig. 4B), whilst in *M. stichopi* this gene had a broad expression pattern including subepidermal cells, the mouth and two posterior lateral rows of cells (Supplementary Fig. 4B’). *Zo1* was marking anterior cells (probably the brain), the mouth and scattered cells of the parenchyme, resembling neurons in *I. pulchra* (Supplementary Fig. 4C), while in *M. stichopi* was restricted in gut-affiliated cells and the mouth (Supplementary Fig. 4C’). Finally, from the three *stomatin/podocin* paralogs found in *I. pulchra*, two were expressed in the brain (Supplementary Fig. 4D1, D3) and the third one in cells of the digestive syncytium (Supplementary Fig. 4D2) and in *M. stichopi* the five paralogs were mainly expressed in proximal lateral rows of cells, resembling neural bundles (Supplementary Fig. 4D1’, D5’), scattered subepidermal cells (Supplementary Fig. 4D3’-D4’) and the mouth (Supplementary Fig. 4D2’).

The broad expression of solute carrier transporters and slit diaphragm-related components in acoelomorphs suggests that they are not part of defined excretory domains, as in nephrozoans.

### The expression of a number of ammonia excretion-related genes and aquaporins suggests digestive-associated domains as putative excretion sites in *Isodiametra pulchra* and *Meara stichopi*

Genes involved in ammonia excretion (*rhesus/amts, nka, v-ATPase B, ca, hcn*) were mainly demarcating neural, digestive-associated and other parenchymal/subepidermal cells, as well as epidermal cells (Fig. 4 and 5). In particular, the ammonia transporter *rhesus* was expressed in cells of the anterior tip, gut-affiliated cells and the posterior ventral epidermis both in *I. pulchra* and *M. stichopi* (Fig. 4A, 5A). The six *amt* paralogs found in *I. pulchra* were broadly expressed including individual cells of the anterior tip (Fig. 4B5), scattered parenchymal cells resembling neurons (Fig. 4B4), the brain (Fig. 4B1, B5), a cup-shape region of the anterior tip (Fig. 4B1, B2), the mouth (Fig. 4B1), parenchymal and gut-affiliated cells (Fig. 4B3, B5), whist the only *amt* present in *M. stichopi* was restricted to the whole epidermis (Fig. 5B). Both *nka* paralogs of *I. pulchra* were strongly expressed in a sac-shaped gut-wrapping domain (Fig. 4C). To check whether the expression was extending to the male gonadal domain double FISH with markers of the gonads (*piwi* and *vasa*) were conducted and showed that the expression of *nka* is restricted in a defined gut-wrapping domain, adjacent to the male gonads (testes) (Supplementary Fig. 3B). Similarly, the single *nka* copy of *M. stichopi* was labeling gut-affiliated cells (Fig. 5C). Nine genes belonging to the family of alpha-Carbonic Anhydrases were found in *I. pulchra* and six in *M. stichopi*. *Ca a* was expressed in few distal cells at the anterior tip, cells at the posterior tip adjacent to the male gonopore, two rows of proximal parenchymal cells, resembling gut wrapping cells, and the mouth (Fig. 4D1), *ca b, ca c* and *ca h* were all expressed in scattered parenchymal cells resembling neurons (Fig. 4D2-D3, D5), *ca x* was labeling the brain (Fig. 4D6), while *ca d* was expressed in a cup-shape region of the anterior tip and parenchymal cells in *I. pulchra* (Fig. 4D4). In *M. stichopi ca a* and *ca c* were marking gut-affiliated (Fig. 5D1, D3) and *ca b* was expressed in gut-wrapping cells (Fig. 5D2). *Ca d* was expressed in scattered cells resembling neurons and cells of the posterior epidermis (Fig. 5D4). *Hcn* was expressed in the brain in *I. pulchra* (Fig. 4E and Supplementary Fig. 3C), whilst in *M. stichopi* was labeling gut-affiliated cells (Fig. 5E). *V-ATPase B* was strongly expressed in the digestive syncytium in *I. pulchra* (Fig. 4F and Supplementary Fig. 3A) and in gut-affiliated cells in *M. stichopi* (Fig. 5F and Supplementary Fig. 3D). In *M. stichopi v-ATPase B1* was additionally expressed in two posterior proximal lateral rows of cells, resembling neurite bundles (Fig. 5F2) and *v-ATPase B2* in scattered subepidermal cells (Fig. 5F1).

**Figure 4.**
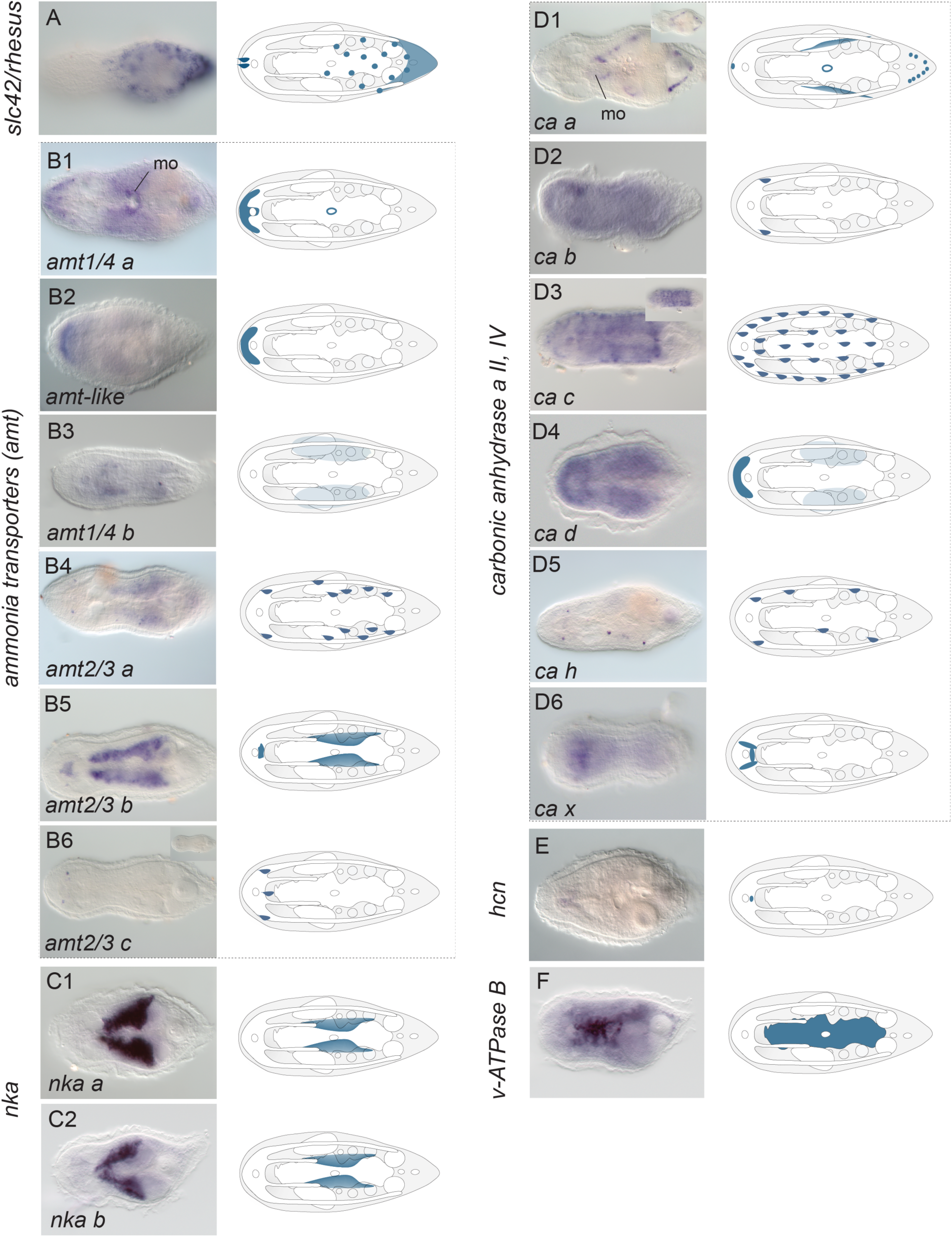
Expression of ammonia excretion-related genes in *I. pulchra.* Whole mount in situ hybridization (WMISH) of genes encoding rh, v-ATPase B, nka, ca A 2 & 4, hcn and amts. The inset in panel B6 and D1 show different focal planes. Illustrations with colored gene expression correspond to the animals shown in the previous column. Anterior is to the left. The depicted expression patterns are for guidance and not necessarily represent exact expression domains. Drawings are not to scale. Abbreviations: mo, mouth.

**Figure 5.**
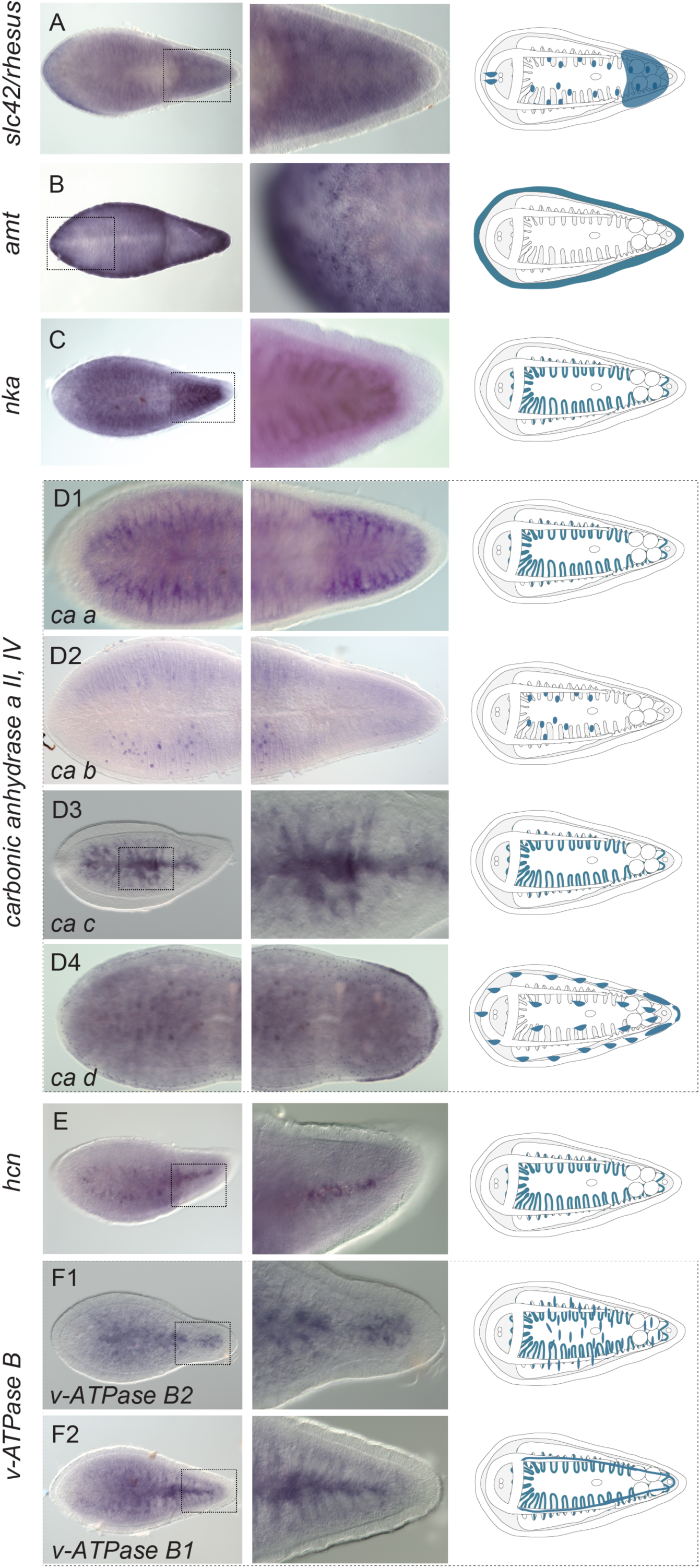
Expression of ammonia excretion-related genes in *M. stichopi.* Whole mount in situ hybridization (WMISH) of genes encoding rh, v-ATPase B, nka, ca A 2 & 4, hcn and amts. The columns next to the panels show higher magnifications of the indicated domains. Illustrations with colored gene expression correspond to the animals shown in the previous column. Anterior is to the left. The depicted expression patterns are for guidance and not necessarily represent exact expression domains. Drawings are not to scale.

The majority of genes encoding *aquaporins* were labeling digestive-associated domains and neural domains. From the seven paralogs found in *I. pulchra, aq c, aq d, aq f* and *aq g* were expressed in gut-wrapping cells (Fig. 6C-F) and *aq b* was labeling the digestive syncytium (Fig. 6B). *Aq g* was additionally expressed in individual cells lining the gonads, whilst *aq a* was expressed in the brain (Fig. 6A) and *aq e* was labeling parenchymal cells (Fig. 6G). In *M. stichopi* six paralogs were found. *Aq d* and *aq f* were marking gut-wrapping cells (Fig. 7D, F), while *aq a* and *aq c* were labeling proximal lateral rows of cells, resembling neurite bundles (Fig. 7A, C). Finally, *aq b* was localized in individual subepidermal cells and *aq e* was expressed in cells of the anterior tip and the ventral epidermis (Fig. 7E).

**Figure 6.**
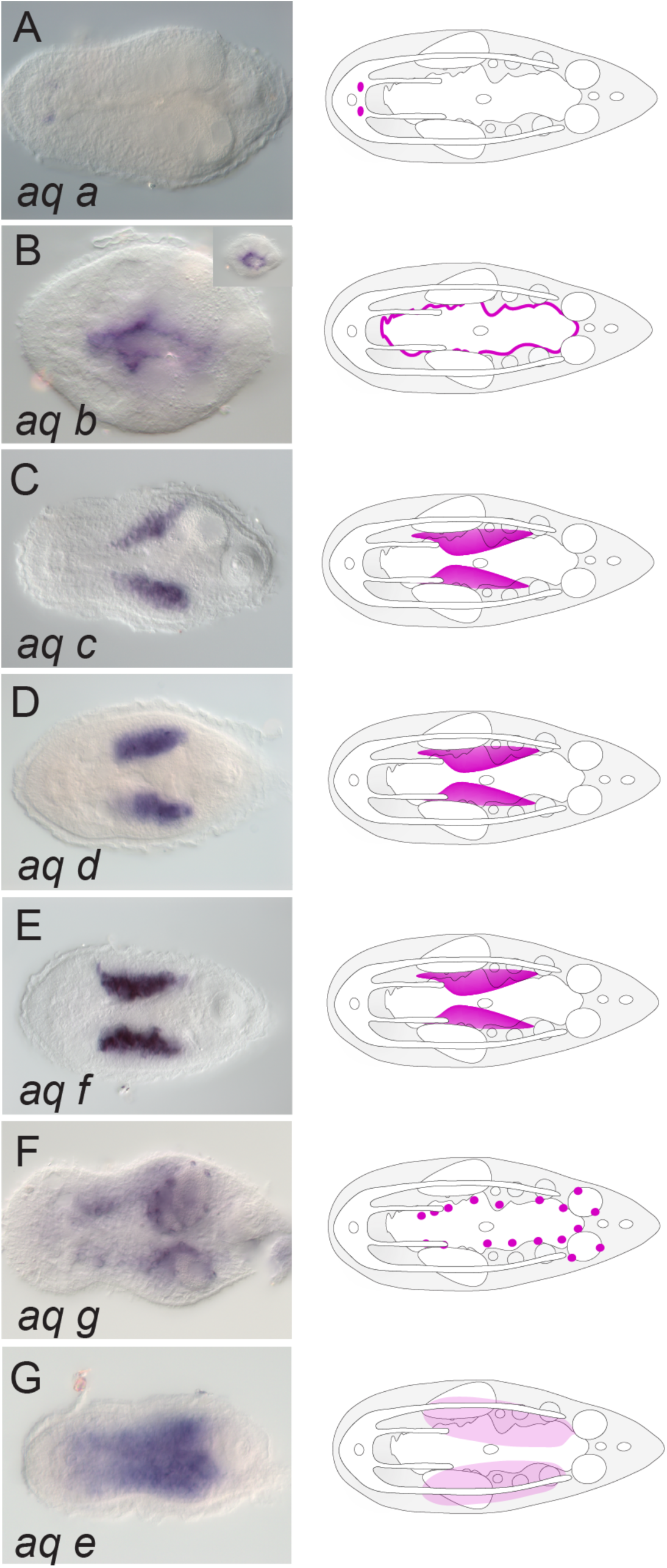
Expression of aquaporins in *I. pulchra.* Whole mount in situ hybridization (WMISH) of aquaporin genes. The inset in panel B shows a different focal plane. Illustrations with colored gene expression correspond to the animals shown in the previous column. Anterior is to the left. The depicted expression patterns are for guidance and not necessarily represent exact expression domains. Drawings are not to scale.

**Figure 7.**
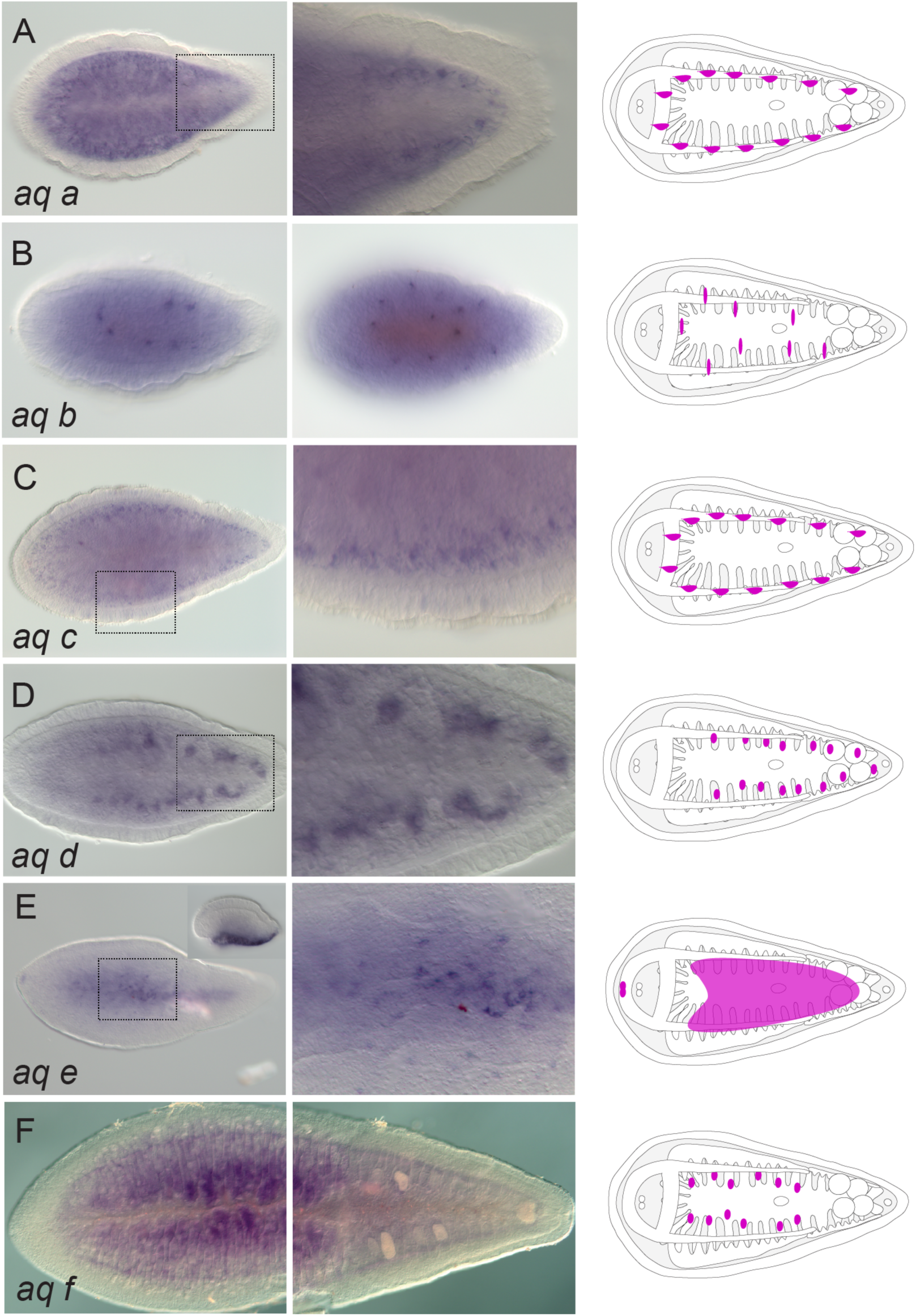
Expression of aquaporins in *M. stichopi.* Whole mount in situ hybridization (WMISH) of aquaporin genes. The inset in panel E shows a different focal plane (side view). The columns next to the panels show higher magnifications of the indicated domains except of B that shows a different view form the animal (side view). Illustrations with colored gene expression correspond to the animals shown in the previous column. Anterior is to the left. The depicted expression patterns are for guidance and not necessarily represent exact expression domains. Drawings are not to scale.

The broad expression of the ammonia excretion-related genes and *aquaporins* shows that these genes do not label defined excretory domains, as already seen with the *slcs* and ultrafiltration genes. However, since a considerable number of them (*rhesus*, members of *amt* (only in *I. pulchra*), *nka*, members of *ca, hcn* (only in *M. stichopi*), *v-ATPase B* as well as several *aquaporins)* were expressed in association with the digestion system, including different types of gut-wrapping and gut-affiliated (syncytial in *I. pulchra* and gut epithelial in *stichopi*) cells, the possibility that digestive-associated tissues could act as excretory sites was raised.

### High environmental ammonia exposure indicates a diffusion mechanism in *I. pulchra*

To reveal the excretory mechanism in Xenacoelomorphs, we conducted high environmental ammonia (HEA) incubation experiments, as previously performed in a large array of animals (summarized in^5,51^), using *I. pulchra* because of its availability in sufficient numbers. We first measured the pH of incubation mediums with different HEA concentrations (up to 1 mM) and found no difference in pH, which could otherwise have influenced any excretion rates. We then exposed animals to different HEA concentrations for a short period (2 hours) and measured the ammonia excretion during the following two hours after bringing them back into normal conditions, to test excretion via diffusion. The ammonia excretion rates of exposed animals remained unchanged after exposure to 50 and 100 µM NH_4_Cl, compared to the control conditions, but increased gradually after exposure to NH_4_Cl concentrations of 200 and 500 µM NH_4_Cl (Fig. 8a). The increase in ammonia excretion rate could be explained by a concentration-dependent ammonia uptake during the HEA exposure and a subsequent release in normal conditions. These results suggest that ammonia excretion is concentration-dependent, which is indicative of a diffusion mechanism.

**Figure 8.**
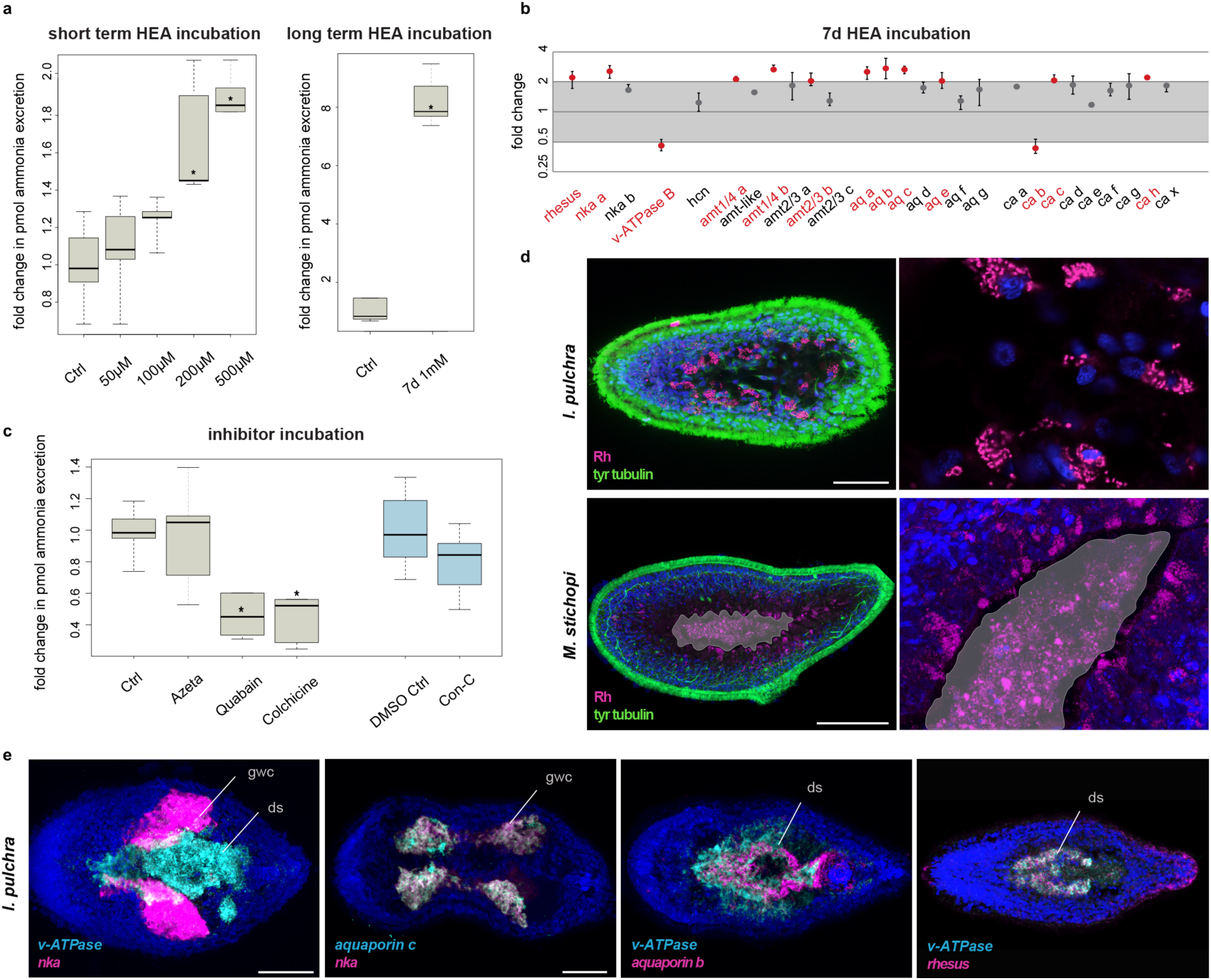
Excretion in acoelomorphs. (**a**) Ammonia excretion rates of *I. pulchra* before (control) and after exposure for 2 hours to 50, 100, 200 and 500 µM and after exposure for 7 days in 1mM NH_4_Cl (boxplot). Excretion was measured over two hours following the HEA treatments in at least three independent biological replicates, each divided into two separate samples (six measurements in total). Bold horizontal bar in boxes indicate the median, lower and upper box border indicate lower and upper quartile, whiskers indicate minimum and maximum. Asterisks label significant changes (p < 0.02 in an unpaired, 2-tailed t-test with unequal variance). (**b**) Quantitative relative expression of *rhesus, nka, v-ATPase B, amts, aquaporins*, and *ca* after 7 days exposure in HEA (1 mM NH_4_Cl). Each circle indicates the average of three independent biological replicates, each with four technical replicates. Error bars indicate minimum and maximum of the biological replicates (averaged technical replicates). One-fold change represents no change; ≥ 2 indicates significantly increased expression level; ≤ 0.5 indicates significantly decreased expression level (red labels). (**c**) Effects of different inhibitors on ammonia excretion rates in *I. pulchra (boxplot, with illustration and replicates similar to Figure 4a).* The concentrations used were 5 µM Concanamycin C as a v-ATPase A/B inhibitor, 1 mM Azetazolamide as an inhibitor of the CA, 1 mM Quabain as a NKA inhibitor and 2 mM Colchicine for inhibiting the microtubule network. Concanamycin C was diluted in 0,5% DMSO for which we used an appropriate control with 0,5% DMSO. (**d**) Protein localization of Rhesus in *I. pulchra* and *M. stichopi*. Syncytium and gut are indicated in grey and the magenta staining of the lumen in *M. stichopi* is false positive staining of the gut content. Fluorescent pictures are projections of merged confocal stacks. The nervous system is stained green with tyrosinated tubulin. (**e**) Double fluorescent WMISH of *v-ATPase* and *nka, aquaporin c* and *nka, v-ATPase* and *aquaporin b* and *v-ATPase* and *rhesus* in *I. pulchra.* White areas in the first panel are the result of merged stacks and not of overlapping expression. Nuclei are stained blue with 4’,6-diamidino-2-phenylindole (DAPI). Anterior is to the left. Abbreviations: ctrl, control; ds, digestive syncytium; gwc, gut-wrapping cells. Scale bars, 50 µm for *I. pulchra* and 100 µm for *M. stichopi.*

### HEA exposure influences the expression of some excretion-related genes in *I. pulchra*

To test for a possible involvement of the excretion-related genes in the excretory mechanism of xenacoelomorphs, we tested for alteration of mRNA expression levels in chronically HEA exposed animals by quantitative relative expression experiments (qPCR) in *I. pulchra*. We first exposed animals to 1mM HEA for 7 days, similar to conditions used in previous studies (summarized in^5,51^), and measured the ammonia excretion over 2 hours after bringing the animals into normal conditions. As expected, the ammonia excretion rates were strongly increased, in line with the short termed HEA exposure experiments (Fig. 8a). When we tested the expression level of excretion-related genes, we found that the expression of the passive ammonia transporters *rhesus* and three *amts*, as well as the active ammonia transporter *nka* altered significantly (Fig. 8b). Other differentially expressed genes were four *aquaporins*, the *v-ATPase*, three *cas* and four *slcs* (Fig. 8b and Supplementary Fig. 5). These results indicate a putative role of these genes in ammonia excretion and suggest that acoels might not only excrete by diffusion and via passive transporters (*rhesus, amts*), but also by an alternative active transport mechanism (*nka*).

### Inhibitor experiments support an active excretion mechanism via NKA transporter as well as a passive vesicular transport mode, possibly mediated by Rhesus transporter

We further tested the involvement of NKA, v-ATPase A/B and CA proteins in excretion, as well as a possible involvement of a vesicular transport mechanism, by conducting pharmacological inhibitor assays in *I. pulchra*, as previously demonstrated in other animals (summarized in^5,51^). Inhibition of the CA by Azetazolamide, did not show any significant change in ammonia excretion. Inhibition of the V-ATPase by Concanamycin C seemed to lead to a decrease in ammonia excretion, although a 2-tailed t-test did not support a significant change. In contrast, when perturbing the function of NKA with Quabain, the ammonia excretion dropped significantly (Fig. 8c), which further support an active excretion mechanism via NKA, similar to what is described for many nephrozoans^10-13,16,18,19,24-26,30,52-54^. Interference with the vesicular transport using Colchicine also led to a significant decrease in ammonia excretion, indicating a possible vesicular ammonia-trapping excretion mode (Fig. 8c), as demonstrated in the midgut epithelium of the tobacco hornworm^25^, the gills of the shore crab^26^ and the integument of the nematode^13^. To test whether vesicular transport might occur through Rhesus transporter as shown in other studies (summarized in^5,51^), we revealed Rhesus protein localization by immunohistochemistry (Fig. 8d). The protein localization mimicked the gene expression and revealed, apart of cells at the anterior tip and cells of the posterior epidermis, individual parenchymal cells affiliated with the digestive syncytium that extend ventrally. Higher magnification showed that the transporter was present in cytoplasmic vesicles and not on the cellular membrane (Fig. 8d). This further indicated the presence of a vesicular transport mechanism, in which cellular ammonia moves via Rhesus transporters into vesicles and gets transferred to the membrane through the microtubule network^55^ (Fig. 1c). The antibody specificity was confirmed by an alignment of the epitope and the endogenous protein, as a well as a western blot analysis (Supplementary Fig. 6). Similar vesicular protein localization was also observed in *M. stichopi*, suggesting a similar cytoplasmic-vesicular role of Rhesus transporter in gut-affiliated cells, also in nemertodermatids (Fig. 8d). These data further supported the involvement of NKA and Rhesus transporters in ammonia excretion and also indicated the presence of a putative vesicular transport mode in *I. pulchra.*

### Double fluorescent WMISH of differentially expressed genes shows similar spatial arrangement in gut-associated domains in *I. pulchra* and *M. stichopi*

To obtain a better resolution and understanding of the relative topology of the differentially expressed genes, double fluorescent WMISH were conducted for *v-ATPase, nka* and *aquaporins b* and *c* and *rhesus* (Fig. 8e). *Nka* and *v-ATPase* were expressed in a mutually exclusive manner, with *v-ATPase* to be restricted in the ventral region digestive syncytium and *nka* in an adjacent parenchymal, distal sac-shaped gut-wrapping domain. *Aquaporin c* was coexpressed with *nka* and *aquaporin b* was partially overlapping with *v-ATPase*, with *aquaporin b* expression extending into the posterior region of the digestive syncytium. Finally, *rhesus* was partly coexpressed with *v-ATPase* in the ventral region of the digestive syncytium. Similar coexpression analysis of the orthologous genes was also conducted for *M. stichopi* and revealed striking similarities in their spatial arrangement to *I. pulchra* (Supplementary Fig. 7). *V-ATPase* expression was not overlapping with *nka*, as *v-ATPase* was restricted to the gut epithelium and in two proximal lateral rows of subepidermal cells whilst *nka* was limited to cells lining the distal part of the epithelial branches of the gut extending towards the subepidermis. *V-ATPase* was partly coexpressed with *rhesus* in ventral gut-affiliated cells. Overall, these data revealed a similar spatial arrangement in gut-associated domains in both animals, which seems to be unrelated to the presence of an epithelial gut or a syncytium.

Taken together, our findings suggest that *I. pulchra* uses different mechanisms for ammonia excretion that are also known from nephrozoans; an active ammonia excretion mechanism via NKA through the digestive system, as suggested by in situ hybridization, and a passive vesicular transport mechanism likely mediated by Rhesus through digestive and likely also epidermal tissues. Given the commonalities in the expression of the involved genes in both animals, these excretory mechanisms could be plesiomorphic for Xenacoelomorpha.

### HEA experiments suggest a diffusion mechanism also in the cnidarian *Nematostella vectensis*

Because our results showed the involvement of active and passive transport mechanisms across digestive tissues outside Nephrozoa, we also investigated a non-bilaterian species, the cnidarian *Nematostella vectensis* (Supplementary Fig. 1b), to test whether this excretion mode might also be present outside Bilateria. The only available excretion studies in cnidarians are few morphological studies, which suggested that the septa filaments of the anthozoan mesenteries and the radial canals of medusozoans could serve as putative excretory sites^56^, as well as some isotopic exchange experiments on *Hydra oligactis* that have shown that the gastrodermis seems to be involved in osmoregulation^57^. Moreover, there is evidence that Rh and AMT genes are generally involved in ammonium excretion in corals^58^, but localization studies that would suggest excretion sites are missing.

We first tested whether *N. vectensis* excretes via diffusion by exposing early juvenile animals to HEA for 2 hours and measuring their ammonia excretion rates afterwards, similar to the experiments performed with *I. pulchra* (Fig. 9a). We found that also in *N. vectensis* ammonia excretion increased significantly after HEA exposure starting at 200 and 500 µM NH_4_Cl (Fig. 9a). Measurements of the pH of each incubation medium showed that the pH dropped by 0.2 when the medium contained 500 µM NH_4_Cl. However, when we measured the excretion of animals over 2 hours in a medium with an accordingly lowered pH we found that a difference of 0.2 did not change the excretion rates (Supplementary Table 2). Therefore, the increase in ammonia excretion rates at 200 and 500 µM NH_4_Cl indicates that ammonia excretion is concentration-dependent, supporting a diffusion mechanism also in *N. vectensis*.

**Figure 9.**
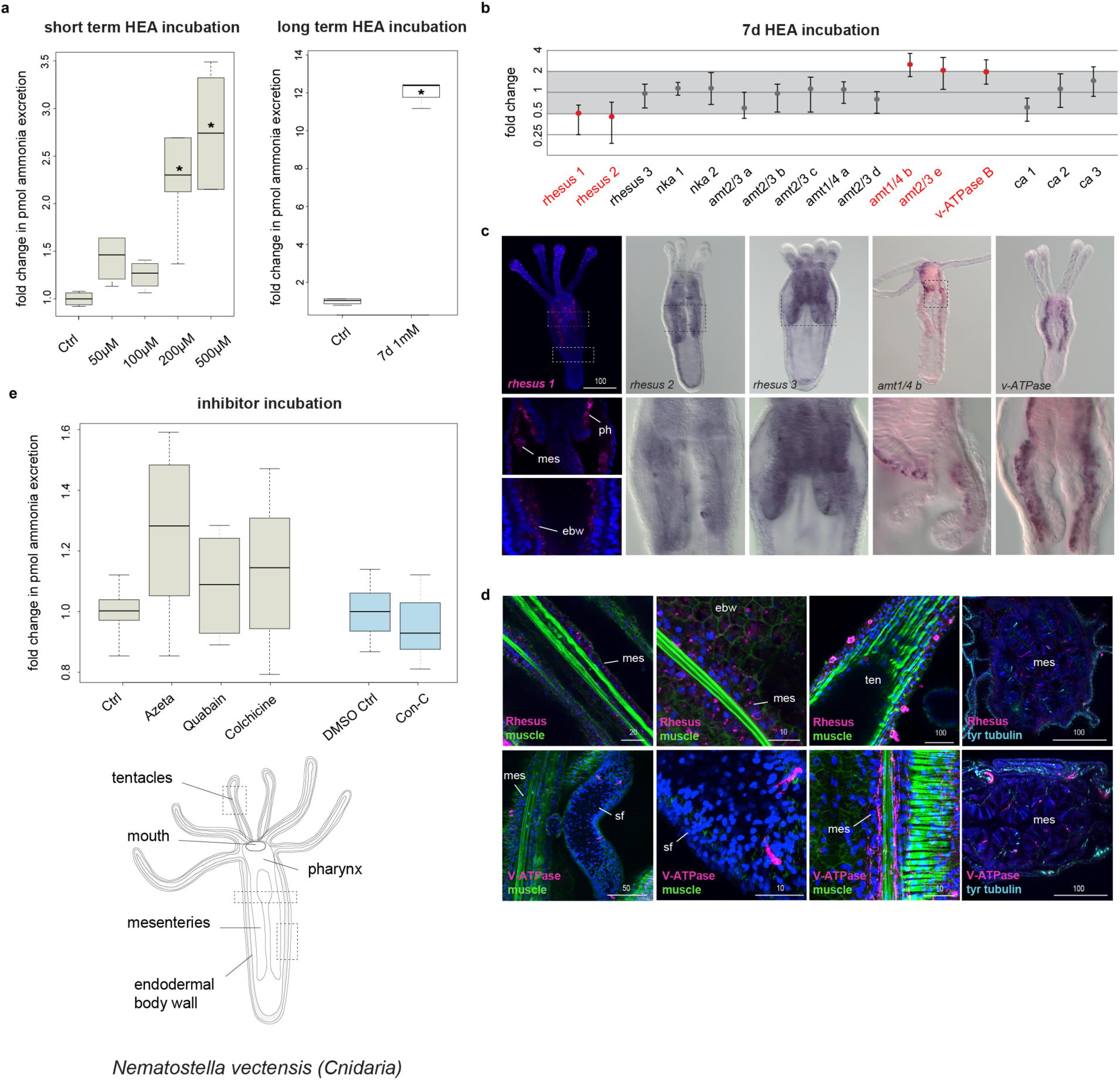
Excretion in *Nematostella vectensis*. (**a**) Ammonia excretion rates of *N. vectensis* before (control) and after exposure for 2 hours to 50, 100, 200 and 500 µM and after exposure for 7 days in 1mM NH_4_Cl (boxplot). Excretion was measured over two hours following the HEA treatments in at least 3 independent biological replicates, each divided into two separate samples (6 measurements in total). Bold horizontal bar in boxes indicate the median, lower and upper box border indicate lower and upper quartile, whiskers indicate minimum and maximum. Asterisk labels significant changes. Significance, p < 0.02 (unpaired t-test with unequal variance). (**b**) Quantitative relative expression of *rhesus, nks, v-ATPase B, amts* and *ca* after exposure for 7 days in HEA (1 mM NH_4_Cl). Each circle represents the average of five independent biological replicates, each with three technical replicates. One fold change represents no change; ≥ 2 indicates increased expression level significantly; ≤ 0.5 indicates decreased expression level significantly (red labels). (**c**) Whole mount in situ hybridization of *rh 1, rh 2, rh 3, v-ATPase B* and *amt6* in feeding primary polyps. Anterior is to the top. (**d**) Protein localization of Rh and v-ATPase in *N. vectensis* early juvenile polyps. The muscle filaments are labeled green with phalloidin and the nervous system is stained cyan with tyrosinated tubulin. Every picture is a full projection of merged confocal stacks. Nuclei are stained blue with 4’,6-diamidino-2-phenylindole (DAPI). The regions shown are indicated with dashed boxes in the illustrated animal. (**e**) Effects of different inhibitors on ammonia excretion rates in *N. vectensis (boxplot, with illustration and replicates similar to Figure 5a).* The concentrations used were 5-15 µM Concanamycin C as a v-ATPase A/B inhibitor, 1-3 mM Azetazolamide as an inhibitor of the CA, 1-5 mM Quabain as a NKA inhibitor and 2-10 mM Colchicine for inhibiting the microtubule network. Quabain was diluted in 0,5% DMSO for which we used an appropriate control with 0,5% DMSO. N=3 for all treatments. Abbreviations: ctrl, control; ebw, endodermal body wall; mes, mesenteries; ph, pharynx; sf, septal filament; ten, tentacles.

### Quantitative gene expression and inhibitor experiments indicates an involvement of passive but not active transport mechanisms in *N. vectensis*

We then exposed animals to 1mM HEA for 7 days and tested the expression of the orthologous genes altered in *I. pulchra* treatments (*rh/amt, nka, v-ATPase B* and *ca*) by qPCR. As expected from the short-term HEA exposure experiment, specimens exposed for 7 days in HEA condition showed increased ammonia excretion rates (Fig. 9a). Also, the expression of the passive transporters *rhesus1, rhesus2, amt6* and *amt7* as well as *v-ATPase* altered significantly (Fig. 9b). In contrast to *I. pulchra*, none of the two *nka* transporters showed a significant change in expression. To test whether Rhesus acts via a vesicular transport mechanism we conducted the same pharmacological experiment as in *I. pulchra*. Contrary to the results from acoels, we found that inhibition of vesicular transport did not alter the ammonia excretion (Fig. 9e). We also inhibited the excretory function of v-ATPase and CA proteins and found that none of them showed any significant change in ammonia excretion rates (Fig. 9e). Finally, when we perturbed the function of NKA the ammonia excretion rates did not alter (Fig. 9e), confirming the qPCR results and further supporting the non-involvement of this transporter in excretion (Fig. 9a). These results suggest that the ammonia excretion of *N. vectensis* is likely mediated via the passive Rhesus and AMT transporters but neither relies on active transport mechanism mediated by NKA or on vesicular ammonia-trapping excretion mode (Fig. 9e).

### Gene expression of excretion related genes reveals the gastrodermis as excretory site in *N. vectensis*

To understand whether these genes were expressed in gastrodermal or epidermal cells, we revealed the spatial expression of *rhesus, amts, nka* and *v-ATPase B* by WMISH in feeding primary polyps. All genes were mainly demarcating gastrodermal domains, such as the endodermal body wall, the directive mesenteries, septal filaments and the pharynx (Fig. 9c and Supplementary Fig. 8). *Rhesus 1* was additionally expressed in the tentacular ectoderm (Fig. 9c). Protein localization of Rh, NKA and V-ATPase B revealed by immunohistochemistry reflected the transcript expression patterns (Fig. 9d and Supplementary Fig. 9). High magnification of Rhesus antibody staining further revealed that the transporter was not expressed in cytoplasmic vesicles supporting a non-vesicular transport mechanism, in agreement to the inhibitor experiments. Also, it showed that Rhesus was localized in individual cells of the tentacular ectoderm with clumped structures at the tentacle surface, which resembled gland cells^59^ (Fig. 9d). The NKA antibody was localized in endodermal neurons and individual cells of the mesenteries, likely neural precursors (Supplementary Fig. 9), thus suggesting a non-excretory function of this transporter, as expected from the qPCR and inhibitor experiments. These data imply that gastrodermis affiliated domains likely serve as excretory sites in *N. vectensis*.

## Discussion

Overall, our findings show that acoelomorphs use, in addition to diffusion, also active transport mechanisms, in contrast to what has been previously assumed for non-nephrozoans^1,38,39^. Our results also suggest that excretion takes place across digestive tissues and likely also across the epidermis, as indicated from the *rhesus* (both animals) and *amt* (only in *M. stichopi*) expression. *N. vectensis* also seem to use gastrodermal tissues as excretory sites but we were not able to detect any active transport mechanism. However, we don’t know whether the absence of active transport is common to all cnidarian excretion or just a feature of *N. vectensis.* More studies in other members of Cnidaria are necessary to elucidate this issue. Interestingly, digestive tissues with additional or assigned excretory roles have also been reported in several nephrozoans (e.g. vertebrates, annelids, insects, nematodes, tunicates, chaetognaths)^15,25,30,60-66^. In the light of our results this excretion mechanism likely reflects an ancient mechanism, before the evolution of specialized organs, such as nephridia (Supplementary Fig. 10). The molecular spatial arrangement of the excretion sites in non-nephrozoans, however, is not sharing topological arrangements with common nephridial domains of nephrozoans (Fig. 1b), suggesting that they are evolutionary unrelated to nephridia. It still remains unclear whether these domains are multifunctional or consist of specialized excretory subdomains, however a degree of cell sub-functionalization seems to be present, as indicated by the localized gene expression in different groups of gut wrapping and gut epithelial cells. We can however exclude the presence of ultrafiltration sites, in agreement with previous ultrastructure studies in acoelomorphs^49^, since the homologous essential molecular components of the ultrafiltration sites of nephridia and nephrocytes are mostly expressed in neural domains in acoelomorphs, and are absent in non-bilaterians (nephrins), suggesting their later recruitment in the nephrozoan filtration apparatus’ (Fig. 10).

**Figure 10.**
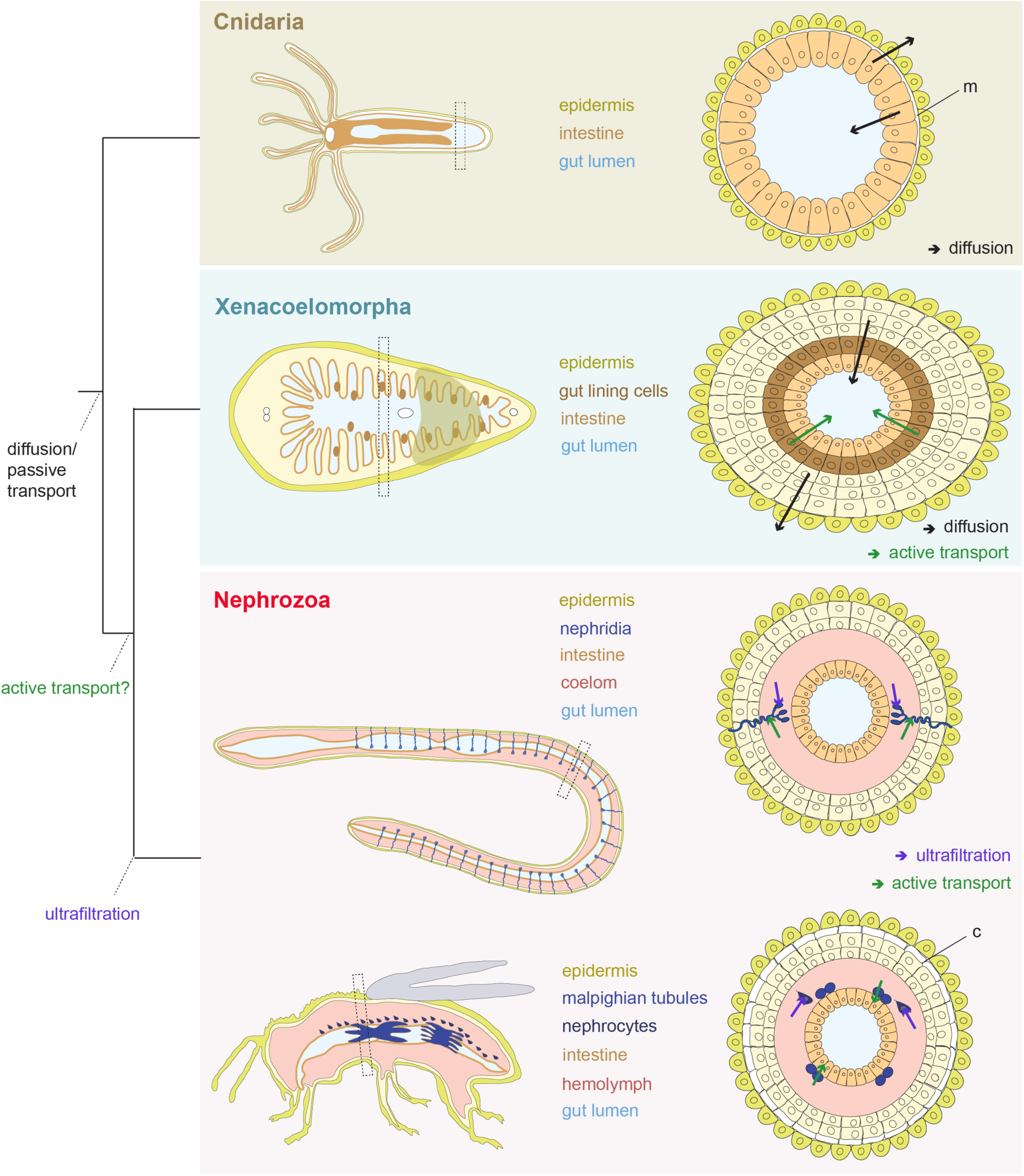
Evolution of excretory mechanisms. Illustration of the proposed direction of fluxes in Cnidaria and Xenacoelomorpha, and evolution of active ammonia transport and ultrafiltration mechanisms. Cnidaria excrete across their intestinal epithelium via diffusion whilst in Xenacoelomorphs excretion occurs both via diffusion across the epidermis and gut-associated tissues and via active transport across gut-associated tissues. Ultrafiltration mechanism originated within Nephrozoa. Abbreviations: m, mesoglea; c, cuticle.

To conclude, our study shows that active transport mechanism and excretion through digestive tissues predates the evolution of specialized excretory systems. If this is based on a convergent recruitment, or if it reflects an ancestral state for Bilateria remains unclear. However, if the latter is true, it correlates with the emergence of multilayered body plans and solid internal parenchymes that separate the body wall from their digestive tract, as seen in xenacoelomorphs and nephrozoans. We thus propose that diffusion mechanisms were the major excretory modes present in animals with single-layered epithelial organization (Fig. 10). The emergence of more complex, multilayered body plan necessitated an active transport of excretes, which was later recruited in specific compartments of the complex excretory organs in the lineage of Nephrozoa.

## Methods

No statistical methods were used to predetermine sample size. The experiments were not randomized. The investigators were not blinded to allocation during experiments and outcome assessment.

### Gene cloning and orthology assignment

Putative orthologous sequences of genes of interest were identified by tBLASTx search against the transcriptome (SRR2681926) of *Isodiametra pulchra*, the transcriptome (SRR2681155) and draft genome of *Meara stichopi* and the genome of *Nematostella vectensis* (http://genome.jgi.doe.gov). Additional transcriptomes of Xenacoelomorpha species investigated were: *Childia submaculatum* (Acoela) (SRX1534054), *Convolutriloba macropyga* (Acoela) (SRX1343815), *Diopisthoporus gymnopharyngeus* (Acoela) (SRX1534055), *Diopisthoporus longitubus* (Acoela) (SRX1534056), *Eumecynostomum macrobursalium* (Acoela) (SRX1534057), *Hofstenia miamia* (Acoela) (PRJNA241459), *Ascoparia* sp. (Nemertodermatida) (SRX1343822), *Nemertoderma westbladi* (Nemertodermatida) (SRX1343819), *Sterreria* sp. (Nemertodermatida) (SRX1343821), *Xenoturbella bocki* (*Xenoturbella*) (SRX1343818) and *Xenoturbella profunda* (Xenoturbella) (SRP064117). Sequences for the placozoan *Trichoplax adhaerens*, the sponge *Amphimedon queenslandica*, the ctenophore *Mnemiopsis leidy*, the protist *Capsaspora owczarzaki*, the amoeba *Dictyostelium discoideum* and the nephrozoans *Homo sapiens, Saccoglossus kowalevskii, Strongylocentrotus purpuratus, Xenopus laevis, Branchiostoma lanceolatum, Capitella teleta, Crassostrea gigas, Lottia gigantea, Schmidtea mediterranea, Tribolium castaneum, Caenorhabditis elegans* and *Drosophila melanogaster* were obtained from Uniprot and NCBI public databases. Gene orthology of genes of interest identified by tBLASTx was tested by reciprocal BLAST against NCBI Genbank and followed by phylogenetic analyses. Amino acid alignments were made with MUSCLE^67^. RAxML (version 8.2.9)^68^ was used to conduct a maximum likelihood phylogenetic analysis. Fragments of the genes of interest were amplified from cDNA of *I. pulchra, M. stichopi* and *N. vectensis* by PCR using gene specific primers. PCR products were purified and cloned into a pGEM-T Easy vector (Promega, USA) according to the manufacturer’ s instruction and the identity of inserts confirmed by sequencing. Gene accession numbers of the gene sequences are listed in the Supplementary Table 3.

### Data Availability

All newly determined sequences have been deposited in GenBank under accession numbers XXX-XXX. Primer sequences are available on request.

### Animal systems

Adult specimens of *Isodiametra pulchra* (Smith & Bush, 1991), *Meara stichopi* Westblad, 1949 and *Nematostella vectensis* Stephenson, 1935 were kept and handled as previously described^69-72^.

### Whole Mount In Situ Hybridization

Animals were manually collected, fixed and processed for *in situ* hybridization as described^73,74^. Labeled antisense RNA probes were transcribed from linearized DNA using digoxigenin-11-UTP (Roche, USA), or labeled with DNP (Mirus, USA) according to the manufacturer’s instructions. For *I. pulchra* and *M. stichopi*, colorimetric Whole Mount In Situ Hybridization (WMISH) was performed according to the protocol outlined in^73^. For *N. vectensis*, we followed the protocol described by^75^. Double Fluorescent In Situ Hybridization (FISH) was performed as the colorimetric WMISH with the following modifications: After the post-hybridization steps, animals were incubated overnight with peroxidase-conjugated antibodies at 4°C (anti-DIG-POD, Roche 1:500 dilution and anti-DNP, Perkin Elmer, 1:200 dilution) followed by the amplification of the signal with fluorophore-conjugated tyramides (1:100 TSA reagent diluents, Perkin Elmer TSA Plus Cy3 or Cy5 Kit). Residual enzyme activity was inhibited via 45-minute incubation in 0.1% hydrogen peroxide in PTW followed by four PTW washes prior to addition and development of the second peroxidase-conjugated antibody^76^.

### Whole Mount Immunohistochemistry

Animals were collected manually, fixed in 4% paraformaldehyde in SW for 50 minutes, washed 3 times in PBT and incubated in 4% sheep serum in PBT for 30 min. The animals were then incubated with commercially available primary antibodies (anti-RhAG (ab55911) rabbit polyclonal antibody, dilution 1:50 (Abcam, USA), anti-Na^+^/K^+^ ATPase (a1 subunit) rat monoclonal antibody, dilution 1:100 (Sigma-Aldrich, USA) and anti-v-ATPase B1/2 (L-20) goat polyclonal antibody, dilution 1:50 (Biotechnology, Santa Cruz, USA) overnight at 4°C, washed 10 times in PBT, and followed by incubation in 4% sheep serum in PBT for 30 min. Specimens were then incubated with a secondary antibody (anti rabbit-AlexaFluor 555, Invitrogen or anti rat-AlexaFluor 555 and anti-goat-AlexaFluor 555) diluted 1:1000 overnight at 4°C followed by 10 washes in PBT. Nuclei were stained by incubation of animals in DAPI 1:1000 and f-actin was stained by incubation in BODIPY labeled phallacidin 5 U/ml overnight.

### Inhibitor and HEA experiments

For excretion experiments approximately 300 *I. pulchra* (the number varied slightly between the biological replica, but was similar in the corresponding controls and treatments) and 10 *N. vectensis* were placed into glass vials with 2 ml UV sterilized natural seawater (1:4 diluted with distilled water for *N. vectensis*) containing the appropriate inhibitor or ammonia concentration. Animals were given 10 minutes to adjust to the medium before the solution was exchanged with 2 ml of fresh medium with the same appropriate condition. For the inhibitor experiment the medium was removed after 2 hours and stored at −80°C for later measurements. Animals from the short-term high environmental ammonia (HEA) experiments were incubated for 2 hours, then rinsed 5 times over 20-30 minutes and incubated for another 2 hours in fresh medium without additional ammonia, after which the medium was removed and frozen at −80°C. We tested different inhibitor concentrations that were used in previous studies in other invertebrates^10,12,13,18,26^. The concentrations of 5-15 µM Concanamycin C for inhibiting v-ATPase A/B, 1-3 mM Azetazolamide as an inhibitor of the CA, 1-5 mM Quabain to inhibit the NKA and 2-10 mM Colchicine for inhibiting the microtubule network were selected, as no other effects like shrinking or obvious changes in morphology or behavioral were observed. After the inhibitor incubations, the animals were washed several times and monitored in normal conditions for several days to ensure that the inhibitors did not cause any unspecific permanent effects. Concanamycin C was diluted in DMSO with a final concentration of 0.5% DMSO per sample for which we used an appropriate control with 0.5% DMSO. For the HEA experiments we enriched seawater with NH_4_Cl to the final ammonia concentrations of 50 µM, 100 µM, 200 µM, 500 µM, and 1 mM. We also measured the pH of both incubation mediums (HEA and control) and we found no difference. All experiments were independently repeated at least three times at different time points, and each repeat was divided into two samples.

### Determination of ammonia excretion

Ammonia concentrations were measured with an ammonia sensitive electrode (Orion, Thermo Scientific) according to^53^. Samples were diluted 1:4 with distilled water to prevent salt precipitation (900 µl sample + 2.7 ml water) and total ammonia was transformed into gaseous NH_3_ by adding 54 µl ionic strength adjusting solution (1.36 ml/l trisodiumcitrate dihydrate, 1 M NaOH). Due to the small ammonia concentrations the electrode-filling solution was diluted to 10% with distilled water, as suggested in the electrode manual. In control conditions we determined an average excretion of 44 pmol per adult animal per hour, although the excretion varied between different biological replicates from different generations (min. 32 pmol/animal/hour, max. 52 pmol/animal/hour), possibly due to slightly fluctuating conditions during long-term animal rearing. Solutions with defined concentrations of NH_4_Cl for the standard curves were made together with the solutions used in the experiments and stored in a similar way at −80°C. The differences in excretion rates were tested for significance with an unpaired, 2-tailed t-test with unequal variance and a p value < 0.02 was seen as significant. Boxplots were created with ‘R’.

### Quantitative Gene Expression

100 treated *I. pulchra* and 5 *N. vectensis* were collected after 7 days of incubation in HEA conditions (1 mM NH_4_Cl) and tested for quantitative gene expression using the BIORAD CFX96 Real time PCR detection system. ddCt values were calculated between treated and control animals and converted to fold differences. All experiments were repeated three to five times with different specimens (three biological replicates for *I. pulchra* and five biological replicates for *N. vectensis*) and two to dour technical replicates were tested for each biological replicate (four biological replicates for *I. pulchra* and three biological replicates for *N. vectensis*). Fold changes were calculated using polyubiquitin, actin and 18S as references for *I. pulchra* and *ATPsynthase* and *EF1b* as references for *N. vectensis*^77^, and a threshold of two-fold difference was chosen as a significant change. The Ct values are provided in Supplementary Table 4 and the primer sequences used are provided in Supplementary Table 5.

### Western blot

Whole animal extracts (50 *I. pulchra* adults and 5 *N. vectensis* juveniles) were fractionated by SDS-PAGE, loaded on Mini-PROTEAN® TGX Stain-Free Precast Gels (Bio-Rad) and transferred to a nitrocellulose membrane using a transfer apparatus according to the manufacturer’s protocols (Bio-Rad). After incubation with 5% nonfat milk in TBST (10 mM Tris, pH 8.0, 150 mM NaCl, 0.5% Tween 20) for 60 min, the membrane was washed once with TBST and incubated with antibodies against Rhesus (1:1000) and NKA (1: 500) at 4 °C for 12 h. Membranes were washed three times for 10 min and incubated with a 1:5000 dilution of horseradish peroxidase-conjugated anti-mouse or anti-rabbit antibodies for 2 h. Blots were washed with TBST three times and developed with the ECL system (Amersham Biosciences) according to the manufacturer’s protocols.

### Documentation

Colorimetric WMISH specimens were imaged with a Zeiss AxioCam HRc mounted on a Zeiss Axioscope A1 equipped with Nomarski optics and processed through Photoshop CS6 (Adobe). Fluorescent-labeled specimens were analyzed with a SP5 confocal laser microscope (Leica, Germany) and processed by the ImageJ software version 2.0.0-rc-42/1.50d (Wayne Rasband, NIH). Figure plates were arranged with Illustrator CS6 (Adobe).

## Acknowledgments

We thank Fabian Rentzsch, Patrick Steinmetz and Hanna Kraus (Sars Centre, University of Bergen, Norway) for providing *N. vectensis* animals and cDNA. Peter Ladurner provided the *I. pulchra* cultures that were kept in the animal facility at the Sars Centre. We thank all S9 lab members for the help with the collections of *M. stichopi*. The study was supported by the core budget of the Sars Centre, the European Research Council Community’s Framework Program Horizon 2020 (2014–2020) ERC grant agreement 648861 (EVOMESODERM) and FP7-PEOPLE-2012-ITN grant no. 317172 (NEPTUNE) to A.H. and Coca-Cola Foundation to J.A.R-S..

## Author contributions

C.A. and A.H. designed the study. C.A., D.T. and J.A.R.-S. performed the experiments. C.A. analyzed the data and C.A. and A.H. wrote the manuscript. All authors read and approved the final manuscript.

