## Supplementary Files for "Active mode of excretion across digestive tissues predates the origin of excretory organs"

**Index**

- Supplementary Figures 1-10
- Supplementary Tables 1-5

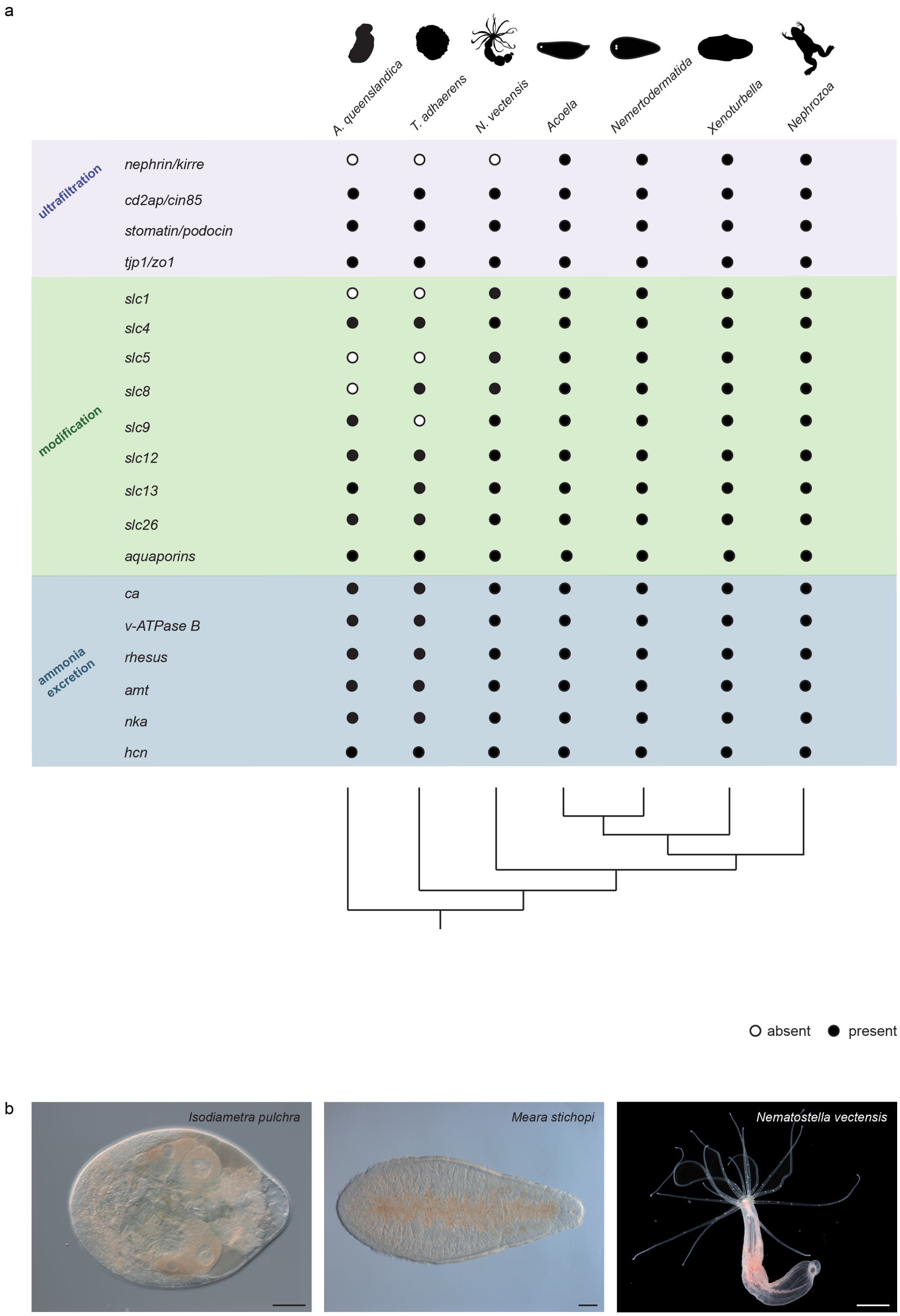

**Supplementary Figure 1.** (**a**) Excretion-related gene complement in different animal lineages and outgroups. Transcriptome and genome mining of excretion-related gene repertoire in *Isodiametra pulchra, Hofstenia miamia, Convolutriloba macropyga, Diopisthoporus longitubus, Diopisthoporus gymnopharyngeus, Eymecynostomum macrobursalium* and *Childia submaculatum* as representatives of Acoela, *Sterreria* sp., *Ascoparia* sp., *Meara*

*stichopi* and *Nemertoderma westbladi* as representatives of Nemertodermatida, *Xenoturbella bocki* and *Xenoturbella profunda* as representatives of *Xenoturbella, Nematostella vectensis* as a representative of cnidarians, *Trichoplax adhaerens* as a representative of placozoans, *Amphimedon queenslandica* as a representative of sponges and the deuterostomes *Homo sapiens, Saccoglossus kowalevskii, Strongylocentrotus purpuratus, Xenopus laevis, Branchiostoma lanceolatum* and protostomes *Capitella teleta, Crassostrea gigas, Lottia gigantea, Schmidtea mediterranea, Tribolium castaneum, Caenorhabditis elegans* and *Drosophila melanogaster* as representatives of Nephrozoa. Data are based on this study unless stated otherwise. (**b**) Pictures of the acoelomorph representatives *Isodiametra pulchra* (scale bar = 50 µm) and *Meara stichopi* (scale bar = 100 µm), and the cnidarian representative *Nematostella vectensis* (scale bar = 2 mm). Animal illustrations are taken from phylopic.org.

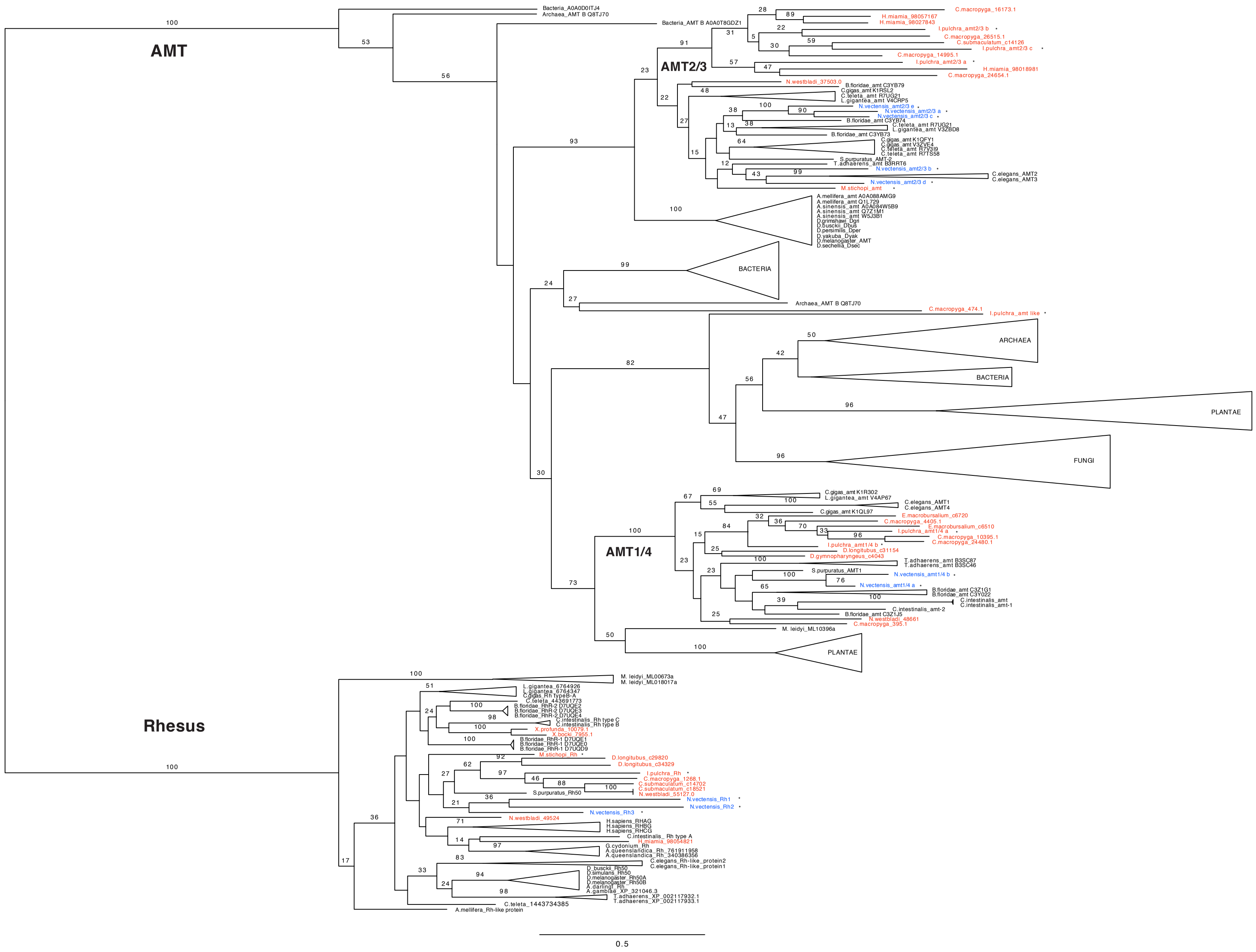
AMT
AMT2/3
AMT1/4
BACTERIA
ARCHAEA
PLANTAE
FUNGI
Rhesus
0.5

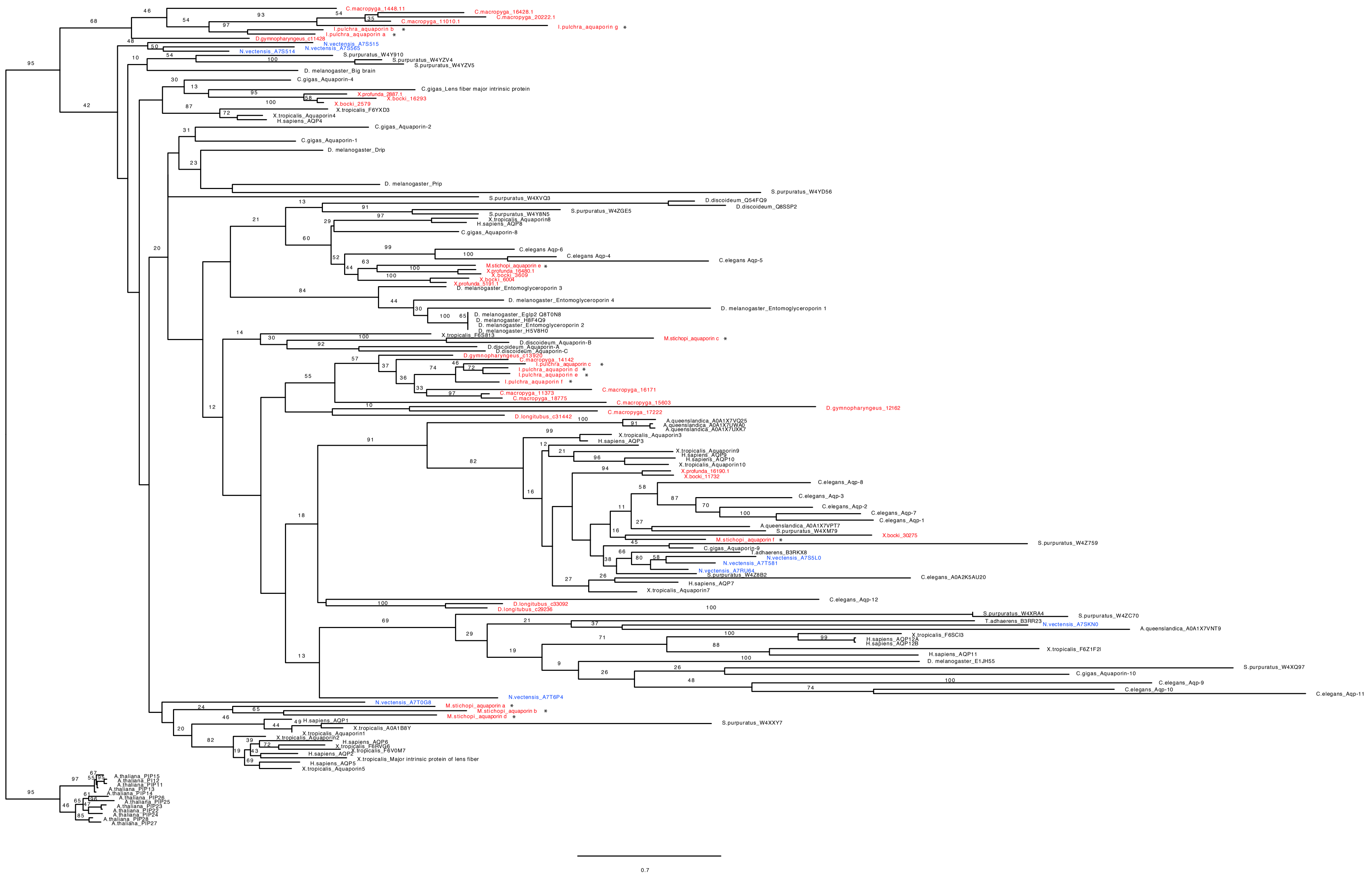
C.macropyga_1448.11
C.macropyga_16428.1
C.macropyga_20222.1
C.macropyga_11010.1
I.pulchra_aquaporin g *
I.pulchra_aquaporin b *
I.pulchra_aquaporin a *
D.gymnopharyngeus_c11428
N.vectensis_A7S515
N.vectensis_A7S565
N.vectensis_A7S514
S.purpuratus_W4Y910
S.purpuratus_W4YZV4
S.purpuratus_W4YZV5
D. melanogaster_Big brain
C.gigas_Aquaporin-4
C.gigas_Lens fiber major intrinsic protein
X.profunda_2887.1
X.bocki_16293
X.bocki_2579
X.tropicalis_F6YXD3
X.tropicalis_Aquaporin4
H.sapiens_AQP4
C.gigas_Aquaporin-2
C.gigas_Aquaporin-1
D. melanogaster_Drip
D. melanogaster_Prip
S.purpuratus_W4YD56
S.purpuratus_W4XVQ3
D.discoideum_Q54FQ9
D.discoideum_Q8SSP2
S.purpuratus_W4ZGE5
S.purpuratus_W4Y8N5
X.tropicalis_Aquaporin8
H.sapiens_AQP8
C.gigas_Aquaporin-8
C.elegans Aqp-6
C.elegans Aqp-4
C.elegans Aqp-5
M.stichopi_aquaporin e *
X.profunda_16480.1
X.bocki_3609
X.bocki_6004
X.profunda_5191.1
D. melanogaster_Entomoglyceroporin 3
D. melanogaster_Entomoglyceroporin 4
D. melanogaster_Entomoglyceroporin 1
D. melanogaster_Eglp2 Q8T0N8
D. melanogaster_H8F4Q9
D. melanogaster_Entomoglyceroporin 2
D. melanogaster_H5V8H0
X.tropicalis_F6S813
M.stichopi_aquaporin c *
D.discoideum_Aquaporin-B
D.discoideum_Aquaporin-A
D.discoideum_Aquaporin-C
D.gymnopharyngeus_c13920
C.macropyga_14142
I.pulchra_aquaporin c *
I.pulchra_aquaporin d *
I.pulchra_aquaporin e *
I.pulchra_aquaporin f *
C.macropyga_16171
C.macropyga_11373
C.macropyga_18775
D.gymnopharyngeus_12162
C.macropyga_15603
C.macropyga_17222
D.longitubus_c31442
A.queenslandica_A0A1X7VQ25
A.queenslandica_A0A1X7UWA0
A.queenslandica_A0A1X7UXK7
X.tropicalis_Aquaporin3
H.sapiens_AQP3
X.tropicalis_Aquaporin9
H.sapiens_AQP9
H.sapiens_AQP10
X.tropicalis_Aquaporin10
X.profunda_16190.1
X.bocki_11732
C.elegans_Aqp-8
C.elegans_Aqp-3
C.elegans_Aqp-2
C.elegans_Aqp-7
C.elegans_Aqp-1
A.queenslandica_A0A1X7VPT7
S.purpuratus_W4XM79
X.bocki_30275
M.stichopi_aquaporin f *
S.purpuratus_W4Z759
C.gigas_Aquaporin-9
T.adhaerens_B3RKX8
N.vectensis_A7S5L0
N.vectensis_A7T581
N.vectensis_A7RU64
S.purpuratus_W4Z8B2
C.elegans_A0A2K5AU20
H.sapiens_AQP7
X.tropicalis_Aquaporin7
C.elegans_Aqp-12
D.longitubus_c33092
D.longitubus_c29236
S.purpuratus_W4XRA4
S.purpuratus_W4ZC70
T.adhaerens_B3RR23
N.vectensis_A7SKN0
A.queenslandica_A0A1X7VNT9
X.tropicalis_F6SCI3
H.sapiens_AQP12A
H.sapiens_AQP12B
X.tropicalis_F6Z1F2I
H.sapiens_AQP11
D. melanogaster_E1JH55
S.purpuratus_W4XQ97
C.gigas_Aquaporin-10
C.elegans_Aqp-9
C.elegans_Aqp-10
C.elegans_Aqp-11
N.vectensis_A7T6P4
N.vectensis_A7T0G8
M.stichopi_aquaporin a *
M.stichopi_aquaporin b *
M.stichopi_aquaporin d *
H.sapiens_AQP1
S.purpuratus_W4XXY7
X.tropicalis_A0A1B8Y
X.tropicalis_Aquaporin1
X.tropicalis_Aquaporin2
H.sapiens_AQP6
X.tropicalis_F6RVG6
X.tropicalis_F6V0M7
H.sapiens_AQP2
X.tropicalis_Major intrinsic protein of lens fiber
H.sapiens_AQP5
X.tropicalis_Aquaporin5
A.thaliana_PIP15
A.thaliana_PIP12
A.thaliana_PIP11
A.thaliana_PIP13
A.thaliana_PIP14
A.thaliana_PIP26
A.thaliana_PIP25
A.thaliana_PIP23
A.thaliana_PIP22
A.thaliana_PIP24
A.thaliana_PIP28
A.thaliana_PIP27
0.7

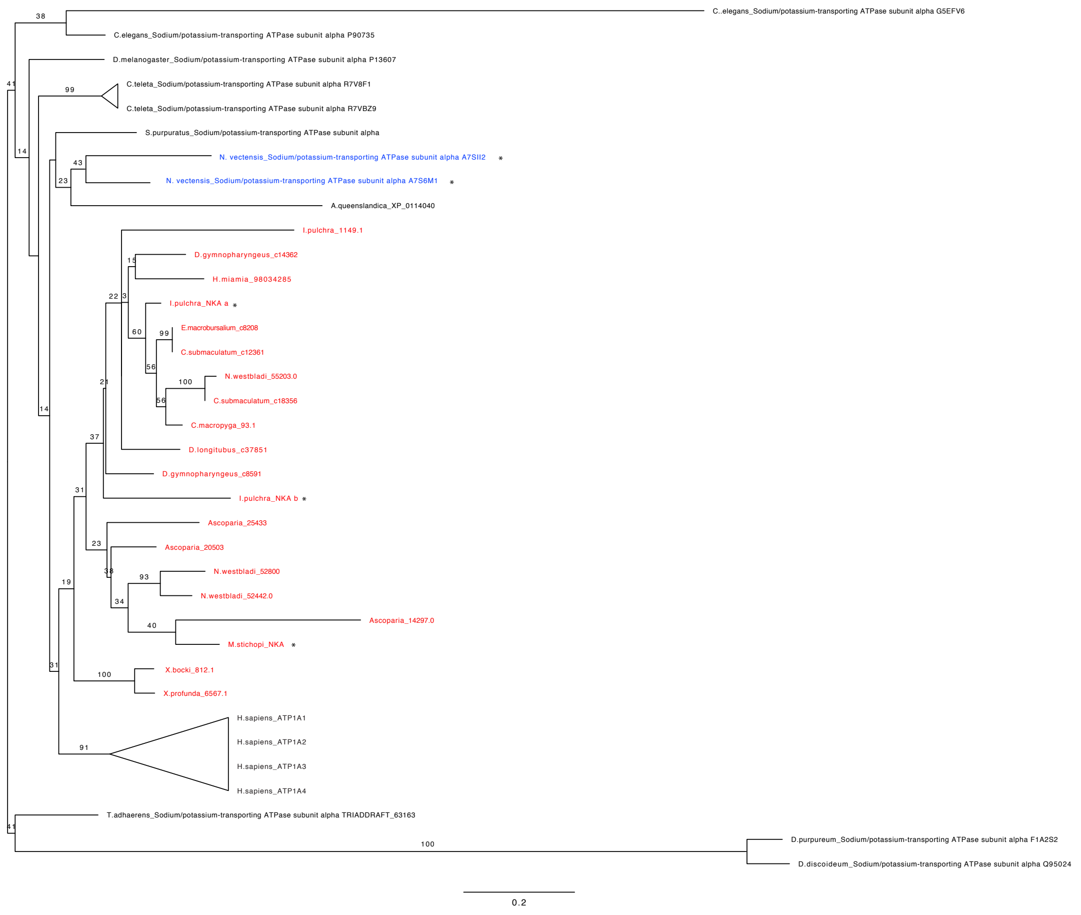
C..elegans_Sodium/potassium-transporting ATPase subunit alpha G5EFV6
C.elegans_Sodium/potassium-transporting ATPase subunit alpha P90735
D.melanogaster_Sodium/potassium-transporting ATPase subunit alpha P13607
C.teleta_Sodium/potassium-transporting ATPase subunit alpha R7V8F1
C.teleta_Sodium/potassium-transporting ATPase subunit alpha R7VBZ9
S.purpuratus_Sodium/potassium-transporting ATPase subunit alpha
N. vectensis_Sodium/potassium-transporting ATPase subunit alpha A7SII2 *
N. vectensis_Sodium/potassium-transporting ATPase subunit alpha A7S6M1 *
A.queenslandica_XP_0114040
I.pulchra_1149.1
D.gymnopharyngeus_c14362
H.miamia_98034285
I.pulchra_NKA a *
E.macrobursalium_c8208
C.submaculatum_c12361
N.westbladi_55203.0
C.submaculatum_c18356
C.macropyga_93.1
D.longitubus_c37851
D.gymnopharyngeus_c8591
I.pulchra_NKA b *
Ascoparia_25433
Ascoparia_20503
N.westbladi_52800
N.westbladi_52442.0
Ascoparia_14297.0
M.stichopi_NKA *
X.bocki_612.1
X.profunda_6567.1
H.sapiens_ATP1A1
H.sapiens_ATP1A2
H.sapiens_ATP1A3
H.sapiens_ATP1A4
T.adhaerens_Sodium/potassium-transporting ATPase subunit alpha TRIADDRAFT_63163
D.purpureum_Sodium/potassium-transporting ATPase subunit alpha F1A2S2
D.discoideum_Sodium/potassium-transporting ATPase subunit alpha Q95024
0.2

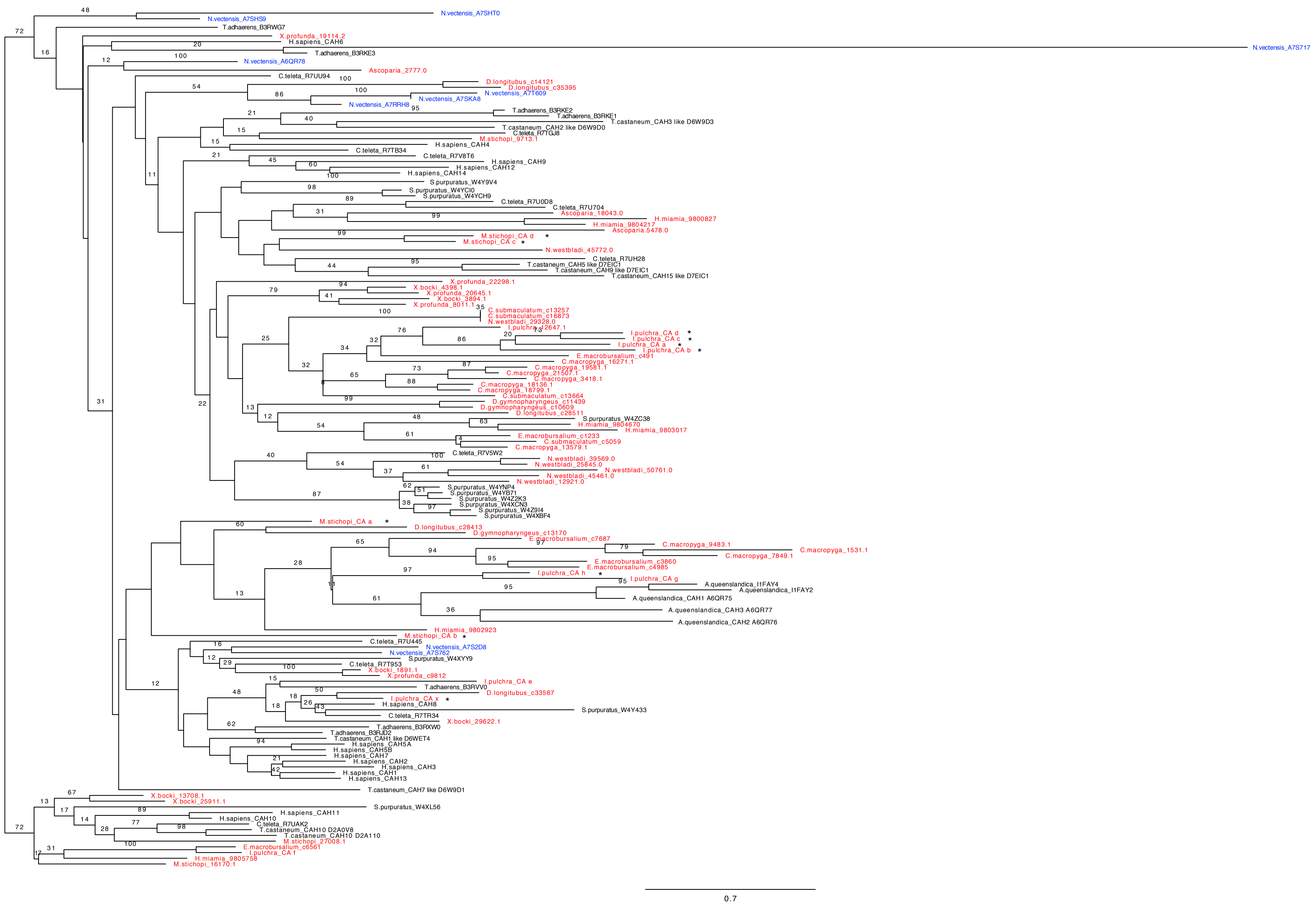

N.vectensis_A7SHT0
N.vectensis_A7SHS9
T.adhaerens_B3RWG7
X.profunda_19114.2
H.sapiens_CAH6
N.vectensis_A7S717
T.adhaerens_B3RKE3
N.vectensis_A6QR78
Ascoparia_2777.0
C.teleta_R7UU94
D.longitubus_c14121
D.longitubus_c35395
N.vectensis_A7T609
N.vectensis_A7SKA8
N.vectensis_A7RRH8
T.adhaerens_B3RKE2
T.adhaerens_B3RKE1
T.castaneum_CAH3 like D6W9D3
T.castaneum_CAH2 like D6W9D0
C.teleta_R7TGJ8
M.stichopi_9713.1
H.sapiens_CAH4
C.teleta_R7TB34
C.teleta_R7V8T6
H.sapiens_CAH9
H.sapiens_CAH12
H.sapiens_CAH14
S.purpuratus_W4Y9V4
S.purpuratus_W4YCI0
S.purpuratus_W4YCH9
C.teleta_R7U0D8
C.teleta_R7U704
Ascoparia_18043.0
H.miamia_9800827
H.miamia_9804217
Ascoparia_5478.0
M.stichopi_CA d *
M.stichopi_CA c *
N.westbladi_45772.0
C.teleta_R7UH28
T.castaneum_CAH5 like D7EIC1
T.castaneum_CAH9 like D7EIC1
T.castaneum_CAH15 like D7EIC1
X.profunda_22298.1
X.bocki_4398.1
X.profunda_20645.1
X.bocki_3894.1
X.profunda_8011.1
C.submaculatum_c13257
C.submaculatum_c16873
N.westbladi_29328.0
I.pulchra_12647.1
I.pulchra_CA d *
I.pulchra_CA c *
I.pulchra_CA a *
I.pulchra_CA b *
E.macrobursalium_c491
C.macropyga_16271.1
C.macropyga_19581.1
C.macropyga_21507.1
C.macropyga_3418.1
C.macropyga_18136.1
C.macropyga_18799.1
C.submaculatum_c13664
D.gymnopharyngeus_c11439
D.gymnopharyngeus_c10609
D.longitubus_c28511
S.purpuratus_W4ZC38
H.miamia_9804670
H.miamia_9803017
E.macrobursalium_c1233
C.submaculatum_c5059
C.macropyga_13579.1
C.teleta_R7V5W2
N.westbladi_39569.0
N.westbladi_25845.0
N.westbladi_50761.0
N.westbladi_45461.0
N.westbladi_12921.0
S.purpuratus_W4YNP4
S.purpuratus_W4YB71
S.purpuratus_W4Z2K3
S.purpuratus_W4XCN3
S.purpuratus_W4Z9I4
S.purpuratus_W4XBF4
M.stichopi_CA a *
D.longitubus_c28413
D.gymnopharyngeus_c13170
E.macrobursalium_c7687
C.macropyga_9483.1
C.macropyga_1531.1
C.macropyga_7849.1
E.macrobursalium_c3860
E.macrobursalium_c4985
I.pulchra_CA h *
I.pulchra_CA g
A.queenslandica_I1FAY4
A.queenslandica_I1FAY2
A.queenslandica_CAH1 A6QR75
A.queenslandica_CAH3 A6QR77
A.queenslandica_CAH2 A6QR76
H.miamia_9802923
M.stichopi_CA b *
C.teleta_R7U445
N.vectensis_A7S2D8
N.vectensis_A7S762
S.purpuratus_W4XYY9
C.teleta_R7T953
X.bocki_1891.1
X.profunda_c9812
I.pulchra_CA e
T.adhaerens_B3RVV0
D.longitubus_c33567
I.pulchra_CA x *
H.sapiens_CAH8
S.purpuratus_W4Y433
C.teleta_R7TR34
X.bocki_29622.1
T.adhaerens_B3RXW0
T.adhaerens_B3RJD2
T.castaneum_CAH1 like D6WET4
H.sapiens_CAH5A
H.sapiens_CAH5B
H.sapiens_CAH7
H.sapiens_CAH2
H.sapiens_CAH3
H.sapiens_CAH1
H.sapiens_CAH13
T.castaneum_CAH7 like D6W9D1
X.bocki_13708.1
X.bocki_25911.1
S.purpuratus_W4XL56
H.sapiens_CAH11
H.sapiens_CAH10
C.teleta_R7UAK2
T.castaneum_CAH10 D2A0V8
T.castaneum_CAH10 D2A110
M.stichopi_27008.1
E.macrobursalium_c6561
I.pulchra_CA f
H.miamia_9805758
M.stichopi_16170.1
0.7

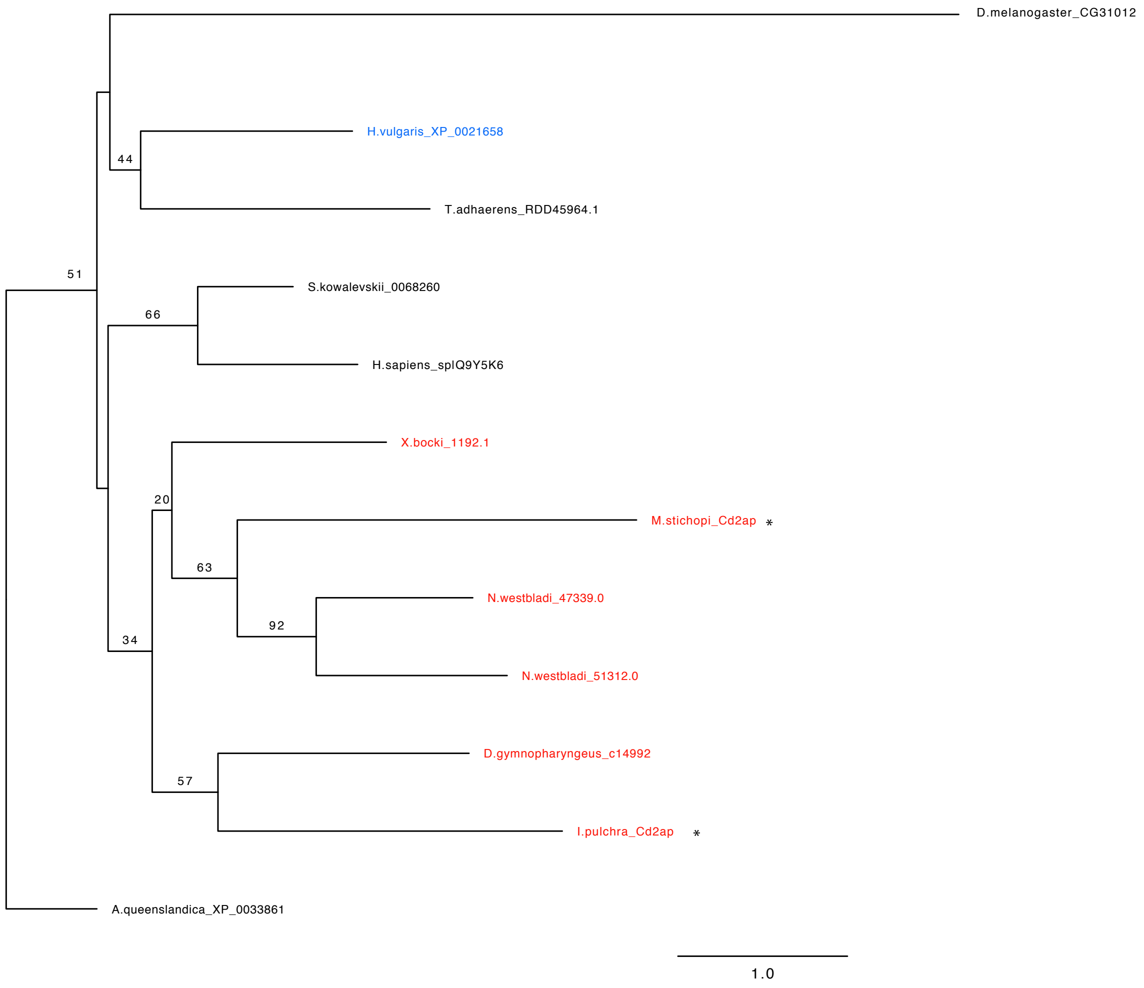
D.melanogaster_CG31012
H.vulgaris_XP_0021658
44
T.adhaerens_RDD45964.1
51
S.kowalevskii_0068260
66
H.sapiens_splQ9Y5K6
X.bocki_1192.1
20
M.stichopi_Cd2ap *
63
N.westbladi_47339.0
92
34
N.westbladi_51312.0
D.gymnopharyngeus_c14992
57
I.pulchra_Cd2ap *
A.queenslandica_XP_0033861
1.0

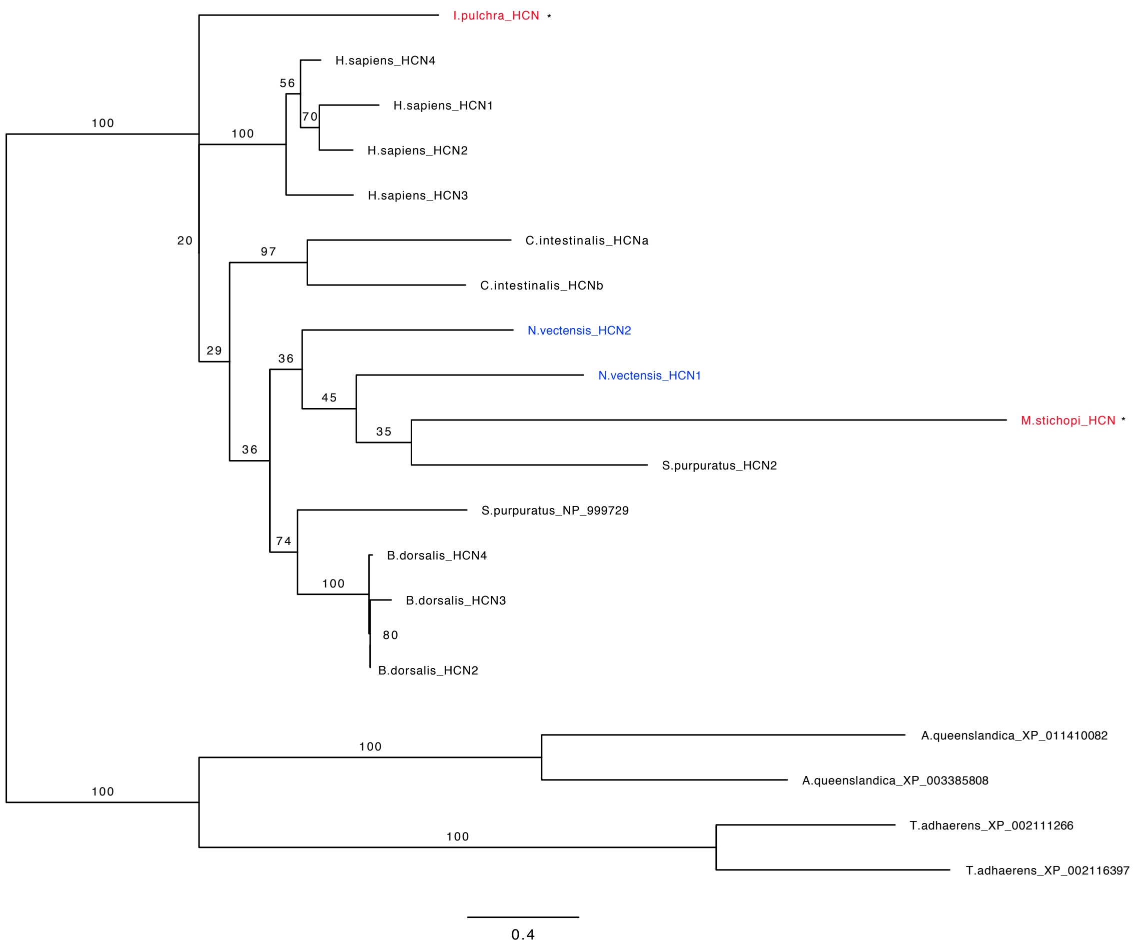
I.pulchra_HCN *
H.sapiens_HCN4
56
H.sapiens_HCN1
70
100
100
H.sapiens_HCN2
H.sapiens_HCN3
20
97
C.intestinalis_HCNa
C.intestinalis_HCNb
N.vectensis_HCN2
29
36
N.vectensis_HCN1
45
35
M.stichopi_HCN *
36
S.purpuratus_HCN2
S.purpuratus_NP_999729
74
B.dorsalis_HCN4
100
B.dorsalis_HCN3
80
B.dorsalis_HCN2
A.queenslandica_XP_011410082
100
A.queenslandica_XP_003385808
100
T.adhaerens_XP_002111266
100
T.adhaerens_XP_002116397
0.4

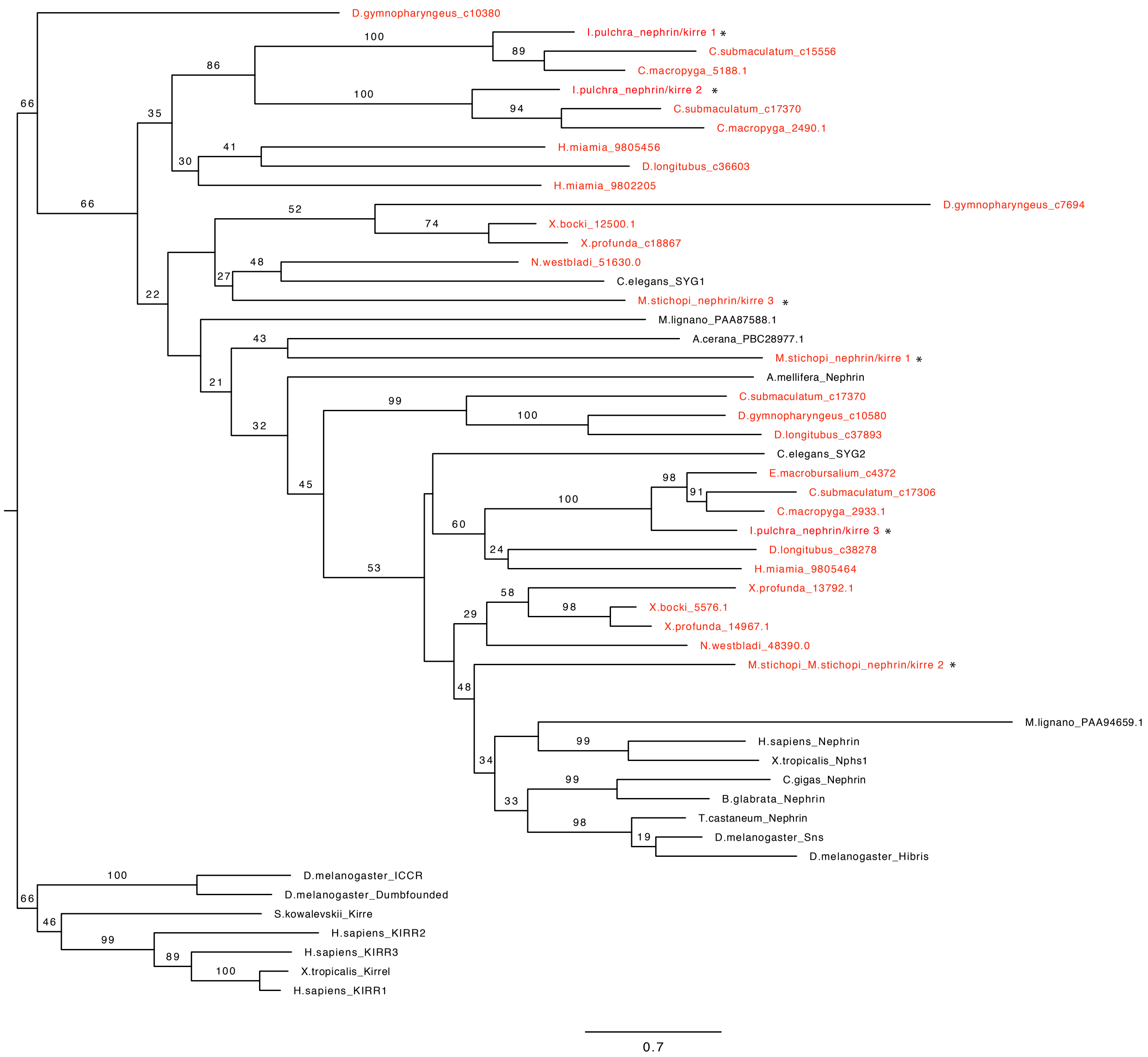
D.gymnopharyngeus_c10380
I.pulchra_nephrin/kirre 1 *
C.submaculatum_c15556
C.macropyga_5188.1
I.pulchra_nephrin/kirre 2 *
C.submaculatum_c17370
C.macropyga_2490.1
H.miamia_9805456
D.longitubus_c36603
H.miamia_9802205
D.gymnopharyngeus_c7694
X.bocki_12500.1
X.profunda_c18867
N.westbladi_51630.0
C.elegans_SYG1
M.stichopi_nephrin/kirre 3 *
M.lignano_PAA87588.1
A.cerana_PBC28977.1
M.stichopi_nephrin/kirre 1 *
A.mellifera_Nephrin
C.submaculatum_c17370
D.gymnopharyngeus_c10580
D.longitubus_c37893
C.elegans_SYG2
E.macrobursalium_c4372
C.submaculatum_c17306
C.macropyga_2933.1
I.pulchra_nephrin/kirre 3 *
D.longitubus_c38278
H.miamia_9805464
X.profunda_13792.1
X.bocki_5576.1
X.profunda_14967.1
N.westbladi_48390.0
M.stichopi_M.stichopi_nephrin/kirre 2 *
M.lignano_PAA94659.1
H.sapiens_Nephrin
X.tropicalis_Nphs1
C.gigas_Nephrin
B.glabrata_Nephrin
T.castaneum_Nephrin
D.melanogaster_Sns
D.melanogaster_Hibris
D.melanogaster_ICCR
D.melanogaster_Dumbfounded
S.kowalevskii_Kirre
H.sapiens_KIRR2
H.sapiens_KIRR3
X.tropicalis_Kirrel
H.sapiens_KIRR1
0.7

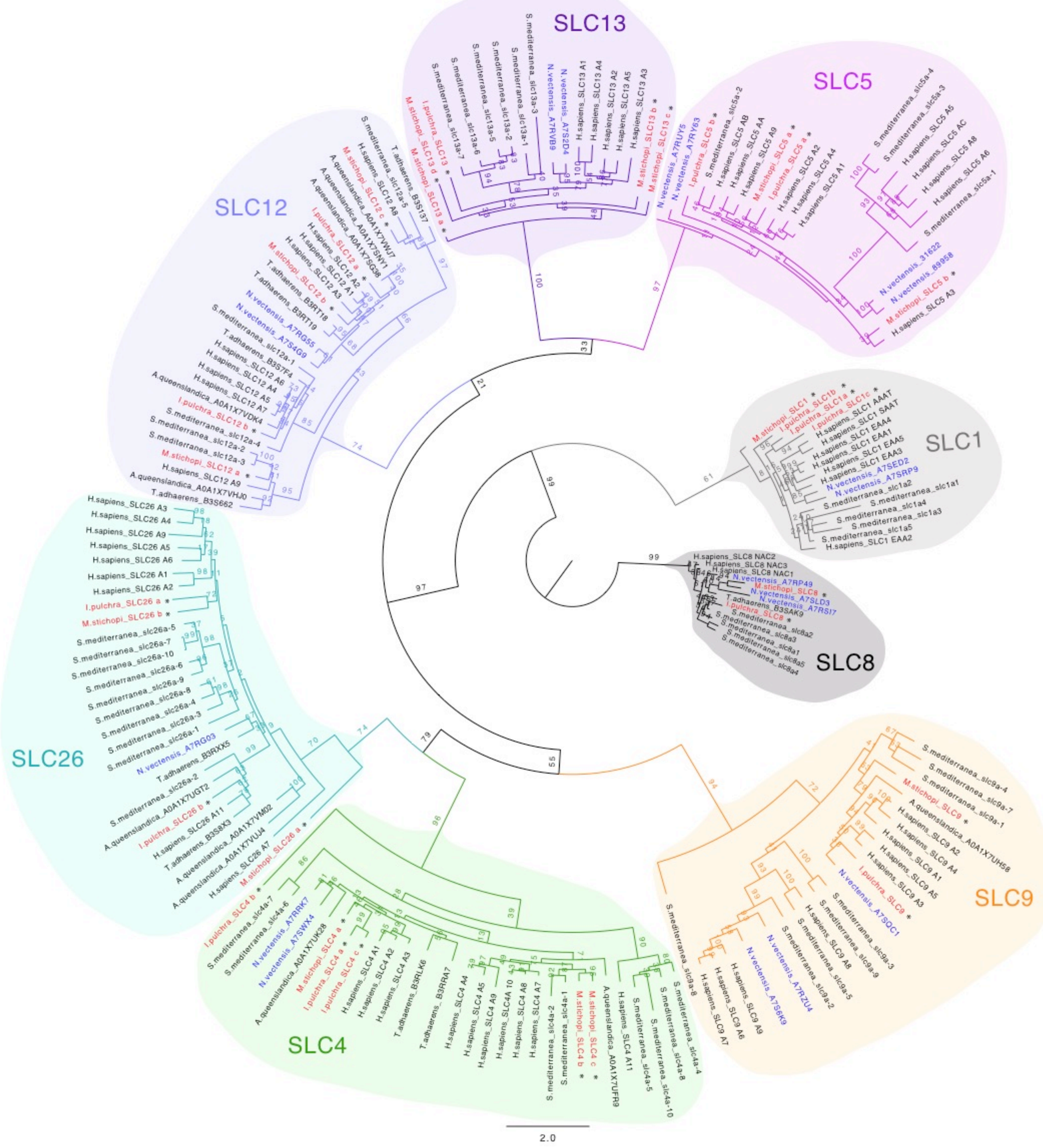
SLC13
SLC5
SLC12
SLC1
SLC8
SLC26
SLC9
SLC4
2.0

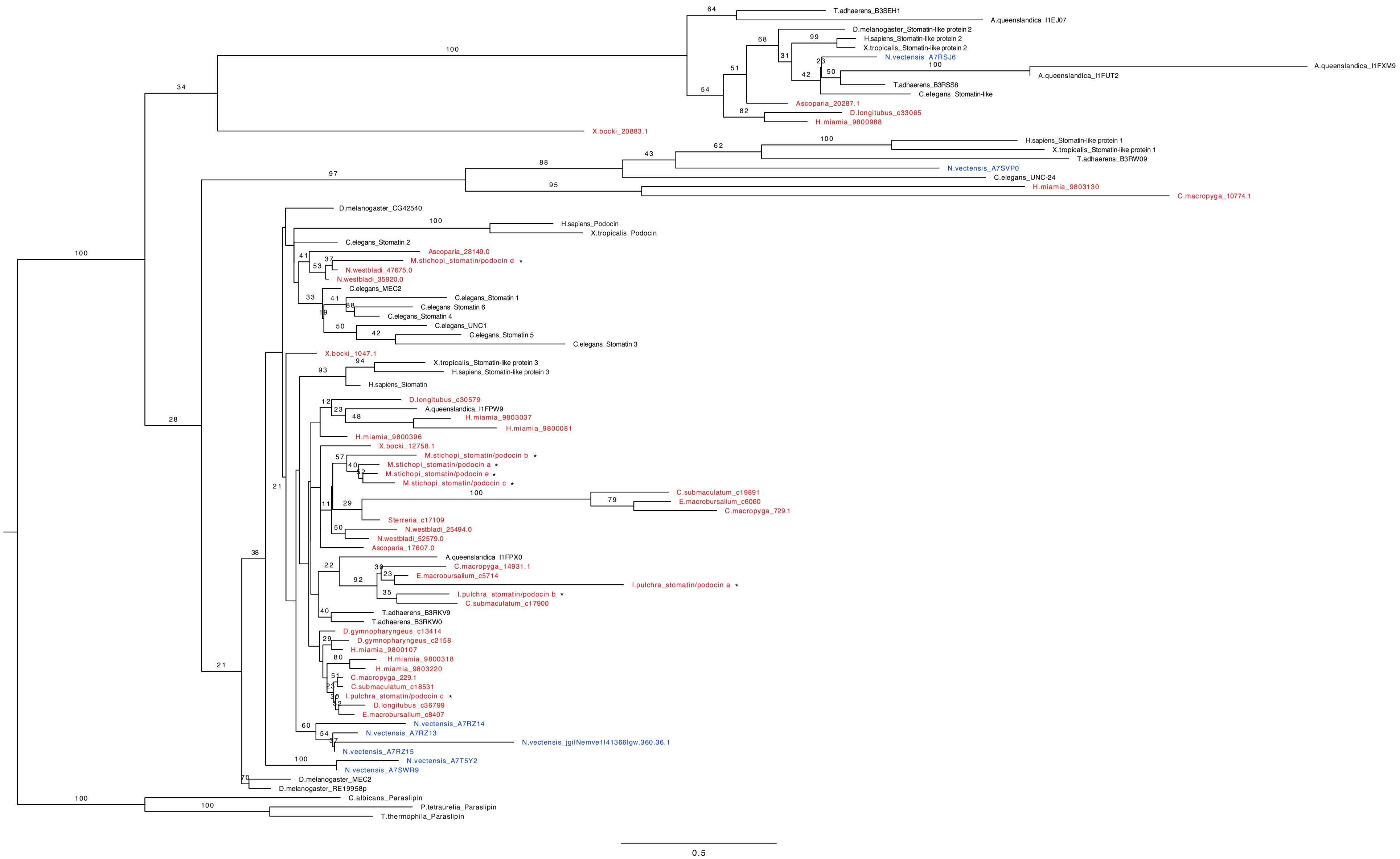
T.adhaerens_B3SEH1
A.queenslandica_I1EJ07
D.melanogaster_Stomatin-like protein 2
H.sapiens_Stomatin-like protein 2
X.tropicalis_Stomatin-like protein 2
N.vectensis_A7RSJ6
A.queenslandica_I1FXM9
A.queenslandica_I1FUT2
T.adhaerens_B3RSS8
C.elegans_Stomatin-like
Ascoparia_20287.1
D.longitubus_c33065
H.miamia_9800988
X.bocki_20883.1
H.sapiens_Stomatin-like protein 1
X.tropicalis_Stomatin-like protein 1
T.adhaerens_B3RW09
N.vectensis_A7SVP0
C.elegans_UNC-24
H.miamia_9803130
C.macropyga_10774.1
D.melanogaster_CG42540
H.sapiens_Podocin
X.tropicalis_Podocin
C.elegans_Stomatin 2
Ascoparia_28149.0
M.stichopi_stomatin/podocin d *
N.westbladi_47675.0
N.westbladi_35920.0
C.elegans_MEC2
C.elegans_Stomatin 1
C.elegans_Stomatin 6
C.elegans_Stomatin 4
C.elegans_UNC1
C.elegans_Stomatin 5
C.elegans_Stomatin 3
X.bocki_1047.1
X.tropicalis_Stomatin-like protein 3
H.sapiens_Stomatin-like protein 3
H.sapiens_Stomatin
D.longitubus_c30579
A.queenslandica_I1FPW9
H.miamia_9803037
H.miamia_9800081
H.miamia_9800396
X.bocki_12758.1
M.stichopi_stomatin/podocin b *
M.stichopi_stomatin/podocin a *
M.stichopi_stomatin/podocin e *
M.stichopi_stomatin/podocin c *
C.submaculatum_c19891
E.macrobursalium_c6060
C.macropyga_729.1
Sterreria_c17109
N.westbladi_25494.0
N.westbladi_52579.0
Ascoparia_17607.0
A.queenslandica_I1FPX0
C.macropyga_14931.1
E.macrobursalium_c5714
I.pulchra_stomatin/podocin a *
I.pulchra_stomatin/podocin b *
C.submaculatum_c17900
T.adhaerens_B3RKV9
T.adhaerens_B3RKW0
D.gymnopharyngeus_c13414
D.gymnopharyngeus_c2158
H.miamia_9800107
H.miamia_9800318
H.miamia_9803220
C.macropyga_229.1
C.submaculatum_c18531
I.pulchra_stomatin/podocin c *
D.longitubus_c36799
E.macrobursalium_c8407
N.vectensis_A7RZ14
N.vectensis_A7RZ13
N.vectensis_jgi|Nemve1|41366|gw.360.36.1
N.vectensis_A7RZ15
N.vectensis_A7T5Y2
N.vectensis_A7SWR9
D.melanogaster_MEC2
D.melanogaster_RE19958p
C.albicans_Paraslipin
P.tetraurelia_Paraslipin
T.thermophila_Paraslipin
0.5

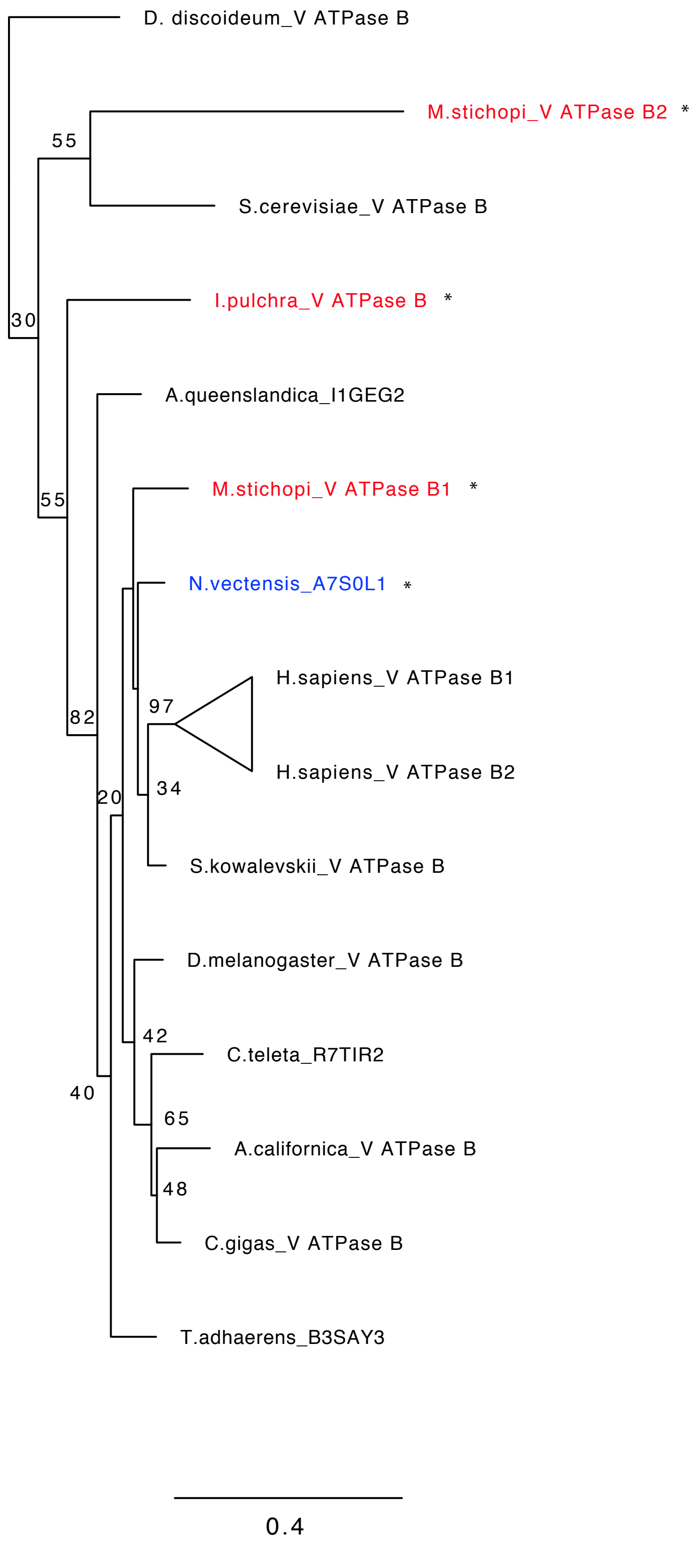
D. discoideum_V ATPase B
M.stichopi_V ATPase B2 *
55
S.cerevisiae_V ATPase B
I.pulchra_V ATPase B *
30
A.queenslandica_I1GEG2
M.stichopi_V ATPase B1 *
55
N.vectensis_A7S0L1 *
H.sapiens_V ATPase B1
97
82
H.sapiens_V ATPase B2
34
20
S.kowalevskii_V ATPase B
D.melanogaster_V ATPase B
42
C.teleta_R7TIR2
40
65
A.californica_V ATPase B
48
C.gigas_V ATPase B
T.adhaerens_B3SAY3
0.4

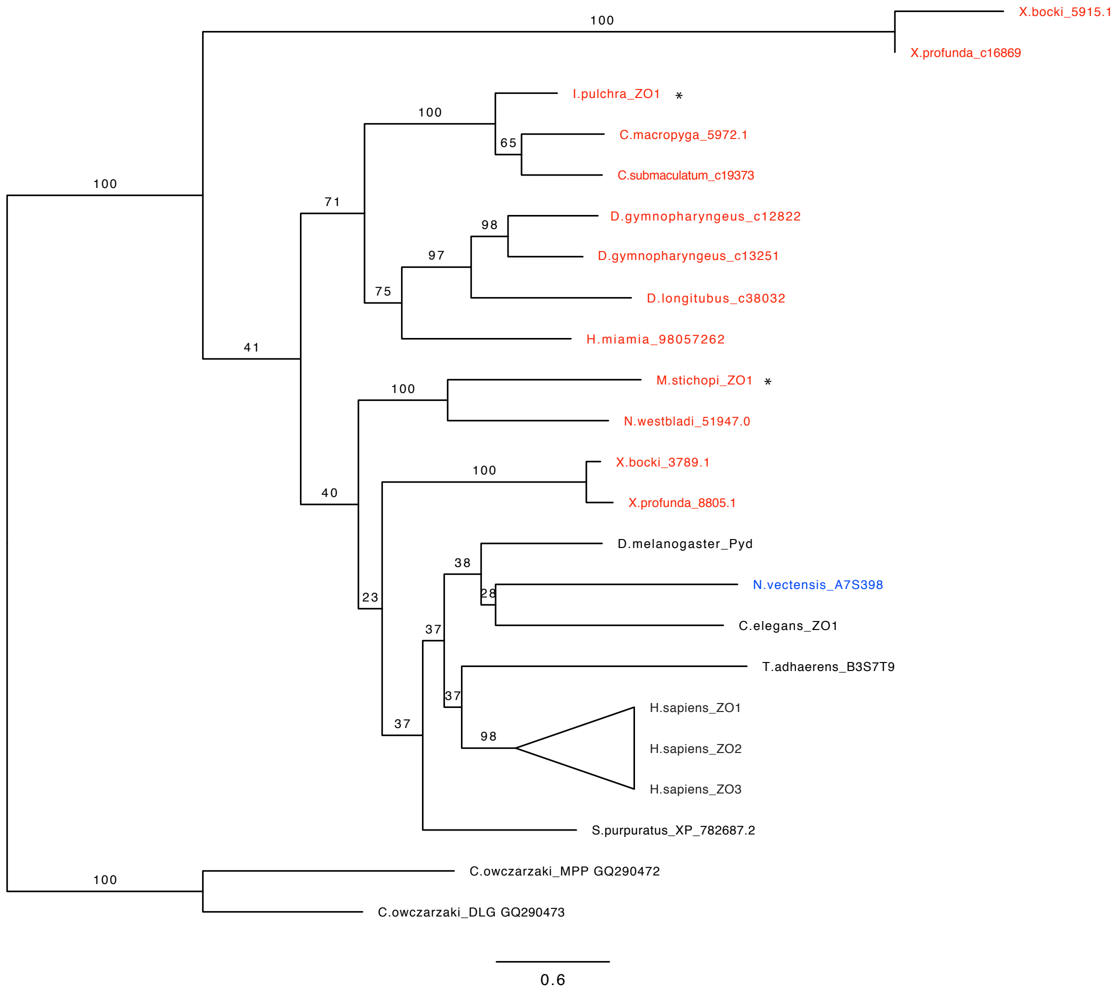
X.bocki_5915.1
X.profunda_c16869
I.pulchra_ZO1 *
C.macropyga_5972.1
C.submaculatum_c19373
D.gymnopharyngeus_c12822
D.gymnopharyngeus_c13251
D.longitubus_c38032
H.miamia_98057262
M.stichopi_ZO1 *
N.westbladi_51947.0
X.bocki_3789.1
X.profunda_8805.1
D.melanogaster_Pyd
N.vectensis_A7S398
C.elegans_ZO1
T.adhaerens_B3S7T9
H.sapiens_ZO1
H.sapiens_ZO2
H.sapiens_ZO3
S.purpuratus_XP_782687.2
C.owczarzaki_MPP GQ290472
C.owczarzaki_DLG GQ290473
100
100
100
71
65
98
97
75
41
100
40
100
38
28
23
37
37
98
37
100
0.6

**Supplementary Figure 2.** Orthology analysis. a-p, Putative orthologous sequences of genes of interest were identified by tBLASTx search against the transcriptome (SRR2681926) of *Isodiametra pulchra*, the transcriptome (SRR2681155) and draft genome of *Meara stichopi* and the genome of *Nematostella vectensis* (http://genome.jgi.doe.gov). Additional transcriptomes of Xenacoelomorpha species investigated were: *Childia submaculatum* (Acoela) (SRX1534054), *Convolutriloba macropyga* (Acoela) (SRX1343815), *Diopisthoporus gymnopharyngeus* (Acoela) (SRX1534055), *Diopisthoporus longitubus* (Acoela) (SRX1534056), *Eumecynostomum macrobursalium* (Acoela) (SRX1534057), *Hofstenia miamia* (Acoela) (PRJNA241459), *Ascoparia sp.* (Nemertodermatida) (SRX1343822), *Nemertoderma westbladi* (Nemertodermatida) (SRX1343819), *Sterreria sp.* (Nemertodermatida) (SRX1343821), *Xenoturbella bocki* (Xenoturbella) (SRX1343818) and *Xenoturbella profunda* (Xenoturbella) (SRP064117). Bayesian phylogenetic analysis is supporting orthology for genes investigated in this study. Red color refers to Xenacoelomorpha taxa and blue color refers to *N. vectensis*. Bootstrap values are shown when equal or above 20%. Branches crossed by a double slash were shortened to make figures more compact. Names of genes or proteins, if available, follow the name of organism(s); otherwise the accession number is written. Asterisks indicate genes with a spatial expression by WMISH.

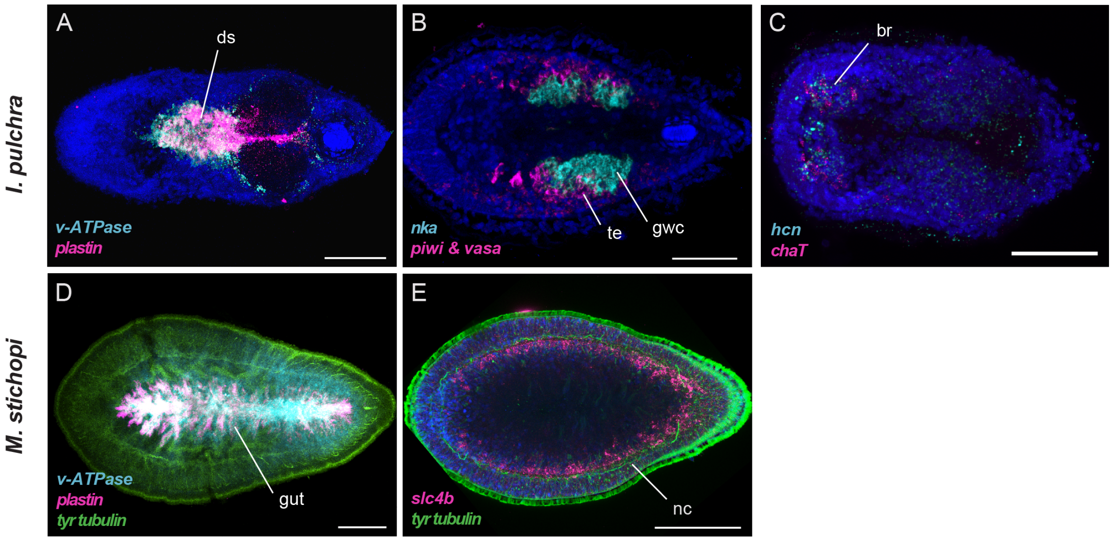

**Supplementary Figure 3.** Co-expression analysis of excretion-related components and molecular markers of digestive (*plastin*), nervous (*tyrosinated tubulin*) and reproductive systems (*piwi, vasa*) by double FISH in *I. pulchra* and *M. stichopi*. Every picture is a full projection of merged confocal stacks. Nuclei are stained blue with 4',6-diamidino-2-phenylindole (DAPI). Anterior is to the left. Abbreviations: br, brain; ds, digestive syncytium; gwc, gut-wrapping cells; nc, nerve cords; te, testis; Scale bars are 50 µm for *I. pulchra* and 100 µm for *M. stichopi*.

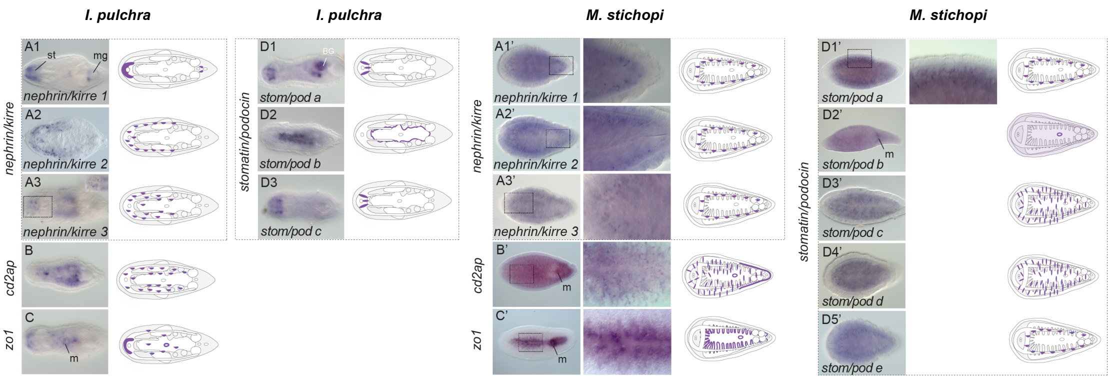

**Supplementary Figure 4.** Whole mount in situ hybridization (WMISH) of genes encoding the slit diaphragm components that are related to ultrafiltration *nephrin/kirre, cd2ap, zo1* and *stomatin/podocin*. The inset in panel A3 shows a different focal plane. The columns next to *M. stichopi* panels show higher magnifications of the indicated domains. Illustrations with colored gene expression correspond to the animals shown in the previous column. Anterior is to the left. The depicted expression patterns are for guidance and not necessarily represent exact expression domains. Drawings are not to scale. Abbreviations: mo, mouth; mg, male gonopore; st, statocyst.

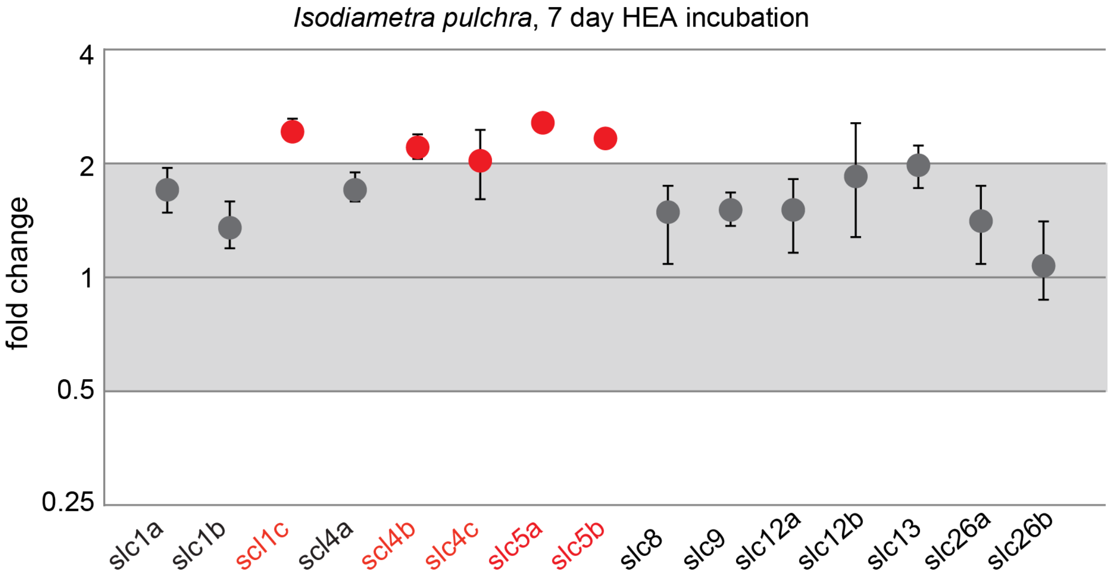

**Supplementary Figure 5.** Quantitative relative expression of *slcs* after 7d exposure in HEA (1 mM $NH_4Cl$) in *I. pulchra*. Each circle represents the average of four independent measurements of three independent biological experiments. One fold change represents no change; ≥ 2 indicates increased expression level significantly; ≤ 0.5 indicates decreased expression level significantly (red labels).

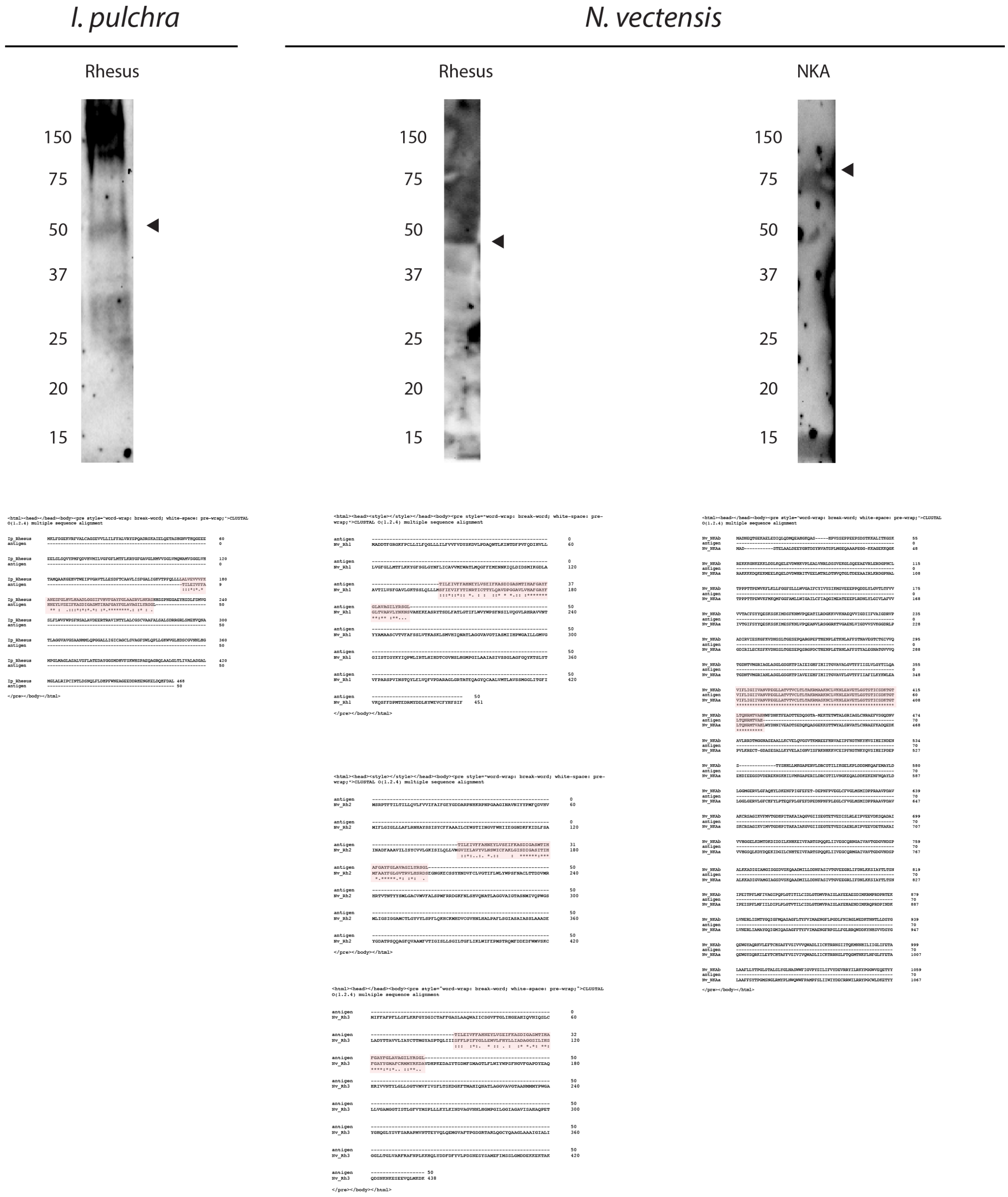

**Supplementary Figure 6.** Western blots of Rhesus in *I. pulchra and Rhesus and NKA in N. vectensis*. Below each blot, the sequence alignment of the endogenous protein and the antigen is provided, highlighted in pink.

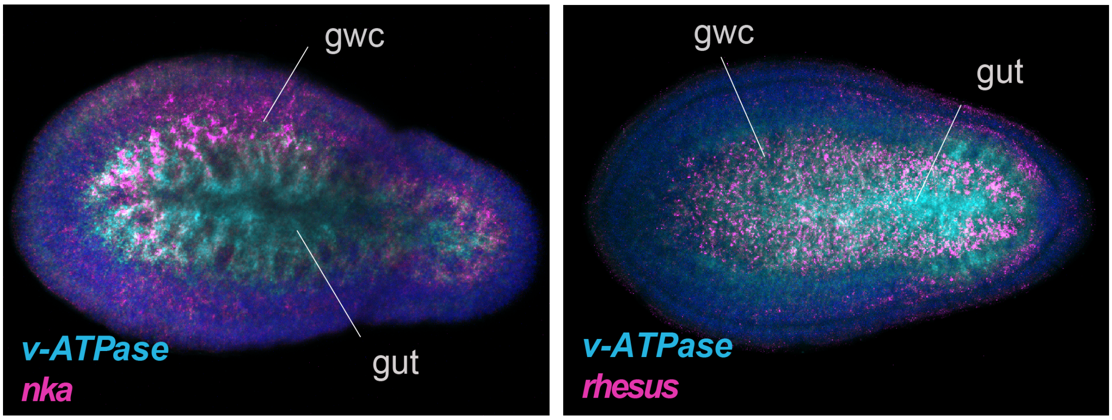

**Supplementary Figure 7.** Double fluorescent WMISH of *v-ATPase* and *nka,* and *v-ATPase* and *rhesus* in *M. stichopi.* Every picture is a full projection of merged confocal stacks. Nuclei are stained blue with 4',6-diamidino-2-phenylindole (DAPI). Anterior is to the left. Abbreviations: dlr, distal lateral rows; ds, digestive syncytium; gwc, gut-wrapping cells.

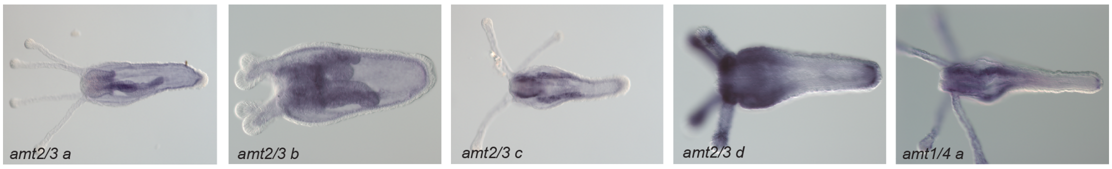

**Supplementary Figure 8.** Gene expression of ammonia transporters in *N. vectensis*. Whole mount in situ hybridization (WMISH) of *amt2/3a, amt2/3b, amt2/3c, amt2/3d* and *amt1/4a* in juvenile polyps. Anterior is to the left.

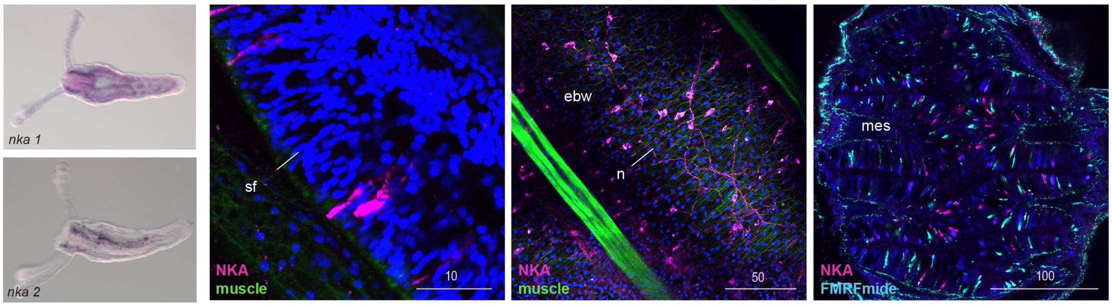

**Supplementary Figure 9.** Gene expression and protein localization of NKA in *N. vectensis*. WMISH of *nka1* and *nka2* in juvenile polyps. Anterior is to the left. Protein localization of NKA in *N. vectensis* juvenile polyps. The muscle filaments are labeled green with phalloidin and the nervous system is stained cyan with tyrosinated tubulin. Every picture is a full projection of merged confocal stacks. Nuclei are stained blue with 4',6-diamidino-2-phenylindole (DAPI). The regions shown are indicated with dashed boxes in the illustrated animal. Abbreviations: ebw, endodermal body wall; mes, mesenteries; sf, septal filaments; n, neurons.

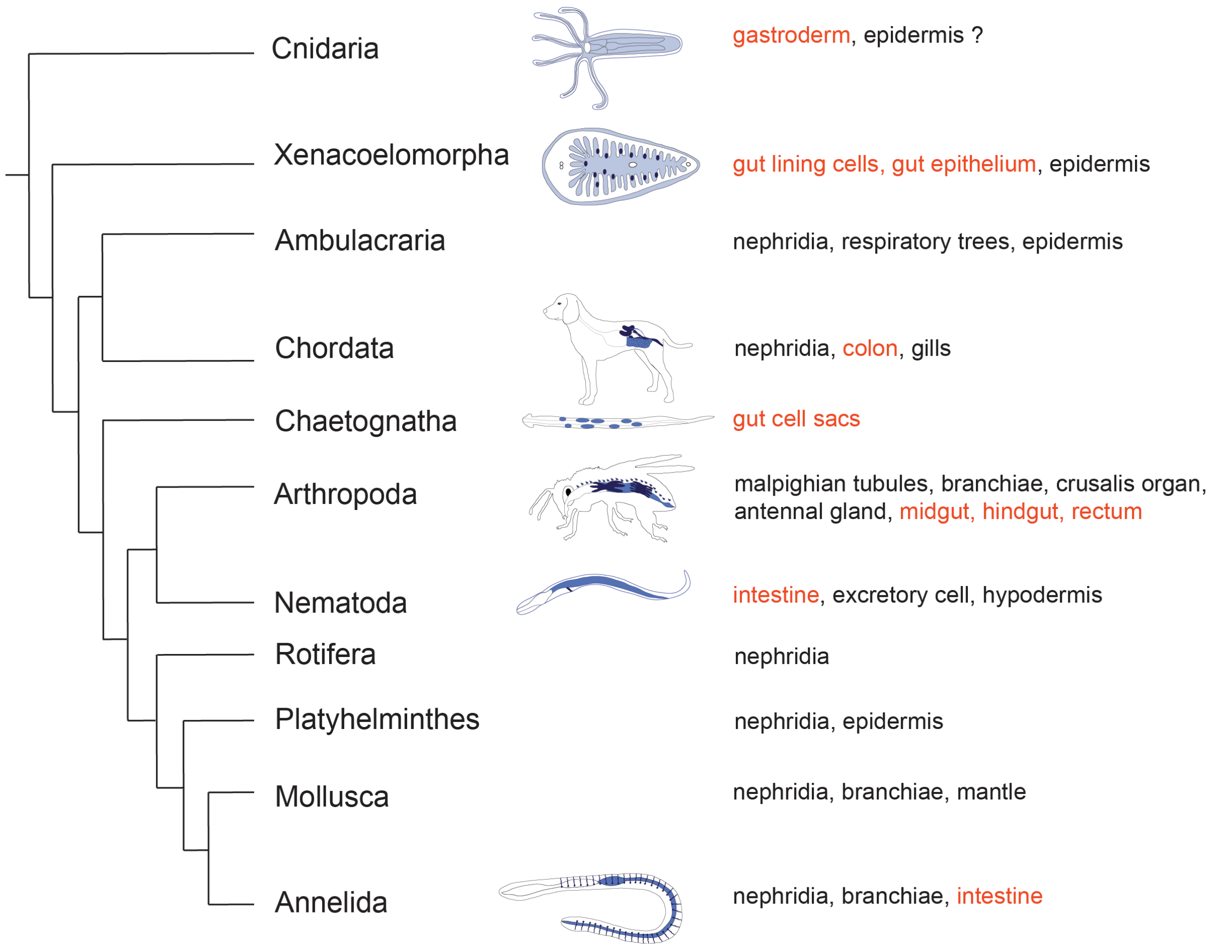

**Supplementary Figure 10.** Phylogenetic tree showing the relationship of animal groups where the role of gut in excretion has been demonstrated or proposed.

| | *Nephrin/Kirre* | *Cd2ap/Cin8* | *Zo1* | *Slipins* | *Aquaporins* |
|---|---|---|---|---|---|
| **Arthropoda** | nephrocytes (Weavers et al., 2009), eye (Ramos et al., 1993), muscle (Strunkelnberg et al., 2001), CNS (Dworak et al., 2001) | nephrocytes (Weavers et al., 2009), eye (Johnson et al., 2008), germline (Eikenes et al., 2013), epithelial sheet (Yasin et al., 2016) | nephrocytes (Weavers et al., 2009), eye (Seppa et al., 2008), germline, adult wing (Djiane et al., 2011), epithelia (Wei and Ellis, 2001) | nephrocytes (Weavers et al., 2009) | malpighian tubules, fat body, pharynx, epidermis, ovaries, gut (sum. in (Campbell et al., 2008)) |
| **Platyhelminthes** | terminal cells (Thi-Kim Vu et al., 2015), stem cells (Nakamura et al., 2014) | ? | ? | ? | ubiquitous, brain (Hwang et al., 2015) |
| **Mollusca** | rhogocytes (Kokkinopoulou et al., 2014) | ? | ? | ? | ? |
| **Annelida** | ? | ? | ? | ? | ? |
| **Nematoda** | neural axons, vulval muscle (Wanner et al., 2011) | ? | ? | melanosensory neurons (Zhang et al., 2004) | excretory cell, neurons, intestine, pharynx, muscle, hypodermis (Huang et al., 2007) |
| **Acoela** | neurons, brain, gonadal domain | neurons | brain, neurons, mouth | digestive syncytium, brain | brain, digestive syncytium, gut wrapping cells, parenchyme |
| **Nemertodermatida** | neurons | subepidermis, anterior domain, posterior lateral rows of cells, mouth | gut epithelium, mouth | neurons, mouth, subepidermis | subepidermal cells, epidermis gut wrapping cells, brain |
| **Cnidaria** | absent | ? | ectoderm(Fei et al., 2000) | ? | ? |
| **Vertebrata** | podocytes, muscles, CNS, testis, pangreatic cells, cardiovascular development, spleen, lymph node (sum. in (Li and He, 2015)) | podocytes (Shih et al., 1999), brain, heart, pancreas, salivary gland (Li et al., 2000) | podocytes (Itoh et al., 2014), eye (Sugrue and Zieske, 1997), epithelial sheet, notochord, vascular development, neural tube (Bauer et al., 2010) | podocytes, erythrocyte membranes, sensory neurons, skin, dorsal root ganglia (sum. in (Lapatsina et al., 2012)) | blood, kidney, brain, testis, oocyte oocyte eyes, pancreas, muscle, lung, digestive tract (sum. in (Ishibashi et al., 2011)), |
| **Eubacteria/Archaea** | absent | absent | absent (de Mendoza et al., 2010) | ? (Green and Young, 2008) | osmoregulation (Abascal et al., 2014) |
| **Filasterea/ Choanoflagellata** | absent | ? | ? (de Mendoza et al., 2010) | ? ((Green and Young, 2008), this study) | ? (Abascal et al., 2014) |
| **Holomycota** | absent | absent | absent (de Mendoza et al., 2010) | ? (Green and Young, 2008) | spores/ endoplasmic reticulum, plasma membrane (sum. in (Gomes et al., 2009)) |

| | *Rh/AMTs* | *$Na^+/K^+$ ATPase* | *CA a (cytosol & membrane)* | *V-ATPase A/B* |
|---|---|---|---|---|
| **Arthropoda** | branchiae (Martin et al., 2011), brain, salivary glands (Chintapalli et al., 2007), head, thorax (Wu et al., 2010), malpighian tubules, ganglia, trachea, fat body, head, midgut, hindgut, eye (Weihrauch, 2006; Weihrauch et al., 2009) | malpighian tubules, rectum, midgut, hindgut, anal papillae (Patrick et al., 2006), Johnston's organ of hearing (Roy et al., 2013), Crusalis organs (Gerber et al., 2016) testis, eye, brain (Yasuhara et al., 2000), (Chintapalli et al., 2007), branchiae (Weihrauch, 2006; Weihrauch et al., 1999) | midgut and hindgut (sum. in (Linser et al., 2009)), malpighian tubules, branchiae (sum. in (Weihrauch and O'Donnell, 2015)), epidermis (Calhoun and Zou, 2016) | malpighian tubules, fat body, midgut, hindgut, ganglia, trachea, antennal gland (Blaesse et al., 2010; Tsai and Lin, 2014), anal papilla (Patrick et al., 2006), branchiae (sum. in (Weihrauch and O'Donnell, 2015)), Crusalis organs (Gerber et al., 2016), neurons, (Williamson et al., 2010a; Williamson et al., 2010b) brain, testis, ovarioles, duct (Allan et al., 2005) |
| **Platyzoa** | epidermis (Weihrauch et al., 2012), duct of protonephridia (Thi-Kim Vu et al., 2015) | epidermis (Weihrauch et al., 2012), protonephridia (Scimone et al., 2011) | epidermis (Weihrauch et al., 2012), protonephridia (Scimone et al., 2011) | epidermis (Weihrauch et al., 2012) |
| **Mollusca** | epidermis (Hu et al., 2013), branchiae (Hu et al., 2014) | foot muscle, hepatopancreas (Ramnanan and Storey, 2006), branchiae, pancreatic appendages, nerves, nephridia (Hu et al., 2010; Hu et al., 2017) mantle (Li et al., 2016) | mantle (Le Roy et al., 2014), branchiae (Hu et al., 2011) | branchiae (Hu et al., 2011), mantle (Li et al., 2016) |
| **Annelida** | epidermis (Quijada-Rodriguez et al., 2015), branchiae (Thiel et al., 2016) | branchiae (Thiel et al., 2016), epidermis (sum. in (Schnizler et al., 2002)) | epidermal glands (Le Roy et al., 2014), branchiae (Thiel et al., 2016), epidermis (Quijada-Rodriguez et al., 2015) | branchiae (Thiel et al., 2016), epidermis (Quijada-Rodriguez et al., 2015) |
| **Nematoda** | hypodermis | hypodermis (Adlimoghaddam et al., 2015) | intestine (Adlimoghaddam et al., 2015) | hypodermis, intestine, neurons (Oka et al., 2001) |
| **Acoela** | gut wrapping cells, neurons, brain, parenchyme, mouth, epidermis | gut wrapping cells | mouth, parenchyme, neurons brain, posterior cells, gut wrapping cells | digestive syncytium |
| **Nemertodermatida** | gut wrapping cells, brain, epidermis | gut epithelium | gut wrapping cells, gut epithelium, epidermis, neurons | gut epithelium, subepidermis, nerve cords |
| **Cnidaria** | tentacular ectodermal cells, gastroderm, mesenteries, pharynx | pharyngeal endoderm, mesenteries, neurons, septal filament, pharynx | tentacular ectoderm, mesenteries, gastroderm (this study), oral and aboral endoderm, calicoblastic cells, ectoderm (sum. in (Bertucci et al., 2013)) | gastroderm, mesenteries, pharynx, septal filaments |
| **Vertebrata** | blood, esophagael epithelia, brain, kidney duct, liver, gastrointestinal tract testis (sum. in (Huang and Ye, 2010)), gills (Nakada et al., 2007; Nawata et al., 2007), muscle (Takeda and Takemasa, 2015), skin (Cruz et al., 2013) | kidney tubule and duct (Garvin et al., 1985; Wall and Koger, 1994), skin (Cruz et al., 2013), intestine, gut, eye, heart, testis, liver, brain (Rahman et al., 2015; Worrell et al., 2008), gills (Li et al., 2014; Mallery, 1983) | brain, bones, muscle, intestine, kidney intercalated cells, tubule, duct, eye, erythrocytes, liver, urinary bladder, teeth, inner ear (sum. in (McKenna and Frost, 2014)), gills (Nawata et al., 2007; Weihrauch et al., 2009) | pancreas (Sun-Wada et al., 2006), kidney tubule and duct (Wagner et al., 2004), eye (Jouhou et al., 2007; Kawamura et al., 2010), osteoclast (Toyomura et al., 2003), skin (Shih et al., 2008), testis (Pietrement et al., 2006), brain (Moriyama and Futai, 1990), gills (Nawata et al., 2007; Weihrauch et al., 2009) |
| **Eubacteria/Archaea** | $NH_3$ transport (Huang and Peng, 2005) | osmoregulation (Chan et al., 2010; Saez et al., 2009) | photosynthesis, $CO_2$ fixation (Smith and Ferry, 2000) | proton pump (Gogarten et al., 1992) |
| **Filasterea/ Choanoflagellata** | $NH_3$ transporter (King et al., 2008) | osmoregulation (Chan et al., 2010; Saez et al., 2009) | ? | ? |
| **Holomycota** | $NH_3$ transport (Marini et al., 2000; Marini et al., 1997) | osmoregulation (Benito et al., 2002) | $CO_2$ sensing system (Elleuche and Poggeler, 2010) | vacuole, Golgi/endosome (Kawasaki-Nishi et al., 2001) |

| | *SLC1* | *SLC4* | *SLC5* | *SLC8* |
|---|---|---|---|---|
| **Arthropoda** | eye, central nervous system (Besson et al., 1999; Umesh et al., 2003) (Kucharski et al., 2000), glia cells (Soustelle et al., 2002) | alimentary canal (Linser et al., 2012), malpighian tubules (Yamahiro et al., 2008), heart (Perrin et al., 2004), neurons (Romero et al., 2000) | malpighian tubules (Stergiopoulos et al., 2009), intestine (Obi et al., 2011), hepatopancreas (Ahearn et al., 1985), glia cells (Freeman et al., 2003) | retina, brain (Schwarz and Benzer, 1997) |
| **Platyzoa** | protonephridia (Thi-Kim Vu et al., 2015) | protonephridia (Thi-Kim Vu et al., 2015) | protonephridia (Thi-Kim Vu et al., 2015) | protonephridia (Thi-Kim Vu et al., 2015) |
| **Mollusca** | nervous system (Hatakeyama et al., 2010) | branchiae ((Hu et al., 2011), heart, neurons, optic lobe, testis (Piermarini et al., 2007) | hepatopancreas (Blaya et al., 1998), gills, mantle edge (Hanquet et al., 2011) | muscle, neurons (Rasgado-Flores et al., 1996) (Rosenthal and Gilly, 1993) |
| **Annelida** | central nervous system, glia cells (Hirth and Deitmer, 2006) | mesoderm, epidermis muscle, mesenchyme (Miyamoto et al., 2017), neurons (Munsch and Deitmer, 1994) | bacteriocytes (Miyamoto et al., 2017) | ? |
| **Nematoda** | muscle, pharynx, head neurons, excretory canal (Mano et al., 2007) | neurons, hypodermis, muscle (Sherman et al., 2005) | neurons (Okuda et al., 2000) | neurons, muscle, intestine (Sharma et al., 2013) |
| **Acoela** | brain, gonadal domain | parenchyme, brain anterior tip | parenchyme, brain gonadal domain | anterior tip |
| **Nemertodermatida** | neurons | nerve cords, mouth, female gonads, subepidermis, gut wrapping cells | posterior lateral rows of cells, subepidermis | gut epithelium, mouth, posterior lateral rows of cells |
| **Cnidaria** | ? | oral/aboral endoderm and ectoderm, calicoblastic ectoderm (Zoccola et al., 2015) | ? | ? |
| **Vertebrata** | kidney, intestine, brain (sum. in (Grewer et al., 2014)) | brain, kidney duct and tubule, ovary, testis, salivary gland, erythrocytes, heart, intestine, heart, pancreas, liver, smooth muscle, lung, retina, spleen (sum. in (Romero et al., 2013)), gill (Lee et al., 2011) | kidney, intestine, salivary glands, brain, retina, muscle, heart, uterus, testis, lung, placenta, liver (sum. in (Wright, 2013)) | brain, heart, kidney, pancreas, muscle, liver (sum. in (Khananshvili, 2013) |
| **Eubacteria/Archaea** | glytamate transport (Yernool et al., 2004) | $NO_2$/ $NO_3$ exchange, $Cl^-$ influx (Parker and Boron, 2013) | absent (Höglund et al., 2011) | $Na^+$/ $Ca^{2+}$ exchange (Liao et al., 2012) |
| **Filasterea/ Choanoflagellata** | absent ((Höglund et al., 2011), this study) | ? | absent ((Höglund et al., 2011), this study) | ((Cai and Clapham, 2012), this study) |
| **Holomycota** | absent (Höglund et al., 2011) | borate transport (Jennings et al 2007) (Parker and Boron, 2013) | absent (Höglund et al., 2011) | absent (Cai and Lytton, 2004) (Pittman and Hirschi, 2016) |

| | ***SLC9*** | ***SLC12*** | ***SLC13*** | ***SLC26*** |
|---|---|---|---|---|
| **Arthropoda** | midgut (Blaesse et al., 2010), gill (Weihrauch et al., 1998), malpighian tubules (Kang'ethe et al., 2007) | salivary gland, ventral nerve cord, gut, anal pad (Sun et al., 2010), malpighian tubules (Piermarini et al., 2011), glia cells (Leiserson et al., 2011) | fat body, oenocytes, midgut (Inoue et al., 2002) | gut epithelial cells, malpighian tubules ((Hirata et al., 2012)), auditory organs (Weber et al., 2003) |
| **Platyzoa** | protonephridia (Thi-Kim Vu et al., 2015) | protonephridia (Thi-Kim Vu et al., 2015) | protonephridia (Thi-Kim Vu et al., 2015) | protonephridia (Thi-Kim Vu et al., 2015) |
| **Mollusca** | mantle (Li et al., 2016) | ? | ? | mantle (Li et al., 2016) |
| **Annelida** | ? | ? | root epidermis (Miyamoto et al., 2017) | ? |
| **Nematoda** | hypodermis, intestine, excretory cell, neurons (Nehrke and Melvin, 2002) | muscle, neurons, intestine, excretory cell (Tanis et al., 2009) (Spencer et al., 2011) | intestinal tract (Fei et al., 2003) | muscle, neurons, midgut, intestine, body wall, pharynx, excretory cell (Sherman et al., 2005) |
| **Acoela** | mouth | parenchyme, brain | brain, parenchyme | no expression revealed |
| **Nemertodermatida** | nerve cords | female gonads, testis | neurons | female gonads, gut wrapping cells |
| **Cnidaria** | ? | ? | ? | oral tissues (Zoccola et al., 2015) |
| **Vertebrata** | gill (Hwang, 2009), intestine stomach, endothelial cells, distal tubule of kidney, cardiac myocytes, gall bladder, epididymis, ovary, thymus, brain, pancreas, salivary gland, skin, sperm, osteoclasts, skeletal muscle (sum. in (Donowitz et al., 2013)) | gill (Hwang, 2009), intestine (Li et al., 2014), ubiquitous, tubules of kidney, neurons, brain, bone (sum. in (Arroyo et al., 2013)) | kidney, small intestine, liver, placenta, brain (sum. in (Bergeron et al., 2013)) | hepatocytes, proximal tubule, intestine, testis, cardiac myocytes, brain, endothelial cells, hair cells, cochlear hair cells, pancreatic duct, thyrocytes (sum. in (Alper and Sharma, 2013), gill (Hwang, 2009), auditory organs (Weber et al., 2003) |
| **Eubacteria/Archaea** | ph regulation, metabolism, salt tolerance (Brett et al., 2005) | ? (Höglund et al., 2011) (Gagnon and Delpire, 2013) | carbon dicarboxylate substrate transport (Hall and Pajor, 2005) | $HCO_3^-$, $C^4$-dicarboxylic acid metabolism (Compton et al., 2014) |
| **Filasterea/ Choanoflagellata** | ? ((Höglund et al., 2011), this study)) | ? (Höglund et al., 2011) (Gagnon and Delpire, 2013) | absent ((Höglund et al., 2011), this study) | ? ((Höglund et al., 2011), this study) |
| **Holomycota** | vacuole trafficking, regulation of cytosolic ph (Brett et al., 2005) | ? (Höglund et al., 2011) | ? (Saier et al., 1999) (Höglund et al., 2011) | ? (Parker and Boron, 2013) |

### ***HCN***

| | |
|---|---|
| **Arthropoda** | gills (Fehsenfeld and Weihrauch, 2016), olfactory neurons (Gisselmann et al., 2005), chemo-sensitivity (Chen and Wang, 2012) |
| **Platyzoa** | ? |
| **Mollusca** | ? |
| **Annelida** | ? |
| **Nematoda** | hypodermis |
| **Acoela** | brain |
| **Nemertodermatida** | gut epithelium |
| **Cnidaria** | ? |
| **Vertebrata** | kidney (Bolivar et al., 2008; Carrisoza-Gaytan et al., 2011), liver (Santoro and Tibbs, 1999), ovary (Yeh et al., 2008), pangreatic cells (El-Kholy et al., 2007), heart (DiFrancesco, 1993), brain (Luthi and McCormick, 1998) |
| **Eubacteria/Archaea** | intracellular cAMP, pH sensor, osmoregulation (Kuo et al., 2007; Nimigean et al., 2004) |
| **Filasterea/ Choanoflagellata** | ? |
| **Holomycota** | ? |

**Supplementary Table 1.** Compilation of excretion-related gene expression/role data for metazoan and non-metazoan taxa for genes investigated in this study, when data available. Data are based on this study unless stated otherwise. Question marks represent missing data.

pH meassurements

| concentration [µmol/l] | Ipul | Nvec |
|---|---|---|
| 0 | 8.14 | 8.53 |
| 50 | 8.14 | 8.53 |
| 100 | 8.14 | 8.52 |
| 200 | 8.14 | 8.50 |
| 500 | 8.12 | 8.46 |
| 1000 | 8.10 | 8.36 |

Nvec Excretion meassurements

| sample | conc1 [µmol/l | conc2 [µmol/l] |
|---|---|---|
| **pH = 8.53** | 41,39786873 | 41,39786873 |
| **pH = 8.36** | 41,39786873 | 39,30019181 |
| **pH = 8.53** | 43,60751072 | 43,60751072 |
| **pH = 8.36** | 39,30019181 | 39,30019181 |
| **pH = 8.53** | 30,30247522 | 28,76701446 |
| **pH = 8.36** | 30,30247522 | 28,76701446 |

meassured: 2 hour incubation of 7 adult Nematostella in 8 ml medium

concentration: µmol NH4 / 2 ml*2h

**Supplementary Table 2.** *N. vectensis* excretion in different ph

### Accession numbers of reference sequences

AMT/RH
(N.vectensis A7SSQ4amt1/4a) (N.vectensis A7RH04amt1/4b) (N.vectensis A7SQ16amt2/3c) (N.vectensis A7S731amt2/3a) (N.vectensis A7S3L2amt2/3e) (N.vectensis A7RNC3amt2/3b) (N.vectensis A7SGD4amt2/3d) (N.vectensis XM_001622754.1Rh1) (N.vectensis 156375209Rh2) (N.vectensis 156369780Rh3) (B.floridae C3Z1J5) (B.floridae C3Y022) (B.floridae C3YB73) (B.floridae C3Z1G1) (B.floridae D7UQE4) (B.floridae D7UQE3) (B.floridae D7UQE2) (B.floridae D7UQE0) (B.floridae D7UQE1) (B.floridae D7UQD9) (B.floridae C3YB74) (B.floridae C3YB79) (C.gigas K1R302) (C.gigas K1QL97) (C.gigas K1QFY1) (C.gigas K1PJE2) (C.gigas K1RSL2) (C.intestinalis Q5VHU3) (C.intestinalis Q6XZ10) (C.intestinalis Q6XZ09) (C.intestinalis Q6XZ08) (C.intestinalis Q5VHU5) (C.intestinalis Q5VHU4) (T. adhaerens B3SC87) (T. adhaerens B3SC46) (T. adhaerens 196016156) (T. adhaerens 196016158) (T. adhaerens B3RRT6) (C.elegans spP54145) (C.elegans Q17663) (C.elegans Q9N2M5) (C.elegans Q20605) (C.elegans Q9N2M4) (C.elegans Q21565) (S.purpuratus W4Y0G6) (S.purpuratus W4YF70) (S.purpuratus W4Y3R7) (L.gigantea V4CRP5) (L.gigantea V3ZVE4) (L.gigantea V4AP67) (L.gigantea V3ZBD8) (L.gigantea 676492623) (L.gigantea 676434770) (C.teleta R7TS58) (C.teleta R7V3I9) (C.teleta R7U542) (C.teleta 443734385) (C.teleta 443691773) (C.teleta R7UG21) (H.sapiens Q02094) (H.sapiens Q9UBD6) (H.sapiens Q9H310) (M.leidyi ML10396a) (M.leidyi ML00673a) (M.leidyi ML018017a) (A.mellifera Q1L729) (A.mellifera Q38SD4) (D.sechellia B4HDZ9) (D.yakuba A0A0R1E419) (D.persimilis B4GLA6) (D.busckii A0A0M3QYI4) (D.melanogaster M9PEN8) (D.melanogaster Q9VFA9) (D.busckii A0A0M4E893) (D.pseudoobscura pseudoobscura Q4VUH9) (A.gambiae 118793733) (A.gambiae Q7Z1M1) (A. sinensis A0A084W5B9) (A.darling W5J3B1) (A.darling 568253901) (G.cydonium c.2546950) (A.queenslandica 761911958) (A.queenslandica 340386356) (G.pyriformis U3LYH9) (G.pyriformis U3LY22) (G.pyriformis U3LZ95) (H.cylindrosporum Q96UX9) (H.cylindrosporum Q96UY0) (H.cylindrosporum Q8NKD5) (S.pombe Q9C0V1) (S.pombe Q9US00) (S.cerevisiae P41948) (S.cerevisiae P40260) (S.cerevisiae P53390) (O.sativa Q8S230) (O.sativa Q8S233) (O.sativa Q84KJ7) (A.thaliana Q9M6N7) (O.sativa Q84KJ6) (O.sativa Q69T29) (O.sativa Q851M9) (O.sativa Q7XQ12) (O.sativa A0A0P0WCF8) (O.sativa Q6K9G1) (O.sativa Q6K9G3) (A.thaliana Q9ZPJ8) (A.thaliana Q9SQH9) (A.thaliana Q9LK16) (A.thaliana P54144) (A.thaliana Q9SVT8) (Archaea Q8TIE5 [Methanosarcina) (Archaea Q8TZ86 [Methanopyrus) (Archaea A0A075IBT0 [marine thaumarchaeote) (Archaea A0A075HAR1 [marine thaumarchaeote) (Archaea A0A075I7G1 [marine thaumarchaeote) (Archaea G4RK28 [Thermoproteus) (Archaea A0A075GR62 [marine thaumarchaeote) (Archaea A0A075GM15 [marine thaumarchaeote) (Archaea A0A075FUF3 [marine thaumarchaeote) (Archaea A0A075H4I2 [marine thaumarchaeote) (Archaea A0A075HUT8 [marine thaumarchaeote) (Archaea Q8TJ70 [Methanosarcina) (Archaea A0A075ICL8 [marine euryarchaeote) (Archaea A0A089ZF16 [Methanobacterium) (Bacteria 769129665 [Lachnospiraceae) (Bacteria 736086132 [Lachnospiraceae) (Bacteria E0RUI [Butyrivibrio) (Bacteria 769141800 [Lachnospiraceae) (Bacteria 769173412[Clostridium) (Bacteria 291528577 [Eubacterium]) (Bacteria 737671611 [Lachnospiraceae) (Bacteria 736431606 [Butyrivibrio) (Bacteria 490181861 [Clostridium) (Bacteria 932916429 [Anaerostipes) (Bacteria 736105025 [Lachnospiraceae) (Bacteria 769254574 [Lachnospiraceae) (Bacteria 551041730 [Lachnospiraceae) (Bacteria Q07429 [Bacillus) (Bacteria O66515 [Aquifex) (Bacteria A0A0D0ITJ4 [Pseudomonas) (Bacteria P69681 [Escherichia)

SLC
(S.mediterranea.slc26a-2|m.283_sm.slc26a-2| g.283 sm.slc26a-2:109-1878) (S.mediterranea.slc13a-4|m.264_sm.slc13a-4| g.264 sm.slc13a-4:3-1787) (S.mediterranea.slc1a-3|m.5_sm.slc1a-3| g.5 sm.slc1a-3:155-1555) (S.mediterranea.slc1a-1|m.1_sm.slc1a-1| g.1 sm.slc1a-1:238-567) (S.mediterranea.slc4a-1|m.16_Sm.slc4a-1| g.16 Sm.slc4a-1:107-3799) (S.mediterranea.slc4a-2|m.25_Sm.slc4a-2| g.25 Sm.slc4a-2:37-2943) (S.mediterranea.slc1a-4|m.10_sm.slc1a-4| g.10 sm.slc1a-4:1-1443) (S.mediterranea.slc1a-5|m.12_sm.slc1a-5| g.12 sm.slc1a-5:202-1548) (S.mediterranea.slc1a-2|m.3_sm.slc1a-2| g.3 sm.slc1a-2:103-1890) (S.mediterranea.slc4a-6|m.55_Sm.slc4a-6| g.55 Sm.slc4a-6:17-3538) (S.mediterranea.slc4a-7|m.60_Sm.slc4a-7| g.60 Sm.slc4a-7:69-3083) (S.mediterranea.slc4a-3|m.34_Sm.slc4a-3| g.34 Sm.slc4a-3:94-5439) (S.mediterranea.slc4a-4|m.46_Sm.slc4a-4| g.46 Sm.slc4a-4:296-2971) (S.mediterranea.slc4a-8|m.67_Sm.slc4a-8| g.67 Sm.slc4a-8:119-2866) (S.mediterranea.slc4a-5|m.52_Sm.slc4a-5| g.52 Sm.slc4a-5:50-2626) (S.mediterranea.slc4a-9|m.75_Sm.slc4a-9| g.75 Sm.slc4a-9:1-1545) (S.mediterranea.slc4a-10|m.78_Sm.slc4a-10| g.78 Sm.slc4a-10:3-509) (S.mediterranea.slc26a-7|m.313_sm.slc26a-7| g.313 sm.slc26a-7:240-2279) (S.mediterranea.slc26a-10|m.334_sm.slc26a-10| g.334 sm.slc26a-10:284-1114) (S.mediterranea.slc26a-6|m.308_sm.slc26a-6| g.308 sm.slc26a-6:145-2025) (S.mediterranea.slc26a-9|m.330_sm.slc26a-9| g.330 sm.slc26a-9:1-1161) (S.mediterranea.slc26a-5|m.303_sm.slc26a-5| g.303 sm.slc26a-5:128-1189) (S.mediterranea.slc26a-4|m.297_sm.slc26a-4| g.297 sm.slc26a-4:279-1808) (S.mediterranea.slc26a-8|m.320_sm.slc26a-8| g.320 sm.slc26a-8:55-2334) (S.mediterranea.slc26a-3|m.289_sm.slc26a-3| g.289 sm.slc26a-3:757-2277) (S.mediterranea.slc26a-1|m.280_sm.slc26a-1| g.280 sm.slc26a-1:28-2529) (S.mediterranea.slc26a-5|m.304_sm.slc26a-5| g.304 sm.slc26a-5:1237-2172) (S.mediterranea.slc26a-4|m.298_sm.slc26a-4| g.298 sm.slc26a-4:1690-2163) (S.mediterranea.slc26a-3|m.290_sm.slc26a-3| g.290 sm.slc26a-3:1-489) (S.mediterranea.slc13a-6|m.275_sm.slc13a-6| g.275 sm.slc13a-6:54-1754) (S.mediterranea.slc13a-5|m.267_sm.slc13a-5| g.267 sm.slc13a-5:88-1785) (S.mediterranea.slc13a-

7|m.278_sm.slc13a-7| g.278 sm.slc13a-7:41-1714) (S.mediterranea.slc13a-3|m.259_sm.slc13a-3| g.259 sm.slc13a-3:69-1811) (S.mediterranea.slc13a-2|m.255_sm.slc13a-2| g.255 sm.slc13a-2:15-287) (S.mediterranea.slc12a-2|m.218_sm.slc12a-2| g.218 sm.slc12a-2:30-3044) (S.mediterranea.slc12a-3|m.229_sm.slc12a-3| g.229 sm.slc12a-3:759-1625) (S.mediterranea.slc12a-4|m.234_sm.slc12a-4| g.234 sm.slc12a-4:327-2621) (S.mediterranea.slc12a-1|m.210_sm.slc12a-1| g.210 sm.slc12a-1:86-3280) (S.mediterranea.slc12a-5|m.244_sm.slc12a-5| g.244 sm.slc12a-5:187-2193) (S.mediterranea.slc12a-4|m.235_sm.slc12a-4| g.235 sm.slc12a-4:2647-3723) (S.mediterranea.slc9a-1|m.159_sm.slc9a-1| g.159 sm.slc9a-1:132-2207) (S.mediterranea.slc9a-4|m.174_sm.slc9a-4| g.174 sm.slc9a-4:112-2274) (S.mediterranea.slc9a-7|m.191_sm.slc9a-7| g.191 sm.slc9a-7:31-1950) (S.mediterranea.slc9a-8|m.196_sm.slc9a-8| g.196 sm.slc9a-8:3-1991) (S.mediterranea.slc9a-2|m.162_sm.slc9a-2| g.162 sm.slc9a-2:45-2078) (S.mediterranea.slc9a-5|m.180_sm.slc9a-5| g.180 sm.slc9a-5:94-2115) (S.mediterranea.slc9a-3|m.166_sm.slc9a-3| g.166 sm.slc9a-3:34-2712) (S.mediterranea.slc9a-9|m.199_sm.slc9a-9| g.199 sm.slc9a-9:889-2871) (S.mediterranea.slc8a-1|m.113_Sm.slc8a-1| g.113 Sm.slc8a-1:350-3103) (S.mediterranea.slc8a-2|m.124_Sm.slc8a-2| g.124 Sm.slc8a-2:82-2637) (S.mediterranea.slc8a-4|m.139_Sm.slc8a-4| g.139 Sm.slc8a-4:354-2546) (S.mediterranea.slc8a-5|m.149_Sm.slc8a-5| g.149 Sm.slc8a-5:68-2665) (S.mediterranea.slc8a-3|m.132_Sm.slc8a-3| g.132 Sm.slc8a-3:1114-2832) (S.mediterranea.slc8a-3|m.133_Sm.slc8a-3| g.133 Sm.slc8a-3:123-1193) (S.mediterranea.slc8a-4|m.140_Sm.slc8a-4| g.140 Sm.slc8a-4:1-450) (S.mediterranea.slc5a-2|m.86_Sm.slc5a-2| g.86 Sm.slc5a-2:62-1897) (S.mediterranea.slc5a-1|m.80_Sm.slc5a-1| g.80 Sm.slc5a-1:66-1862) (S.mediterranea.slc5a-3|m.97_Sm.slc5a-3| g.97 Sm.slc5a-3:53-1801) (S.mediterranea.slc5a-4|m.102_Sm.slc5a-4| g.102 Sm.slc5a-4:22-1860) (N.vectensis 173595) (N.vectensis 230013) (N.vectensis 1 228981) (N.vectensis 16302) (N.vectensis XP_001636607.1) (N.vectensis 156223712) (N.vectensis 89958) (N.vectensis 31622) (N.vectensis XP_001634186.1) (N.vectensis 156221266) (N.vectensis 238013) (N.vectensis 104399) (N.vectensis A7SQZ6) (N.vectensis A7S331) (N.vectensis XM_001623843.1) (N.vectensis XM_001637857.1) (N.vectensis XP_001641754.1) (N.vectensis XM_001641374.1) (N.vectensis XM_001626153.1) (N.vectensis XP_001626203.1) (N.vectensis XM_001632684.1) (N.vectensis XP_001632734.1) (N.vectensis XM_001635065.1) (N.vectensis XP_001635115.1) (N.vectensis A7SNT1) (N.vectensis A7SHZ4) (N.vectensis A7S542) (N.vectensis A7SC64) (N.vectensis A7S539) (N.vectensis A7SX54) (N.vectensis A7SWE4) (N.vectensis A7S2I0) (N.vectensis XP_001638856.1) (N.vectensis 156225980) (N.vectensis XP_001627572.1) (N.vectensis 156214486) (N.vectensis XP_001637705.1) (N.vectensis 156224819) (A.mediterranea A0A1J0SZ44) (A.mediterranea A0A1J0SLW4) (A.mediterranea A0A1J0TAF6) (A.queenslandica I1FF21) (A.queenslandica I1FF19) (A.queenslandica I1FF22) (A.queenslandica I1FS59) (H.sapiens P43004) (H.sapiens P43003) (H.sapiens P48664) (H.sapiens O00341) (H.sapiens P43005) (H.sapiens P43007) (H.sapiens Q15758) (H.sapiens Q2Y0W8) (H.sapiens Q6U841) (H.sapiens Q9Y6M7) (H.sapiens Q9Y6M7) (H.sapiens Q9Y6M7) (H.sapiens Q9Y6M7) (H.sapiens Q9Y6R1) (H.sapiens Q9BY07) (H.sapiens Q96Q91) (H.sapiens P04920) (H.sapiens P48751) (H.sapiens P02730) (H.sapiens Q8NBS3) (H.sapiens P50443) (H.sapiens Q9H2B4) (H.sapiens O43511) (H.sapiens P40879) (H.sapiens P58743) (H.sapiens Q9BXS9) (H.sapiens Q7LBE3) (H.sapiens Q96RN1) (H.sapiens Q8TE54) (H.sapiens Q86WA9) (H.sapiens Q8NG04) (H.sapiens Q13183) (H.sapiens Q86YT5) (H.sapiens Q8WWT9) (H.sapiens Q9BZW2) (H.sapiens Q9UKG4) (H.sapiens P55011) (H.sapiens Q13621) (H.sapiens P55017) (H.sapiens Q9BXP2) (H.sapiens Q9UHW9) (H.sapiens Q9UP95) (H.sapiens Q9H2X9) (H.sapiens Q9Y666) (H.sapiens A0AV02) (H.sapiens Q9UBY0) (H.sapiens Q6AI14) (H.sapiens P19634) (H.sapiens P48764) (H.sapiens Q14940) (H.sapiens Q92581) (H.sapiens Q96T83) (H.sapiens Q8IVB4) (H.sapiens Q9Y2E8) (H.sapiens P32418) (H.sapiens P57103) (H.sapiens Q9UPR5) (H.sapiens P13866) (H.sapiens Q9NY91) (H.sapiens P31639) (H.sapiens Q2M3M2) (H.sapiens A0PJK1) (H.sapiens Q8WWX8) (H.sapiens P53794) (H.sapiens Q8N695) (H.sapiens Q1EHB4) (H.sapiens Q92911) (H.sapiens Q9Y289) (H.sapiens Q9GZV3)

AQUAPORINS
(A.thaliana P43286|PIP21) (A.thaliana P43287|PIP22) (A.thaliana P93004|PIP27) (A.thaliana P61837|PIP11) (A.thaliana Q08733|PIP13) (A.thaliana P30302|PIP23) (A.thaliana Q39196|PIP14) (A.thaliana Q06611|PIP12) (A.thaliana Q9FF53|PIP24) (A.thaliana Q9ZV07|PIP26) (A.thaliana Q9ZVX8|PIP28) (A.thaliana Q9SV31|PIP25) (A.thaliana Q8LAA6|PIP15) (X.tropicalis F6SCI3) (X.tropicalis Q5EBG0) (X.tropicalis F6RVN7) (X.tropicalis F6QPA9) (X.tropicalis (X.tropicalis F7CQ45) (X.tropicalis F7CFD3) (X.tropicalis F6QEC2) (X.tropicalis F6QNV6) (X.tropicalis F6Z564) (X.tropicalis A0A060PX33) (X.tropicalis Q28DL6) (X.tropicalis Q5FW23) (X.tropicalis A0A1B8YA67) (X.tropicalis A4IGW1) (X.tropicalis Q6DJ01) (X.tropicalis F6RVG6) (X.tropicalis F6YXD3) (X.tropicalis B0BM29) (X.tropicalis F6V0M7 (X.tropicalis A0A1B8XXZ7) (X.tropicalis F6S813) (X.tropicalis A0A1B8XXT0) (X.tropicalis F6QCC5) (X.tropicalis B4F717) (X.tropicalis F6Z1F2) (X.tropicalis F6Q4J3) (H.sapiens P29972) (H.sapiens P41181) (H.sapiens Q92482) (H.sapiens P55087) (H.sapiens P55064) (H.sapiens Q13520) (H.sapiens O14520) (H.sapiens O94778) (H.sapiens O43315) (H.sapiens Q96PS8) (H.sapiens Q8NBQ7) (H.sapiens Q8IXF9) (H.sapiens A6NM10) (C.gigas K1QMA6) (C.gigas K1QP58) (C.gigas K1QV92) (C.gigas K1QSP9) (C.gigas K1PA25) (C.gigas K1QNC6) (C.gigas K1QC31) (C.gigas K1RM00) (C.gigas K1R4H0) (C.gigas K1QUZ8) (C.gigas K1RGW0) (C.gigas K1RAK5) (C.gigas K1QBQ1) (C.gigas K1QCC4) (C.gigas K1RA04) (D.melanogaster P23645) (D.melanogaster P23645) (D.melanogaster E1JH55) (D.melanogaster Q7KY01) (D.melanogaster M9PB86) (D.melanogaster A0A0B4KFZ1) (D.melanogaster A0A0C4DHF9) (D.melanogaster

Q9W1M2) (D.melanogaster Q9W1M4) (D.melanogaster Q9W1M3) (D.melanogaster D1Z394) (D.melanogaster Q8MLR2) (D.melanogaster D3DMU9) (D.melanogaster F3YDC9) (D.melanogaster Q6NR72) (D.melanogaster A0A0B4KFN8) (D.melanogaster A0A0B4KET4) (D.melanogaster A0A0B4KFD6) (D.melanogaster A0A0B4LG55) (D.melanogaster A1Z8L8) (D.melanogaster Q95TS2) (D.melanogaster J7K3P9) (D.melanogaster H6V591 (D.melanogaster Q8T0N8 (D.melanogaster H5V8H0 (D.melanogaster H8F4Q9) (C.elegans Q19949) (C.elegans Q8IG23) (C.elegans Q18352) (C.elegans Q21473) (C.elegans Q9XW36) (C.elegans D5MCS1) (C.elegans H2FLH4) (C.elegans Q09369) (C.elegans G5EEK0) (C.elegans O46024) (C.elegans Q17571) (C.elegans Q7JMQ6) (C.elegans Q7JL16) (C.elegans Q7Z137) (C.elegans Q7Z138) (C.elegans Q18469) (C.elegans Q7YWQ1) (C.elegans A0A131MBS2) (C.elegans A0A061ADS9) (C.elegans A0A2K5AU16) (C.elegans A0A2K5AU20) (D.discoideum Q54WT8) (D.discoideum Q9U8P7) (D.discoideum Q54V53) (D.discoideum Q8SSP2) (D.discoideum Q54FQ9) (T.adhaerens B3RKX8) (T.adhaerens B3RR23) (A.queenslandica A0A1X7VPT7) (A.queenslandica A0A1X7UWA0) (A.queenslandica A0A1X7VQ25) (A.queenslandica A0A1X7UXK7) (A.queenslandica A0A1X7VNT9) (S.purpuratus W4ZC70) (S.purpuratus W4Y8N5) (S.purpuratus W4XM79) (S.purpuratus W4ZGE5) (S.purpuratus W4YZV4) (S.purpuratus W4XVQ3) (S.purpuratus W4YZV5) (S.purpuratus W4Y910) (S.purpuratus W4Z8B2) (S.purpuratus W4XXY7) (S.purpuratus W4YYT0) (S.purpuratus W4Z759) (S.purpuratus W4XQ97) (S.purpuratus W4XRA4) (S.purpuratus W4YD56) (N.vectensis A7S565) (N.vectensis A7S514) (N.vectensis A7S515) (N.vectensis A7T0G8) (N.vectensis A7RU64) (N.vectensis A7RU63) (N.vectensis A7S5L0) (N.vectensis A7T581) (N.vectensis A7T6P4) (N.vectensis A7SKN0)

SLIPINS
(C.elegans Q27433GN) (C.elegans Q19958) (C.elegans Q21190) (C.elegans Q22165) (C.elegans Q19200) (C.elegans Q20657) (C.elegans G5ED76) (C.elegans H2FLJ1) (C.elegans Q9XWC6) (C.elegans H2L024) (D.melanogaster Q9VZA4) (D.melanogaster Q8MZ13) (D.melanogaster Q9VWL0) (D.melanogaster Q9W1F7) (H.sapiens Q9NP85) (H.sapiens P27105) (H.sapiens Q9UJZ1) (H.sapiens A0A024R882) (H.sapiens Q8TAV4) (H.sapiens Q9UBI4) (X.tropicalis Q6GLC6) (X.tropicalis Q6P362) (X.tropicalis F6VV69) (X.tropicalis F6WE98) (N.vectensis jgi|Nemve1|91851) (N.vectensis jgi|Nemve1|247670) (N.vectensis jgi|Nemve1|41366) (N.vectensis jgi|Nemve1|218685) (N.vectensis jgi|Nemve1|147236) (N.vectensis jgi|Nemve1|98373) (N.vectensis jgi|Nemve1|164373) (N.vectensis jgi|Nemve1|98301) (Bacteria A0E8T9 [Paramecium) (Bacteria Q5A411 [Candida) (Bacteria I7M6K2 [Tetrahymena) (A.queenslandica I1FPX0) (A.queenslandica I1FPW9) (A.queenslandica I1EJ07) (A.queenslandica I1FUT2) (A.queenslandica I1FXM9) (T.adhaerens B3SEH1) (T.adhaerens B3RKW0) (T.adhaerens B3RW09) (T.adhaerens B3RSS8) (T.adhaerens B3RKV9)

CD2AP
(H.sapiens Q9Y5K6) (D.melanogaster Q9Y154) (A.queenslandica XP_003386167.1) (N.vectensis XP_001630773.1) (H.sapiens P02549) (H.sapiens P11277) (D.melanogaster Q9Y154) (S.kowalevskii XP_006826005.1) (H.vulgaris vulgaris) (T.adhaerens RDD45964.1)

NEPHRIN/KIRRE
(H.sapiens Q96J84) (H.sapiens Q8IZU9) (H.sapiens Q6UWL6) (H.sapiens O60500) (A.mellifera XP_026296514.1) (C.gigas EKC26159.1) (T.castaneum XP_008190700.1) (X.tropicalis F7EAD2) (X.tropicalis F7BA02) (D.melanogaster Q08180) (D.melanogaster Q9N9Y9) (D.melanogaster Q9V787) (D.melanogaster Q9V4Y0) (A.cerana PBC28977.1) (M.lignano PAA87588.1) (M.lignano PAA94659.1) (S.kowalevskii NP_001164704.1) (C.elegans B1Q236) (C.elegans Q9U3P2) (B.glabrata A0A075T6F5)

ZO1
(H.sapiens Q07157) (H.sapiens Q9UDY2) (H.sapiens Q2VPE5) (D.melanogaster Q94880) (T.adhaerens B3S7T9) (C.elegans Q8I103) (S.purpuratus XP_782687.2) (N.vectensis A7S398) (D.melanogaster P31007) (M.musculus Q62108) (GQ290472.1_Capsaspora) (GQ290473_Capsaspora)

CA
(N.vectensis A7T609) (N.vectensis A6QR78) (N.vectensis A7S2D8 CA a1) (N.vectensis A7SHT0) (N.vectensis A7S717) (N.vectensis A7SKA8 CA a3) (N.vectensis A7RRH8) (N.vectensis A7SHS9 CA a2) (N.vectensis A7RR00) (N.vectensis A7S762) (H.sapiens P00918) (H.sapiens P00915) (H.sapiens Q8N1Q1) (H.sapiens P07451) (H.sapiens P43166) (H.sapiens P35218) (H.sapiens Q9Y2D0) (H.sapiens O43570) (H.sapiens Q9ULX7) (H.sapiens Q16790) (H.sapiens P35219) (H.sapiens P22748) (H.sapiens P23280) (H.sapiens O75493) (H.sapiens Q9NS85) (T.castaneum D6WET4) (T.castaneum A0A139WMN7) (T.castaneum D7EIC1) (T.castaneum D2A0V8) (T.castaneum D2A110) (T.castaneum A0A139WMC5) (T.castaneum D6W9D1) (T.castaneum D6W9D0) (T.castaneum D6W9D3) (A.queenslandica A6QR77) (A.queenslandica I1FTL5) (A.queenslandica A6QR75) (A.queenslandica I1FAY3) (A.queenslandica I1FAY2) (A.queenslandica I1FAY4) (A.queenslandica A6QR76) (A.queenslandica I1GDV3) (A.queenslandica I1G206) (T.adhaerens B3RXW0) (T.adhaerens B3RJD2) (T.adhaerens B3RVV0) (T.adhaerens

B3RKE2) (T.adhaerens B3RKE1) (T.adhaerens B3RWG7) (T.adhaerens B3RKE3) (T.adhaerens B3RKE1) (T.adhaerens B3RKE0) (C.teleta R7T953) (C.teleta R7U445) (C.teleta R7UU94) (C.teleta R7V8T6) (C.teleta R7TR34) (C.teleta R7UAK2) (C.teleta R7TGJ8) (C.teleta R7V5W2) (C.teleta R7TB34) (C.teleta R7U704) (C.teleta R7U0D8) (C.teleta R7UH28) (S.purpuratus W4Y433) (S.purpuratus W4YCH9) (S.purpuratus W4YCI0) (S.purpuratus W4YNP4) (S.purpuratus W4Z2K3) (S.purpuratus W4XCN3) (S.purpuratus W4YB71) (S.purpuratus W4XBF4) (S.purpuratus W4Z9I4) (S.purpuratus Q0QBU7) (S.purpuratus W4Y9V4) (S.purpuratus W4XYY9) (S.purpuratus W4XL56) (S.purpuratus W4ZC38)

V-ATPASE B
(N.vectensis A7SRN1) (N.vectensis A7S0L1V-atpaseB) (T.adhaerens B3SAY3) (T.adhaerens B3RW75) (C.teleta R7TIR2) (C.teleta R7V2Q5) (A.queenslandica I1GEG2) (A.queenslandica I1FHW5) (H.sapiens P21281) (H.sapiens P15313) (H.sapiens P38606) (S.cerevisiae P16140) (S.cerevisiae P17255) (D.melanogaster P31409) (D.melanogaster Q27331) (D.melanogaster P48602) (C.gigas EKC36437.1) (S.kowalevskii NP_001171741.1) (A.californica XP_005091800.1) (D.discoideum Q76NU1) (D.discoideum P54647)

NKA
(D.purpureum F1A2S2) (D.purpureum Q95024) (A.queenslandica XP_011404074.1) (C.elegans G5EFV6) (C.elegans P90735) (T.adhaerens B3RIH3) (N.vectensis A7S6M1 NKA A2) (N.vectensis A7SII2 NKA A1) (H.sapiens P50993) (H.sapiens P13637) (H.sapiens P05023) (H.sapiens Q13733) (C.teleta R7V8F1) (C.teleta R7VBZ9) (S.purpuratus W4ZCC9) (D.melanogaster P13607)

HCN
(H.sapiens O60741) (H.sapiens Q9UL51) (H.sapiens Q9P1Z3) (H.sapiens Q9Y3Q4) (B.dorsalis A0A034V7Z9) (B.dorsalis A0A034V8L7) (S.purpuratus NP_999729.1) (S.purpuratus NP_001028182.1) (C.intestinalis H6V963) (C.intestinalis H6V964) (D.melanogaster AAD42059.1) (N.vectensis A0A0S2KP17) (N.vectensis A0A0S2KP23) (A.queenslandica XP_003385808.1) (A.queenslandica XP_011410082.2) (T.adhaerens XP_002111266.1) (T.adhaerens XP_002116397.1)

### Accession numbers of transcripts from Xenacoelomorph transcriptomes

AMT/RH
(H.miamia 98027843) (H.miamia 98057167) (H.miamia 98018981) (H.miamia 98054821) (C.macropyga 395.1) (C.macropyga 24654.1) (C.macropyga 14995.1) (C.macropyga 16173.1) (C.macropyga 10395.1) (C.macropyga 4405.1) (C.macropyga 1268.1) (C.macropyga 474.1) (I.pulchra 17191.1amt2/3b) (I.pulchra 22424.1amt-like) (I.pulchra 15010.1amt2/3a) (I.pulchra 35272.1amt2/3c) (I.pulchra 3446.1Rh) (I.pulchra 8667.1amt1/4b) (I.pulchra 9476.1amt1/4a) (X.bocki 7955.1) (X.profunda 10079.1) (M.stichopi 7327.1Rh) (M.stichopi 22314.1amt) (N.westbladi 55127.0) (N.westbladi 37503.0) (N.westbladi 48661.0) (N.westbladi 49524.0) (D.longitubus c31154) (D.longitubus c29820) (D.longitubus c34329) (D.gymnopharyngeus c4043) (C.submaculatum c14126) (C.submaculatum c14702) (C.submaculatum c18521)

SLC
(I.pulchra 12372.1SLC12a) (I.pulchra 8886.1SLC12b) (I.pulchra 10040.1SLC9) (I.pulchra 8649.1SLC8) (I.pulchra 13252.1SLC13) (I.pulchra 6460.1SLC4c) (I.pulchra 15145.1SLC4a) (I.pulchra 17707.1SLC4b) (I.pulchra 11412.1SLC5a) (I.pulchra 25325.1SLC5b) (I.pulchra 5795.1SLC26a) (I.pulchra 5099.1SLC26b) (I.pulchra 17762.1SLC1b) (I.pulchra 5285.1SLC1a) (I.pulchra 1359.1SLC1c) (M.stichopi 1129.1SLC26b) (M.stichopi 18197.1SLC26a) (M.stichopi 11423.1SLC12c) (M.stichopi 25290.1 SLC12b) (M.stichopi 12379.1SLC12a) (M.stichopi 6906.1SLC13a) (M.stichopi 24659.1SLC13b) (M.stichopi 19295.1SLC13c) (M.stichopi 18299.1SLC13d) (M.stichopi 5799.1SLC4b) (M.stichopi 7282.1SLC4c) (M.stichopi 18993.1SLC4a) (M.stichopi 7085.1 SLC8) (M.stichopi 11587.1 SLC9) (M.stichopi 8391.1SLC5a) (M.stichopi 22651.1SLC5b) (M.stichopi 20595.1SLC1)

AQUAPORINS
(X.bocki 30275.1) (X.bocki 11732.1) (X.bocki 16293.1) (X.bocki 2579.1) (X.bocki 3609.1) (X.bocki 6004.1) (X.profunda 16190.1) (X.profunda 2887.1) (X.profunda 5191.1) (X.profunda 16480.1) (C.macropyga 11373.1) (C.macropyga 18775.1) (C.macropyga 14142.1) (C.macropyga 16171.1) (C.macropyga 11010.1) (C.macropyga 20222.1) (C.macropyga 16428.1) (C.macropyga 1448.1) (C.macropyga 17222.1) (C.macropyga 15603.1) (E.macrobursalium c7828) (C.submaculatum c13797) (C.submaculatum c17016) (C.submaculatum c15544) (D.gymnopharyngeus c13920) (D.gymnopharyngeus c11428) (D.longitubus c31442) (D.longitubus c33092) (D.longitubus c29236) (H.miamia 98039895) (H.miamia 98046060) (H.miamia 98013247) (M.stichopi. 20229AQf) (M.stichopi 7238AQd) (M.stichopi 9653AQc) (M.stichopi 11719AQe) (M.stichopi 16317AQa) (M.stichopi

24323AQb) (I.pulchra 42598AQb) (I.pulchra 28723AQf) (I.pulchra 4312AQa) (I.pulchra 1616AQc) (I.pulchra 21547AQe) (I.pulchra 44478AQd) (I.pulchra 42138AQg)

SLIPINS
(X.bocki 20883.1) (X.bocki 12758.1) (X.bocki 1047.1) (Ascoparia sp. 9383.0) (Ascoparia sp. 17607.0) (Ascoparia sp. 20287.1) (Ascoparia sp. 28149.0) (Sterreria sp. c17109) (C.macropyga 229.1) (C.macropyga 14931.1) (C.macropyga 729.1) (C.macropyga 10774.1) (C.submaculatum c17900) (C.submaculatum c18531) (C.submaculatum c19891) (N.westbladi 47675.) (N.westbladi 52579.0) (N.westbladi 25494.0) (I.pulchra 4222.1Stomatin/podocinA) (I.pulchra 8048.1 Stomatin/podocinB) (I.pulchra 1122.1Stomatin/podocinC) (M.stichopi 2017.2Stomatin/podocinE) (M.stichopi 4077.1Stomatin/podocinD) (M.stichopi 4083.1Stomatin/podocinC) (M.stichopi 4165.1Stomatin/podocinB) (M.stichopi 7738.1Stomatin/podocinA) (D.gymnopharyngeus c2158) (D.gymnopharyngeus c13414) (D.longitubus c30579) (D.longitubus c33065) (D.longitubus c36799) (E.macrobursalium c5714) (E.macrobursalium c6060) (E.macrobursalium c8407) (H.miamia 98000817) (H.miamia 98001079) (H.miamia 98003185) (H.miamia 98003968) (H.miamia 98009885) (H.miamia 98030377) (H.miamia 98031305) (H.miamia 98032202)

CD2AP
(I.pulchra 2785.1) (X.bocki 1192.1) (M.stichopi 4919.1) (N.westbladi 47339.0) (N.westbladi 51312.0) (D.gymnopharyngeus c14992)

NEPHRIN/KIRRE
(H.miamia 9805464) (H.miamia 98022051) (C.macropyga 2490.1) (C.macropyga 2933.1) (C.macropyga 5188.1) (I.pulchra 16124.1Nephrin/Kirre2) (I.pulchra 13602.1Nephrin/Kirre1) (I.pulchra 14166.1Nephrin/Kirre3) (X.bocki 5576.1) (X.bocki 12500.1) (X.profunda c18867) (X.profunda 14967.1) (X.profunda 13792.1) (M.stichopi 19607.1Nephrin/Kirre2) (M.stichopi 2457.1Nephrin/Kirre3) (M.stichopi 17803.1Nephrin/Kirre1) (N.westbladi 48390.0) (N.westbladi 51630.0) (D.longitubus c36603) (D.longitubus c38278) (D.longitubus c37893) (D.gymnopharyngeus c10380) (D.gymnopharyngeus c10580) (C.submaculatum c17370) (C.submaculatum c15556) (C.submaculatum c17370) (C.submaculatum c17306) (E.macrobursalium c4372)

ZO1
(H.miamia 98057262) (C.macropyga 5972.1) (I.pulchra 8350.1) (X.bocki 3789.1) (M.stichopi 7010.1) (N.westbladi 51947.0) (D.longitubus c38032) (D.gymnopharyngeus c12822) (D.gymnopharyngeus c13251) (C.submaculatum c19373)

CA
(Ascoparia sp. 5478.0) (Ascoparia sp. 18043.0) (Ascoparia sp. 25649.0) (Ascoparia sp. 2777.0) (C.submaculatum c1687) (C.submaculatum c13257) (C.submaculatum c5059) (C.submaculatum c13664) (D.gymnopharyngeus c13170) (D.gymnopharyngeus c10609) (D.gymnopharyngeus c11439) (D.gymnopharyngeus c13170) (C.macropyga 16271.1) (C.macropyga 3418.1) (C.macropyga 19581.1) (C.macropyga 21507.1) (C.macropyga 18799.1) (C.macropyga 18136.1) (C.macropyga 13579.1) (C.macropyga 7849.1) (C.macropyga 1531.1) (C.macropyga 9483.1) (D.longitubus c35395) (D.longitubus c14121) (D.longitubus c28511) (D.longitubus c28413) (D.longitubus c33567) (E.macrobursalium c491) (E.macrobursalium c1233) (E.macrobursalium c7687) (E.macrobursalium c4985) (E.macrobursalium c122) (E.macrobursalium c6561) (E.macrobursalium c3860) (N.westbladi 29328.0) (N.westbladi 45461.0) (N.westbladi 39569.0) (N.westbladi 25845.0) (N.westbladi 45772.0) (N.westbladi 12921.0) (N.westbladi 50761.0) (H.miamia 98030173) (H.miamia 98042177) (H.miamia 98029235) (H.miamia 98046701) (H.miamia 98057584) (H.miamia 98008272) (H.miamia 98044099) (Sterreria sp. c22223) (X.profunda 20645.1) (X.profunda 8011.1) (X.profunda c9812) (X.profunda 22298.1) (X.profunda 10281.1) (X.profunda 19114.2) (X.bocki 3894.1) (X.bocki 4398.1) (X.bocki 1891.1) (X.bocki 13708.1) (X.bocki 25911.1) (X.bocki 46303.1) (X.bocki 29622.1) (X.bocki 29916.1) (I.pulchra 8550.1CA a1) (I.pulchra 9669.1CA a2) (I.pulchra 804.1CA a3) (I.pulchra 16851.1 CA a4) (I.pulchra 7996.1) (I.pulchra 12647.1) (I.pulchra 23555.1) (I.pulchra 11814.1) (I.pulchra 2042.1) (I.pulchra 22278.1) (I.pulchra 8550.1) (I.pulchra 804.1) (M.stichopi 5190.1CA a1) (M.stichopi 11897.1CA a2) (M.stichopi 6199.1CA a4) (M.stichopi 3473.1CA a3) (M.stichopi 16170.1) (M.stichopi 9713.1) (M.stichopi 27008.1)

V-ATPASE
(X.bocki 1513.1) (X.profunda 15609) (N.westbladi 51758) (I.pulchra 551.1V-atpaseB) (E.macrobursalium c7821) (C.macropyga 487.1) (D.longitubus c35334) (D.longitubus c35800) (D.longitubus c28900) (C.submaculatum c21601) (M.stichopi 24471.1V-atpaseB2) (M.stichopi 4970.1V-atpaseB1)

NKA
(H.miamia 98034285) (D.longitubus c37851) (C.submaculatum c18356) (C.macropyga 93.1) (X.bocki 812.1) (X.profunda 6567.1) (N.westbladi 52442.0) (M.stichopi 4323.1) (I.pulchra 596.1Na/K ATPaseA1) (I.pulchra

878.1Na/K ATPaseA2)

HCN
(I.pulchra 19714.1) (M.stichopi 18543.1)

**Supplementary Table 3.** Accession numbers of reference sequences used in Supplementary Figure 1.

*I.pulchra*

| GENE | average fold change | replica 1 | replica 2 | replica 3 | min | max | negative error | positive error |
|---|---|---|---|---|---|---|---|---|
| **rhesus** | 2.217183893 | 1.7168389 | 2.56415513 | 2.37055765 | 1.7168389 | 2.56415513 | 0.500344997 | 0.346971237 |
| **NKA a** | 2.596248698 | 2.17808859 | 2.6920789 | 2.9185786 | 2.17808859 | 2.9185786 | 0.418160107 | 0.322329902 |
| **NKA b** | 1.693353664 | 1.59134502 | 1.59429737 | 1.8944186 | 1.59134502 | 1.8944186 | 0.102008643 | 0.201064933 |
| **v-ATPase B** | 0.494488592 | 0.50926761 | 0.44218674 | 0.53201143 | 0.44218674 | 0.53201143 | 0.052301849 | 0.037522836 |
| **HCN** | 1.254995092 | 1.1888023 | 1.02014386 | 1.55603912 | 1.02014386 | 1.55603912 | 0.234851234 | 0.301044031 |
| **amt1/4 a** | 2.155580868 | 2.27056 | 2.15934753 | 2.03683508 | 2.03683508 | 2.27056 | 0.118745787 | 0.114979129 |
| **amt-like** | 1.600769598 | 1.64926114 | 1.521413 | 1.63163465 | 1.521413 | 1.64926114 | 0.079356595 | 0.048491541 |
| **amt1/4 b** | 2.691183113 | 2.94608692 | 2.57428365 | 2.55317876 | 2.55317876 | 2.94608692 | 0.138004349 | 0.25490381 |
| **amt2/3 a** | 1.860833749 | 1.76944979 | 1.32323389 | 2.48981757 | 1.32323389 | 2.48981757 | 0.537599859 | 0.628983821 |
| **amt2/3 b** | 2.072839483 | 2.45194857 | 1.83274149 | 1.93382839 | 1.83274149 | 2.45194857 | 0.240097992 | 0.379109085 |
| **amt2/3 c** | 1.314142054 | 1.24229458 | 1.15235077 | 1.54778081 | 1.15235077 | 1.54778081 | 0.161791289 | 0.233638758 |
| **aquaporin a** | 2.577641717 | 2.85374828 | 2.75857472 | 2.12060215 | 2.12060215 | 2.85374828 | 0.457039564 | 0.276106565 |
| **aquaporin b** | 2.793732272 | 2.78000575 | 3.45075469 | 2.15043638 | 2.15043638 | 3.45075469 | 0.643295893 | 0.657022417 |
| **aquaporin c** | 2.70322904 | 2.88856343 | 2.80330324 | 2.41782045 | 2.41782045 | 2.88856343 | 0.285408591 | 0.185334389 |
| **aquaporin d** | 1.776819335 | 1.54068 | 1.98835987 | 1.80141813 | 1.54068 | 1.98835987 | 0.236139336 | 0.211540536 |
| **aquaporin e** | 2.077006893 | 2.01976667 | 2.50102388 | 1.71023012 | 1.71023012 | 2.50102388 | 0.366776771 | 0.424016989 |
| **aquaporin f** | 1.318446722 | 1.06018873 | 1.45500979 | 1.44014164 | 1.06018873 | 1.45500979 | 0.25825799 | 0.136563071 |
| **aquaporin g** | 1.709907854 | 1.15172762 | 1.85739378 | 2.12060215 | 1.15172762 | 2.12060215 | 0.558180229 | 0.4106943 |
| **ca a** | 1.81344051 | 1.85094045 | 1.8379878 | 1.75139329 | 1.75139329 | 1.85094045 | 0.062047223 | 0.037499938 |
| **ca b** | 0.44152293 | 0.53183025 | 0.38334267 | 0.40939586 | 0.38334267 | 0.53183025 | 0.058180255 | 0.090307325 |
| **ca c** | 2.101369012 | 2.34990803 | 1.99340395 | 1.96079506 | 1.96079506 | 2.34990803 | 0.140573957 | 0.248539015 |
| **ca d** | 1.903583872 | 2.30433541 | 1.49964688 | 1.90676933 | 1.49964688 | 2.30433541 | 0.403936991 | 0.400751534 |
| **ca e** | 1.202587419 | 1.13455072 | 1.23432496 | 1.23888658 | 1.13455072 | 1.23888658 | 0.068036702 | 0.036299156 |
| **ca f** | 1.658881375 | 1.58664991 | 1.94214714 | 1.44784707 | 1.44784707 | 1.94214714 | 0.211034303 | 0.28326577 |
| **ca g** | 1.874941708 | 2.42800103 | 1.3377446 | 1.85907949 | 1.3377446 | 2.42800103 | 0.537197105 | 0.553059325 |
| **ca h** | 2.264203899 | 2.16749309 | 2.32859122 | 2.29652739 | 2.16749309 | 2.32859122 | 0.096710812 | 0.064387323 |
| **ca x** | 1.883258685 | 1.58149009 | 1.68487764 | 1.91528729 | 1.58149009 | 1.91528729 | 0.301768597 | 0.032028609 |

*N.vectensis*

| GENE | replica 1 | replica 2 | replica 3 | replica 4 | replica 5 | average fold change | min | max | negative error | positive error |
|---|---|---|---|---|---|---|---|---|---|---|
| **rhesus 1** | 0.66256924 | 0.58543965 | 0.5453938 | 0.2487322 | 0.42097777 | 0.492622531 | 0.2487322 | 0.66256924 | 0.243890335 | 0.16994671 |
| **rhesus 2** | 0.72148271 | 0.37254871 | 0.57360891 | 0.18700431 | 0.3513349 | 0.441195908 | 0.18700431 | 0.72148271 | 0.254191602 | 0.280286803 |
| **rhesus 3** | 1.16015502 | 0.99525448 | 0.67506186 | 0.60440554 | 1.29971411 | 0.946918203 | 0.60440554 | 1.29971411 | 0.342512661 | 0.352795906 |
| **nka 1** | 1.40156068 | 1.12475528 | 1.2076442 | 0.90980731 | 0.94960978 | 1.118675449 | 0.90980731 | 1.40156068 | 0.208868139 | 0.282885229 |
| **nka 2** | 0.8768764 | 0.9891905 | 1.16772962 | 1.90742149 | 0.67601966 | 1.123447537 | 0.67601966 | 1.90742149 | 0.447427872 | 0.783973957 |
| **amt2/3 a** | 1.00285841 | 0.49106638 | 0.44097256 | 0.59328951 | 0.42250718 | 0.590138809 | 0.42250718 | 1.00285841 | 0.167631628 | 0.412719604 |
| **amt2/3 b** | 1.30287673 | 1.06698811 | 0.57725253 | 0.52395472 | 1.27166109 | 0.948546639 | 0.52395472 | 1.30287673 | 0.424591915 | 0.354330096 |
| **amt2/3 e** | 1.64275158 | 1.10108814 | 1.47424144 | 2.81035319 | 3.15621983 | 2.036930837 | 1.10108814 | 3.15621983 | 0.935842699 | 1.11928899 |
| **amt1/4 b** | 2.14755411 | 3.61492989 | 2.33638672 | 1.6762092 | | 2.443769981 | 1.6762092 | 3.61492989 | 0.767560779 | 1.171159911 |
| **amt2/3 c** | 1.15684685 | 1.6376369 | | 0.5248662 | | 1.106449982 | 0.5248662 | 1.6376369 | 0.581583786 | 0.531186919 |
| **amt1/4 a** | 1.41776293 | 1.01899733 | 1.21231911 | | 0.70347258 | 1.088137987 | 0.70347258 | 1.41776293 | 0.384665408 | 0.329624941 |
| **amt2/3 d** | 0.88755083 | 0.7055751 | 1.01447442 | | 0.51000849 | 0.77940221 | 0.51000849 | 1.01447442 | 0.269393718 | 0.235072214 |
| **V ATPASE B** | 1.30911533 | 1.51601338 | 2.00097628 | 2.94070826 | | 1.941703313 | 1.30911533 | 2.94070826 | 0.632587983 | 0.999004949 |
| **CA 2** | 1.83671485 | 0.88322688 | 0.61075626 | 1.07265708 | | 1.100838767 | 0.61075626 | 1.83671485 | 0.490082508 | 0.735876079 |
| **CA 3** | 1.90235733 | 0.8797604 | 2.32532928 | 0.97401599 | 1.19622438 | 1.455537476 | 0.8797604 | 2.32532928 | 0.575777081 | 0.869791801 |
| **CA 1** | 0.65628293 | 0.83646141 | 0.58710676 | 0.51367514 | 0.38731996 | 0.596169239 | 0.38731996 | 0.83646141 | 0.208849278 | 0.240292173 |

**Supplementary Table 4.** QPCR raw data.

*I.pulchra*

| GENE | QPCR PRIMER F | QPCR PRIMER R |
|---|---|---|
| AqA | GCTATTCTCATCCACCTGCTCG | GCTTGGCTCTTGAAATCTTCTGG |
| AqB | GATAACACGCCTCAATGCCTTTC | TCACATCGTAGTTGTCTCCTCCTTC |
| AqC | TGTTCCAAATCAGAGTCATCCAGG | TTCGCTCACAGTTACTTCGCCTCG |
| AqD | TGGCACAACCACTGGGTCTC | TTCTCGGCGTCTTCGGGAGTAG |
| AqE | AGGCATCATTGTCCGTGTCTGC | TTTGTGTTCCTGGTTGGGGTGGTG |
| AqF | GCAACATCAACCCAAGCGTC | GAACAACCAGCCAGAGAACGAAC |
| AqG | TGGGACTGGGACCACTCGTTAC | CGAATGTCACCAATGAGATGCTCC |
| Rhesus | GAAGTTGTTCAGTGGGGAGAAGG | CTTGATGAGTGACATTGCCGTG |
| V-ATPase B | GAGTTTAGGTCTCCTTTTCCTCAGC | CGATGCTTGCCGTCCTTGTAATAC |
| NKA a | TGGTCGGAGTTTCATCAGAGACAC | CGAAGCAGAGTATGGAGCCTATCC |
| NKA b | ATCGCCAGAGAAATCACCCAC | CAATGAGAAAGACCACAGCCTCG |
| CA a | ACTATTCCCACCAATCGCCC | TGACGAGTTCTCCGTGTGTAACG |
| CA b | AGGGTGAAGAGTGTGTGGAGGAAG | AAAGACAGGATGCGGAGTTGCTGG |
| CA c | AGTCGTTGCTCGGTCACATACTG | TCGTAACACAGCGGGTTGTCTG |
| CA d | CAACTTTGGTGCCTTGACGC | TGAGATTCGGAGATGTAGAGGGG |
| CA x | TTCTGTCGGAGAGGGCTTGTGTTG | TACATCGGGTCGCTCACCATTC |
| CA e | ATCAGTTCCTCGCCCACTACTTC | TTCCTCCAGCAGCATCTTGC |
| CA f | AGTTACGAATCGGCGTTGGG | TCCACCTTCTTCTGCTGGTCAC |
| CA g | GGATAGAATGCTCTCACCCTACTGC | TGGCTGCGAAAGAACTAAGGC |
| CA h | CGCTTCTTCATCTTCAAGTGCC | TTGCTCATCTCCACCAACCG |
| HCN | TCCTGGACGAGATGAACGACTG | TACCACTCCGCTCACGAAAAGG |
| amt1/4 a | TGGCACTGGGAAGAAGACTGGAAC | GGAGAGGATGGTGTTGACGATG |
| amt-like | TGCTCATCACACAACACCGC | CCCGTCGTTCAATAGTTCCTCG |
| amt1/4 b | GACACCAGGGATTACGCATTTG | GGCAACCGAGTAGATGAGATACGAC |
| amt2/3 a | GCTGGTGGCAAATGGAAACTG | CGAGCACTCCGTTGATGATGTAG |
| amt2/3 b | CAAAGATGACCACCAACACTGAAG | GTGATGACAGCGAGCGAGTTATC |
| amt2/3 c | TGCCCTCGCAACATTCATACTC | AACGCTGTGCTCCATCAAGTCC |
| SLC1A | TGAAGCAGTCAGGATGGGAGTC | GTGAGGTAGAGGAGGAGGTCGC |
| SLC1B | GCAGCATCGCCGTCTGTTATTAC | TTTCCCATCACAACCCTGGAC |
| SLC1C | AGCACGAGGGACCGATGTATC | GGACCTTTCCGAAGACGACG |
| SLC5A | GCATTCGTCACTCACTCACCTTTAC | CCATCGCTGTTCCTCCTGC |
| SLC5B | ATGCTGGCGGTCAGTGTGG | AGGACGGCGGTGGAGTAGAC |
| SLC13 | CGTGAGGTTCAAGAAGGGGC | GAGAAGGAGGGCGTTTGGG |
| SLC4A | AGATGGGGGAGCGAGGACAG | TGGAAGGAGAGGGAGGAGACG |
| SLC4B | TCCAATGCGTCGGCAAAAG | TCCACGAAACATCTCAGCAGC |
| SLC4C | GCTTCAACTACAACCTCCACAACAC | GGCGAATCAACTGGGACGG |
| SLC8 | TTCACGGCAACCACTCTCTCC | TCCCAATGTGTAGACGAGCAGC |
| SLC9 | ATGGAGTGTTAGACCGAGAAATGG | ACGATGAAGGGCAGGGCGAC |
| SLC12A | TCAGAGGGAGGCAGGGACTC | GATGTTGAGCAGGCACCGTG |
| SLC12B | ACATCGGAGATTCATCCTCGC | CGGGTGTTGCCATTGTTGAG |
| SLC26A | TCCCAACACCGAAAGATGCTC | ACTGGAGATAGCGATGCCAATG |
| SLC26B | TCGCCTTGACGGAGACGAG | AGACAGCACGACCTCAGCCC |
| UBIQ | ACCCTCACTGGAAAAACCATCAC | TGTAATCAGACAGCGTTCGGC |
| 18S | TGAATCTGCCTGCTGATGAACC | GCTGATGTCACAACCAACCCAG |

*N.vectensis*

| GENE | QPCR PRIMER F | QPCR PRIMER R |
|---|---|---|
| Rhesus 1 | GGTCTCCTTCTTATCCTGTTCGTTG | CCGCTAAAACCATACTTCTTGAGG |
| Rhesus 2 | CGTTCTTACGAAAGCACGCCTAC | AAACTTGTCATTCCCGCCCTCG |
| Rhesus 3 | TCTTCTTCGCTTTCCCTTTCCTAC | GAATACTCCAGAGCAGATGATGGC |
| V-ATPase B | CAAAACAGTCTCTGGTGTCAATGG | CTTGGCGTCAATCCCTGATG |
| NKA a | TGCGAGATTCCTTTCAACTCTACC | TGTTCCTTGCCATTCACGAGG |
| NKA b | TCTTCTCAACCAACGCTGTGG | CAATGGCAATGGGAGTCTTTCC |
| CA 1 | TGGATTGTGTGTGCTTGGTGTC | TATTTCACTGAGGGGAGGCGTC |
| CA 2 | AAACTGGTTCCTGGCTTGTCATC | GACGAGTATTGGGTTCTTGAGCAC |
| CA 3 | CCTCAAGAACAATGGGCACG | TCCGTCTATCAGATGCTCCGAG |
| amt2/3 a | TGTCTTCGTGACGGGAATCTTG | TGGCGATACAACCTACTGGGTC |
| amt2/3 b | ATCGGGATTTGGACTGCTGG | ATGCGTTACTGCTCGGGCTATCTC |
| amt2/3 c | CGTCTTTGGGGCTGGAACATTC | ACCGTGCTTAGGGACATCAAGG |
| amt1/4 a | TAGACCAGTGTAGCCACCACCATC | AGAGTAGTGCGAGCAAAAAGAACC |
| amt2/3 d | TGAAGAGGGCGAGGACAAATC | TCGGTGACTCCAAAAGTGCTCC |
| amt1/4 b | CTCGTGCCTTCTGACTGGATTC | ATGAACAATACCGCTGCCCG |
| amt2/3 e | CGGAATGGTCTCCAGTAAGAACG | CCTCGGCATCGGTGAAAAAG |
| ATPsynthase | TGCTGGGAAAGTTCTGGACCAATG | ACACCCTCCTTGACGGTAACATTC |
| EF1b | TGCTGCA TCAGAACAGAAACCTGC | TAAGCCTTCAAGCGTTCTTGCCTG |

**Supplementary Table 5.** QPCR primers used in this study.
